# Synaptic targets and cellular sources of CB1 cannabinoid receptor and vesicular glutamate transporter-3 expressing nerve terminals in relation to GABAergic neurons in the human cerebral cortex

**DOI:** 10.1101/2024.10.14.618168

**Authors:** Peter Somogyi, Sawa Horie, Istvan Lukacs, Emily Hunter, Barbara Sarkany, Tim James Viney, James Livermore, Puneet Plaha, Richard Stacey, Olaf Ansorge, Salah El Mestikawy, Qianru Zhao

## Abstract

Cannabinoid receptor 1 (CB1) regulates synaptic transmission through presynaptic receptors in nerve terminals, and its physiological roles are of clinical relevance. The cellular sources and synaptic targets of CB1-expressing terminals in the human cerebral cortex are undefined. We demonstrate a variable laminar pattern of CB1-immunorective axons and electron microscopically show that CB1-positive GABAergic terminals make type-2 synapses innervating dendritic shafts (69%), dendritic spines (20%) and somata (11%) in neocortical layers 2-3. Of the CB1-immunopositive GABAergic terminals, 25% were vesicular-glutamate-transporter-3 (VGLUT3)-immunoreactive, suggesting GABAergic/glutamatergic co-transmission on dendritic shafts. *In vitro* recorded and labelled VGLUT3 or CB1-positive GABAergic interneurons expressed cholecystokinin, vasoactive-intestinal-polypeptide and calretinin, had diverse firing, axons and dendrites, and included rosehip, neurogliaform and basket cells, but not double bouquet or axo-axonic cells. CB1-positive interneurons innervated pyramidal cells and GABAergic interneurons. Most glutamatergic synaptic terminals formed type-1 synapses and some were positive for CB1 receptor concentrated in the presynaptic active zone, unlike in GABAergic terminals. From the sampled VGLUT3-positive terminals, 60% formed type-1 synapses with dendritic spines (80%) or shafts (20%) and 52% were also positive for VGLUT1, suggesting intracortical origin. Some VGLUT3-positive terminals were immunopositive for vesicular-monoamine-transporter-2, suggesting 5-HT/glutamate co-transmission. Overall, the results show that CB1 regulates GABA release mainly to dendritic shafts of both pyramidal cells and interneurons, and predict CB1-regulated co-release of GABA and glutamate from single cortical interneurons. We also demonstrate the co-existence of multiple vesicular glutamate transporters in a select population of terminals probably originating from cortical neurons and innervating dendritic spines in the human cerebral cortex.

## 1 INTRODUCTION

Throughout the brain, synaptic transmission and plasticity is regulated by presynaptic receptors expressed in glutamatergic and GABAergic cell types in the cerebral cortex (Bocchio et al., 2019; Hill et al., 2007; Katona et al., 1999; 2006). One of the most abundant presynaptic receptors is the G-protein-coupled CB1 cannabinoid receptor. It is activated by endocannabinoids, such as 2-arachidonoylglycerol (2-AG) and anandamide released by postsynaptic cortical neurons tonically or in an activity dependent manner (for review see Sugaya and Kano, 2022). Glial cells are also involved in endocannabinoid signaling (Ilyasov et al., 2018; Navarrete et al., 2014; Vincente-Acosta et al., 2022; Walter and Stella, 2003). Phasic depolarisation of postsynaptic cortical neurons evokes calcium entry through voltage-gated calcium channels leading to the release of 2-AG synthesized by diacylglycerol-lipase-alpha (DAGLα) (Hashimotodani et al., 2005; Tanimura et al., 2010). Furthermore, activation of G_q/11_ coupled metabotropic receptors, such as group-1 metabotropic glutamate receptors (mGluRs) or M1,3 muscarinic acetylcholine receptors also induce 2-AG release (Fukudome et al., 2004; Kim et al., 2002; Ohno-Shosaku et al., 2002). The postsynaptic synthetic machinery of 2-AG and the presynaptic CB1 receptors are tightly coupled in membrane microdomains (Barti et al., 2024; Dudok et al 2015). When activated, CB1 receptors reduce neurotransmitter release (Kreitzer and Regehr, 2001; Ohno-Shosaku et al., 2001; Wilson and Nicoll, 2001) by suppressing presynaptic Munc18-1 activity necessary for vesicle fusion (Schmitz et al., 2016). In addition, they inhibit voltage-gated calcium channels (Huang et al., 2001) resulting in short-term (STD) or long-term depression (LTD) of synaptic function, as well as may produce a tonic inhibition of transmitter release. The activation of CB1 receptors can also activate G-protein coupled potassium channels (Henry and Chavkin, 1995; Marinelli et al 2008).

Regulation of transmitter release by CB1 receptor activation has been extensively studied in GABAergic cortical neurons from rodents (Ali et al., 2007; Barti et al., 2024; Hajos et al., 2000; Hill et al., 2007; Losonczy et al., 2004; Neu et al., 2007; Trettel and Levine, 2002), but much less in humans (Chou et al., 2022; Kovacs et al., 2012; Ludanyi et al., 2011). These receptors are also the target of recreational and medical use of cannabis-derived substances and synthetic drugs. The drugs and endocannabinoids may also regulate key neurotransmitter receptors directly, such as the GABA-A (Golovko et al., 2014; Sigel et al., 2011) and other receptors (De Petrocellis et al. 2017; Fan, 1995; Morales and Reggio, 2017). Despite of the immense neurobiological significance of the cannabinoid signalling system in health and disease (Bernal-Chico et al., 2022; Foldy et al., 2013; Iversen 2003; Lowe et al., 2021; Lutz et al., 2015; Piomelli and Mabou-Tagne 2022), very little is known about the precise location of the molecular machinery of cannabinoid signalling in human cortical neurons and the cell types affected by it.

Endocannabinoid signalling affects both cortical glutamatergic and GABAergic neuronal synaptic transmission as well as other subcortical afferents. Cortical GABAergic neurons can be divided into PVALB/parvalbumin, SST/somatostatin, VIP/vasoactive intestinal polypeptide and LAMP5/PAX6-expressing families based on immunohistochemical and transcriptomic data (Bakken et al., 2021; Hodge et al., 2018; Krienen et al., 2020), which also correlate with electrophysiological signatures (Lee et al., 2023). The caudal ganglionic eminence derived CCK and/or VIP expressing groups express particularly high level of CB1 receptor in their axons in rodents and humans (Katona et al 1999; http://celltypes.brain-map.org/rnaseq/). The role of a given neuronal type, and in this case the action of endocannabinoids as well as extrinsic drugs acting on CB1 receptors, is determined by its synaptic connections. Quantitative synaptic output target identity evaluation of GABAergic cortical neurons shows that they are highly selective in the selection of both the postsynaptic cell types they innervate and in the placement of their synapses on the surface of postsynaptic neurons (Kawaguchi and Kubota, 1997; Somogyi et al., 1998). Some GABAergic neurons, the axo-axonic cells, only innervate the axon initial segments of pyramidal cells (Somogyi 1977) and act via GABA-A receptors (Buhl et al., 1993). Others, often referred to as basket cells, place varying proportion of their synapses on the somata and proximal dendrites (Szentagothai, 1977; Somogyi et al., 1983). The dendritic tree of postsynaptic neurons is subdivided by distinct types of dendrite-targeting GABAergic neurons (Lukacs et al., 2023; Tamas et al., 1997), mainly innervating the shafts and to a lesser extent dendritic spines of postsynaptic neurons. Furthermore, there are specialised interneurons, such as the double bouquet cell, which place a large proportion of their synaptic boutons on dendritic spines, each of which also receive a glutamatergic synapse (Kawaguchi et al., 1998; Lukacs et al., 2023; Somogyi and Cowey, 1981; Tamas et al., 1997). How such synaptic target selectivity is related to the presynaptic regulation of GABA release specifically in the human cortex is largely unknown.

The subcellular location of the CB1 expressing GABAergic terminals on the surface of postsynaptic neurons is key to understand the contribution of various GABAergic neurons to neuronal information processing. For example, CB1 expressing presynaptic GABAergic terminals on the soma or the axon initial segment and the accompanying endocannabinoid mechanisms governing network excitability through either phasic or tonic endocannabinoid signalling (Barti et al., 2024; Neu et al., 2007), is very likely to regulate the output of the whole neuron following the integration of active inputs. In contrast, CB1 expressing GABAergic terminals selectively targeting parts or the whole dendritic tree would contribute to the regulation of incoming domain-specific inputs (Bloss et al., 2016_;_ Boldog et al., 2018) active at the time of neuronal firing, back-propagating action potential evoked calcium entry and the accompanying endocannabinoid release (Hsieh and Levine, 2013). Such dendritically terminating CB1 expressing terminals are also likely to be influenced by local calcium signals in the postsynaptic dendritic shafts or spines (Larkum et al., 1999; Lovett-Barron et al., 2012). Furthermore, there is evidence in the rodent hippocampus that somatic, but not dendrite targeting GABAergic synapses are under tonic endocannabinoid suppression (Lee et al., 2015). Explaining the selective subcellular regulation of distinct aspects of synaptic plasticity requires knowledge of the circuits, the location of CB1 expressing terminals and the cell types that provide and receive them.

In the present study, we explored the location and postsynaptic targets of presynaptic terminals expressing CB1 receptors in human associational cortical areas. Throughout this paper we use the descriptive terms type-1 (often called asymmetrical) and type-2 (often called symmetrical) synapses (Gray, 1959). To determine the origins of CB1 positive GABAergic terminals, we recorded single cortical interneurons in vitro and visualised their axons for immunohistochemical testing for CB1 receptor as well as for molecular cell type markers, including the vesicular glutamate transporter type-3 (VGLUT3). Axonal characteristics allowed us to identify specific CB1-expressing cell types and relate them to transcriptomic cell types (Bakken et al., 2021; Hodge et al., 2019; Lee et al., 2023).

## 2 METHODS

### 2.1 Ethical approval and patient consent

Human tissue samples from neurosurgery at the John Radcliffe Hospital (Oxford) for the treatment of brain tumours or temporal lobe epilepsy (table 1) were collected in accordance with the Human tissue Act 2004 (UK), under the licence (15/SC/0639) of the Oxford Brain Bank (OBB), John Radcliffe Hospital, Oxford, UK. Fully informed patients consented to providing samples, which were access tissue that were removed in order to access the diseased part of the brain (Table 1).

**Table 1.**
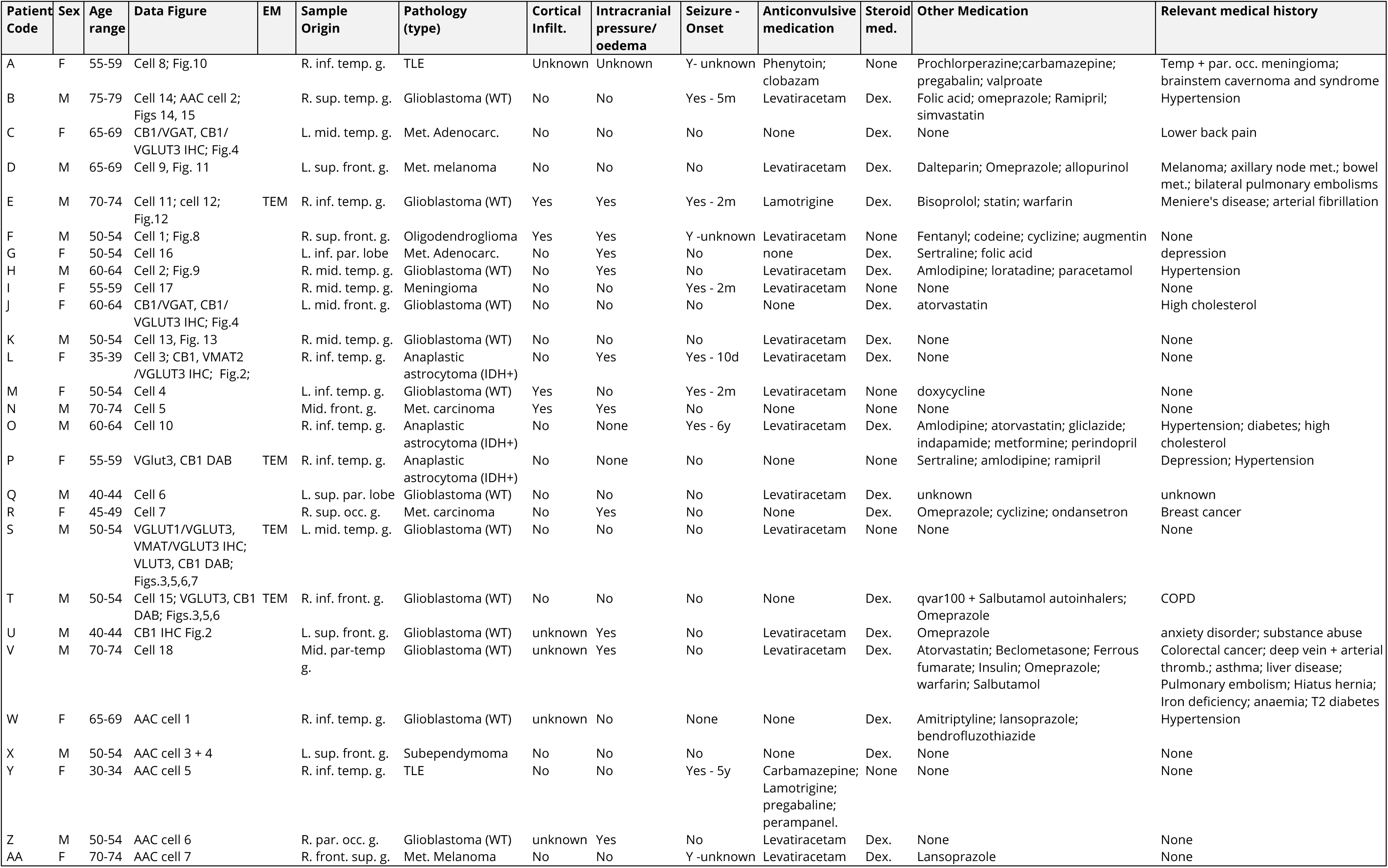
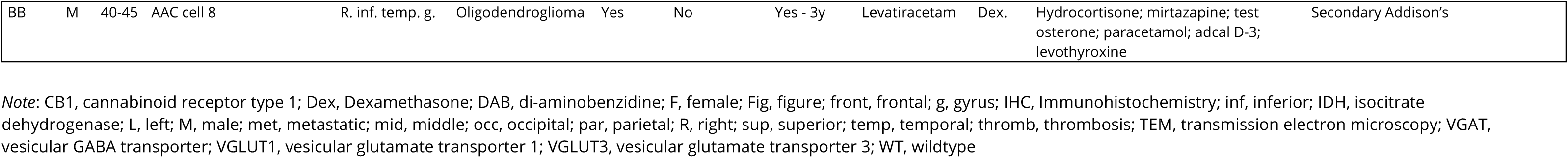
Patient data.

### 2.2 Sample collection and slice preparation

Sample collection, preparation and methods have been described (Bocchio et al., 2019; Field et al., 2021; Lukacs et al., 2023) and are summarised briefly. A small block of neocortex removed with a scalpel was immersed in ice-cold cutting artificial cerebrospinal fluid (ACSF) saturated with carbogen (95 % O_2_, 5 % CO_2_) and containing (in mM): 92 N-methyl-D-glucamine (NMDG), 2.5 KCl, 1.25 NaH2PO4, 30 NaHCO_3_, 20 4-(2-hydroxyethyl)-1-piperazineethanesulfonic acid (HEPES), 25 glucose, 2 thiourea, 5 Na-ascorbate, 3 Na-pyruvate, 0.5 CaCl_2_·4H2 O and 10 MgSO_4_·7H_2_O (pH ∼7.3, ∼300 mOsm/L). Slices of ∼350 µm thickness were prepared as described in Field et al. (2021), and the cutting ACSF was gradually replaced by storing ACSF, which contained the same components as the cutting ACSF except the NMDG was replaced with 92 mM NaCl. Slices were stored in storing ACSF at room temperature until recording. The recording ACSF contained the following (in mM): 130 NaCl, 3.5 KCl, 1.3 NaH_2_PO_4_, 24 NaHCO_3_, 3 CaCl_2_, 1.5 MgSO_4_, 12.5 glucose, (pH ∼7.3, ∼300 mOsm/L). All solutions were continuously bubbled with carbogen.

### 2.3 Electrophysiological recordings

Electrophysiological recordings were performed in recording ACSF saturated with carbogen perfused through the recording chamber at a flow rate of ∼10 ml/min over 10-16 hours after slicing. Glass capillaries (4-7 MΩ) were filled with an internal solution containing the following (in mM): 126 K-gluconate, 4 KCl, 4 ATP-Mg, 0.3 GTP-Na_2_, 10 Na_2_-phosphocreatine, 10 HEPES, 0.03 ethylene glycol-bis(2-aminoethylether)-N,N,N’,N’-tetraacetic acid (EGTA), and 0.05% (w/v) biocytin (pH ∼7.3, 280–290 mOsmol/L). Neurons were visualised by differential interference contrast (DIC) microscopy. Whole-cell patch-clamp recordings were performed from neurons in layers I-III, at 33-37 °C, using either an EPC-10 triple patch clamp amplifier and Patchmaster software (HEKA), or a Multiclamp 700B amplifier and pClamp software (Molecular Devices). Data was digitized at 100 kHz for current-clamp recordings (EPC-10 amplifier), or at 10 kHz in both recording modes (Multiclamp 700B). The reported voltage values are not compensated for a 16.5 mV junction potential. For each cell voltage responses to a series of 800 ms long current square pulses starting from holding current −100 pA until rheobase (RB) +100 pA with 20 pA increments between sweeps were recorded in current clamp mode (I-V traces). The initial holding current was between 0 and −100 pA, required to maintain the membrane voltage at ∼ −75 mV, and was aimed to be a multiple of 20 pA. Bridge balance was not adjusted during current clamp recordings. In paired recordings, a pair or a train of five action potentials (APs) were evoked at 50 ms intervals in one neuron by brief current injection, while the other neuron was current clamped such that its membrane potential was around −50 mV in order to detect evoked monosynaptic inhibitory postsynaptic potentials (eIPSPs). Uncompensated series resistance was monitored every minute by application of a 10 ms voltage step of - 10 mV. Action potentials and ionotropic glutamate receptors were not blocked.

### 2.4 Data analysis and inclusion criteria

The following criteria were applied for inclusion of the cells’ I-V traces in the analysis: 1. The holding current was between 0 and −100 pA for holding the cell at −75 mV; 2. At least one overshooting AP could be elicited by depolarising current injections; 3. Fast pipette capacitance was successfully compensated and no oscillation artefacts were observed in current clamp mode.

PatchMaster (.dat) files were opened in Igor Pro software v7.0.8.1 (WaveMetrics) using Patchers’s Power Tools (Department of Membrane Biophysics, Max Planck Institute for Biophysical Chemistry, Göttingen, Germany, http://www3.mpibpc.mpg.de/groups/neher/index.php?page=aboutppt).

After digitally adjusting the I-V traces to remove the artefact resulting from the bridge balance the following parameters were measured: resting membrane potential (Vm), was measured as the steady state voltage in response to 0 pA current injection. The membrane time constant (*τ*) and whole-cell capacitance (C_m_) were calculated from the double exponential fit to the membrane voltage in response to a −100 pA current step in current-clamp recordings as described in Golowasch et al. (2009) and reported (Lukacs et al., 2023). The sag ratio was calculated as the ratio of the maximum voltage deflection over the difference of the steady state voltage and the baseline voltage, in response to holding current −100 pA current injection. The rheobase was measured as the current injected when the first AP was generated; single AP kinetic parameters were measured on the first AP elicited at RB current injection in Matlab R2020a (MathWorks), using a custom written script as described previously (Field et al., 2021).

### 2.4 Visualisation of recorded neurons and immunohistochemistry

After completion of the recording and filling of the recorded cells with biocytin (at least 5 min), the slices were immersed in a fixative of 4% (w/v) paraformaldehyde and 15% (v/v) saturated picric acid in 0.1 M PB at pH ∼7.2 at 4°C overnight. For some samples the fixative also contained 0.05% (w/v) glutaraldehyde. Slices were re-sectioned into 3-5 ∼60 µm thick sections with a vibratome. Two sections, including the one in which the soma was predicted to be located, were incubated in Alexa 488-conjugated streptavidin (1:1000, Invitrogen) or Cy3 conjugated streptavidin (1:400, Jackson) in PB for the visualisation biocytin.

Immunohistochemical reactions were performed using up to nine different primary antibodies on different sections and produced in different host species. For a list of all primary abs used in this study see Table 2. Secondary abs (donkey, Jackson Immuno Research) against immunoglobulins of the host species of the primary ab, conjugated to different fluorophores, were added at appropriate dilutions (Alexa405/DyLight4/brilliant violet421 (blue)- and cyanine5 (Cy5)/DyLight647 (infra-red)-conjugated abs in 1:250; Alexa488 (green)-conjugated abs in 1:1000; Cy3 (red)-conjugated abs in 1:400).

**TABLE 2.** Primary antibody information.

| Antibodies to molecule | host species | antigen | clonality | lab reference | dilution | stock protein conc. (ug/ml) | source | product code | specificity reference |
| --- | --- | --- | --- | --- | --- | --- | --- | --- | --- |
| calbindin (CB) | rabbit | rec. rat CB D-28k | poly | 989 | 1:5000 | antiserum | SWANT | CB-38 | Airaksinen et al., 1997 |
| calbindin (CB) | goat | mouse CB D-28k expressed in bacteria | poly | 1427 | 1:500 | 200 | Frontier Institute | Af1040 | Nakagawa et al., 1998 |
| cannabinoid type 1 receptor (CB1) | guinea pig | mouse CB1, C-terminal 31 aa | poly | 1274 | 1:5000 | 200 | Frontier Institute | Af530 | Fukudome et al., 2004 |
| cannabinoid type 1 receptor (CB1) | rabbit | synthetic peptide aa residues 443-473, mouse and human | poly | 1558 | 1:10000 | 10000 | Immunogenes Ltd. | Anti-CB1 | Dudok et al., 2015 |
| cholecystokinin 8 (CCK-8) | guinea pig | cysteine tagged CCK-8 |  | 1306 | 1:500 | 380 | Masahiko Watanabe | n/a | Dot blot |
| calretinin (CR) | goat | human rec. CR | poly | 1116 | 1:1000 | antiserum | SWANT | CG1 | Schiffmann et al., 1999 |
| chicken ovalbumin upstream promoter transcription factor II (COUP-TFII) | mouse | rec. human COUP-TFII/ NR2F2 aa 43-64 | mono | 1096 | 1:250 | 1000 | Perseus proteomics | PP-H7147-00 | Lee et al., 2004 |
| Glutamic acid decarboxylase 67 (GAD67) | mouse | rec. GAD67 protein for 67 kDa isoform | mono | 1223 | 1:500 | 1000 | Chemicon | MAB5406 | Chemicon western blot - cell lysate control |
| hexaribonucleotide binding protein-3 (NeuN) | mouse | NeuN protein | mono | 842 | 1:1000 | 1000 | Chemicon | MAB377 | Nuclear labelling pattern |
| parvalbumin (PV) | guinea pig | rat rec. protein aa 1-110 | poly | 1310 | 1:5000 | antiserum | Synaptic Systems | 195 004 | Synaptic Systems |
| PV | goat | rat muscle PV | poly | 1258 | 1:1000 | antiserum | SWANT | PVG-213 | Schwaller et al., 1999 |
| PV | rabbit | rec. rat PV | poly | 1527 | 1:5000 | antiserum | SWANT | PV 27 | Schwaller et al., 1999 |
| pro-CCK | rabbit | c-terminal -9aa pro CCK affinity purified | poly | 1090 | 1:500 | antiserum | Frontier Institute NITTOBO MED CO. | Af350 | Booker et al., 2017 |
| somatostatin (SM) | sheep | SM conj. to carrier protein | poly | 1085 | 1:500 | 5000 | Cortex Biochem | CR2056SP | Immunolabel as others |
| somatostatin (SM) | mouse | SM-14 conj. to carrier protein | mono | 1276 | 1:400 | 140 | GeneTex | GTX71935 | Immunolabel as others |
| vesicular GABA transporter (VGAT) | guinea pig | rat rec. protein aa 2-115 | poly | 1321 | 1:1000 | antiserum | Synaptic Systems | 131 004 | Sy Sy KO verified |
| vasoactive intestinal polypeptide (VIP) | rabbit | porcine VIP coupled to bovine thyroglobulin with carbodiimide linker | poly | 1523 | 1:5000 | antiserum | Immunostar | 20077 | Immunostar |
| vasoactive intestinal polypeptide (VIP) | mouse | VIP | mono | 1053 | 1:25000 | antiserum | CURE/DDRC/RIA Core-NIH #DK41301 | 55 | Wong et al., 2009 |
| vesicular glutamate transporter 1 (VGLUT1) | guinea pig | rec. protein for residues near carboxy terminus of rat VGLUT1 | poly | 1579 | 1:500 | antiserum | Synaptic Systems | 135304 | SySy KO verified |
| vesicular glutamate transporter 3 (VGLUT3) | rabbit | human VGlut3 affinity purified | poly | 1559 | 1:10000 | antiserum | Salah El Mestikawy | n/a | Vigneault et al., 2015 |
| vesicular mono-amine transport 2 (VMAT2) | rabbit | rat VMAT2 468-515aa | poly | 1014 | 1:50,000 | 200 | Frontier institute | Af720 | Frontier Institute Immunoblot |

Note: aa, amino acid; rec, recombinant; conc, concentration; conj, conjugated; mono, monoclonal; poly, polyclonal;

### 2.5 Wide field epifluorescence and confocal laser scanning microscopy

Cells visualised with Alexa 488- or Cy3-conjugated streptavidin and fluorescent immunoreactions were first evaluated in a wide field epifluorescence microscope and recorded with a digital camera controlled by OpenLab software (Improvision). The light source was either a pE-300 LED lamp (CoolLED) or a mercury arc lamp (HBO, Osram), for which the light was spectrally separated by dichroic mirrors to obtain optimal excitation and emission bandwidths for each fluorophore. For higher resolution imaging, confocal laser scanning microscopy was performed using an LSM 710 axioImager.Z1 microscope (Zeiss) and DIC M27 Plan-Apochromat 40x/1.3, 63x/1.4 and alpha Plan-Apochromat 100x/1.46 oil immersion objective lenses, controlled by ZEN 2008 software (v 5, Zeiss), as described previously in detail (Lasztoczi et al., 2011). Signal from each fluorophore was recorded in separate scanning tracks and channels, using the following lasers: for Alexa405, DyLight405 and brilliant violet421, a 405 nm solid state laser; for Alexa488 a 488 nm argon laser; for Cy3 a 543 nm He-Ne laser; for Cy5 and DyLight647 a 633 He-Ne laser. Pinhole size was adjusted optimally for similar optical slice thickness between tracks. The step size along the Z imaging axis was set to half of the thickness of the optical slices, as per the Nyquist criterion for optimal sampling. Details of the optical slice thickness are given in the figure descriptions.

### 2.6 Quantification of CB1/VGLUT3 co-expression in nerve terminals

A total of 10 laser scanning confocal Z stacks were acquired from two cases (patients C, left mid. temp. gyrus; and J, left mid. front. gyrus; Table 1, 5 stacks each) with an LSM 710 axioImager.Z1 microscope (Zeiss) and DIC M27 Plan-Apochromat 40x/1.3 oil immersion objective lens, controlled by ZEN software (v14, Zeiss). Immunoreactivity for CB1 was visualised using a guinea pig antiserum and an Alexafluor488 conjugated secondary antibody and a 488nm argon laser; immunoreactivity for VGLUT3 was visualised using a rabbit antibody (Vigneault et al., 2015) and a Cy3-conjugated secondary antibody detected with a 543 nm He-Ne laser. Z stacks were made up of 16 or 24 optical slices, with a total depth of 10 μm. A single 30×30μm grid square was placed randomly within each stack in ZEN (v16, Zeiss) and all fluorescence dots within a size range of nerve terminals were counted and then indexed as either CB1 immunopositive (CB1+), VGLUT3 immunopositive (VGLUT3+) or both. The size range was estimated from boutons along continuously immunoreactive axons for CB1. Dots were also checked for potential non-immunoreactive signal, e.g. autofluorescence of lipofuscin, in a third recorded ‘empty’ channel, visualised with a 405nm solid state laser. Most dots were followed individually through more than one optical slice to ensure consistent signal overlap, or the lack of it, in a given dot. Some of the VGLUT3-positive dots were clearly negative for CB1 and conversely, indicating no cross-talk between channels.

### 2.7 Quantification of VGLUT3/VGLUT1 co-expression in nerve terminals

A total of 6 Z -stacks were taken from 2 samples with a 63x/1.3 oil immersion objective lens, 3 stacks from each (patient L, right inf. temp. gyrus; and S, left mid. temp. gyrus; Table 1). Immunoreactivity for VGLUT3 was visualised by Alexa488 and for VGLUT1 using a guinea pig antibody (Synaptic Systems, table 2) with a Cy3-conjugated secondary antibody. Stacks were of variable depth due to variability in the penetration of antibodies into the thick sections. Between 11 and 15 optical slices were taken in each Z stack with a total depth of 4.9 - 5.6 μm and optical slice thickness set to optimal as per the software. Within each stack two 50×50μm grid squares were placed at random in ZEN (v16) and all VGLUT3 immunoreactive dots/boutons within these squares in the size range of nerve terminals were counted (356 boutons in total). Each of these boutons was then tested for the presence of VGLUT1 immunoreactivity in multiple optical slices. Most VGLUT1-positive boutons were clearly VGLUT3-neagative and conversely many VGLUT3-positive dots were clearly negative for VGLUT1, indicating that no cross-talk was present between the channels.

### 2.8 Detection of VMAT2 and VGLUT3 in nerve terminals

Reactions were carried out in sections from two patients (patient L, right inf. temp. gyrus; and S, left mid. temp. gyrus; Table 1). Both primary antibodies were produced in rabbits, therefore to distinguish the two fluorescence signals in two different channels, we applied a very low concentration of the rabbit polyclonal antibody to VMAT2 (1:50,000 in TBS-tx, 1% NGS, 3 nights) and used tyramide signal amplification (Tyramide Super Boost kit from Invitrogen). After blocking non-specific binding with 10% goat serum and then any endogenous peroxidase activity with 3% H_2_0_2_, before washing in TBS-tx, the sections were incubated in goat anti-rabbit HRP-conjugated secondary antibody as per manufacturer’s protocol. After 4 hours sections were washed and incubated for 30 mins in Alexa Fluor 647 tyramide reagent (1:100 in TBS, 0.05% H_2_0_2_). Reaction stop mixture was applied and sections were washed again before proceeding with a second immunoreaction in TBS-tx to visualise VGLUT3 immunoreactivity with polyclonal rabbit antibody, as above, and Alexafluor488 conjugated secondary antibody. Controls included omitting both the tyramide amplification step and the primary antibody to VGLUT3, and using only the Alexa488 conjugated secondary anti-rabbit antibody. We could not detect VMAT2 immunoreactivity in these control sections.

### 2.9 Peroxidase reactions of recorded and biocytin labelled neurons

Some or all sections from slices containing recorded and labelled cells were converted by avidin-biotin horseradish peroxidase (HRP) reaction with 3,3’-diaminobenzidine (DAB) as chromogen for the visualisation of biocytin, and embedded in epoxy resin (Sigma) for light microscopic examination and reconstruction. Following fluorescence visualisation and evaluation of biocytin labelled cells, sections were incubated in biotinylated peroxidase complex (B) 1:100 v/v (Vectastatin ABC elite kit, Vector Laboratories) in TBS-Tx for 4 hrs at room temperature, then incubated in avidin plus B (A+B) 1:100 v/v in TBS-Tx over 36 hrs at 4°C. Next, sections were pre-incubated in 0.5 mg/ml DAB in Tris Buffer without saline (TB) for 10 minutes in dark. Subsequently, 0.002% w/v H_2_O_2_ substrate was added to initiate oxidation and precipitation of DAB. Depending on the intensity of the labelling, reactions were stopped after several minutes and sections were washed 4×10 minutes in 0.1M PB. For contrast enhancement, sections were incubated in 0.5% w/v Osmium tetroxide solution in 0.1M PB for 1 hour at room temperature. Before mounting, sections were dehydrated using increasing concentrations of ethanol and a final step of propylene oxide (Sigma). From propylene oxide, sections were quickly transferred into Durcupan epoxy resin (Sigma), left for several hours and mounted on glass.

### 2.10 Light microscopy and reconstruction of labelled cells

Transmitted light microscopic analysis of HRP visualised neurons was used to identify cell types. Some neurons were reconstructed to demonstrate the distribution of their dendrites and axon across different neocortical layers, as well as for the identification of their axonal boutons. Neurons were manually traced using a drawing tube attached to a transmitted light microscope (Leitz Dialux22, Leica), equipped with a Pl Apo 63x/1.4 or 100x oil immersion objectives. After alignment of the tracings of consecutive sections, drawings were overlaid and copied onto a single sheet of paper and digitised.

### 2.11 Peroxidase-based immunohistochemistry and serial section transmission electron microscopy of CB1 and VGLUT3 immunolabelling

For electron microscopic peroxidase immunohistochemistry, 70 µm thick sections were cut from small cortical blocks which were fixed within 3-5 min after removal from the brain using a fixative of 4% (w/v) paraformaldehyde, 15% (v/v) saturated picric acid containing 0.05% (w/v) and glutaraldehyde in 0.1 M PB at pH ∼7.2 at 4°C for several hours. The sections were incubated in 20% sucrose in 0.1M PB for 2 hrs at room temperature for cryoprotection, followed by quickly freezing the sections using liquid nitrogen and thawing them in PB with sucrose once or twice. Guinea pig primary antibody to CB1 (1:10,000, Table 2) or rabbit antibody to VGLUT3 (1:10,000, Table 2) diluted in 1% NGS PB were applied for one day at room temperature, followed by appropriate biotinylated goat secondary antibodies (1:100, Vector Labs) for 5 hours and ABC (Vectastain ABC elite kit, Vector Labs) diluted in 0.1 M PB overnight. All following steps were the same as for HRP reactions on triton-treated sections with the exception that the buffer did not contain detergent. During dehydration, these sections were treated with 1% (w/v) uranyl acetate dissolved in 70% ethanol for 40 min in dark for further enhancement of contrast for electron-microscopy.

Electron-microscopic sections (50-70 nm) were cut using a diamond knife and mounted on pioloform-coated copper slot (2×1 mm) grids and studied at 80 keV accelerating voltage and recorded digitally using GATAN software. Sections were contrasted with lead citrate. The labelled boutons and their synaptic targets were imaged in 2-10 serial electron microscopic sections to identify the synaptic junction and to ensure the differentiation of postsynaptic dendritic shafts and dendritic spines.

### 2.12 Statistics

Statistical tests were carried out in R (The R Project for Statistical Computing) and are reported in full in the results. Descriptive statistics are as mean ± standard error, unless stated otherwise in the text. For testing associations between different synapse types and their synaptic targets and for associations between cortical areas and synapse types, Fisher’s exact test was used.

## 3 RESULTS

### 3.1 Transcriptomic pattern of CB1 and VGLUT3 expression in the human cerebral cortex

To address the cellular and subcellular distribution of CB1 receptor expression we first examined the transcriptomic profiles of cortical neurons in two databases http://celltypes.brain-map.org/rnaseq/. The expression of CNR1 (for CB1) showed a wide distribution of both GAD1-expressing GABAergic and SLC17A7-expressing glutamatergic neurons in the data sets; one from post-mortem tissue samples taken from human primary motor cortex and processed using the 10x Genomics RNA-seq methodology (Figure 1a), the second (Figure 1b) profiled using SMART-seq v4 covering multiple cortical areas (middle temporal gyrus, anterior cingulate cortex, primary visual cortex, primary motor cortex, primary somatosensory cortex, and primary auditory cortex) and the third from the middle temporal gyrus (MTG Figure 1c), using SMART-seq v4 processed data (2018 set), as untrimmed means (Hodge et al., 2019; Bakken et al., 2020; http://celltypes.brain-map.org/rnaseq/; BRAIN Initiative Cell Census Network 2020). Amongst the neurons from multiple areas, 35/54 GABAergic cell groups and 21/67 glutamatergic cell groups showed appreciable levels of CNR1 expression. Likewise, amongst the neurons from the M1 area 46/72 of GABAergic and 15/46 glutamatergic neuron groups showed significant level of CNR1 expression. Amongst the GABAergic neurons, most CALB2 (calretinin) and/or CCK and/or VIP expressing neurons showed high levels of CNR1 expression with few exceptions (Figure 1). In the samples from multiple cortical areas, which include those studied here, at least 14 interneuron groups had CALB2/CCK/VIP and CNR1 co-expression, an additional 5 only CALB2/VIP/CNR1 or only CCK/CNR1 (5 groups) or only CCK/VIP/CNR1, or CALB2/CNR1 (one group each). There were also 7 groups of CCK expressing GABAergic cells without significant CNR1 expression. The LAMP5 expressing subclass mainly includes groups of neurogliaform cells, two of which expresses significant levels of CNR1. Similar trends are found also in the samples from M1 (Fig.1).

**FIGURE 1.**
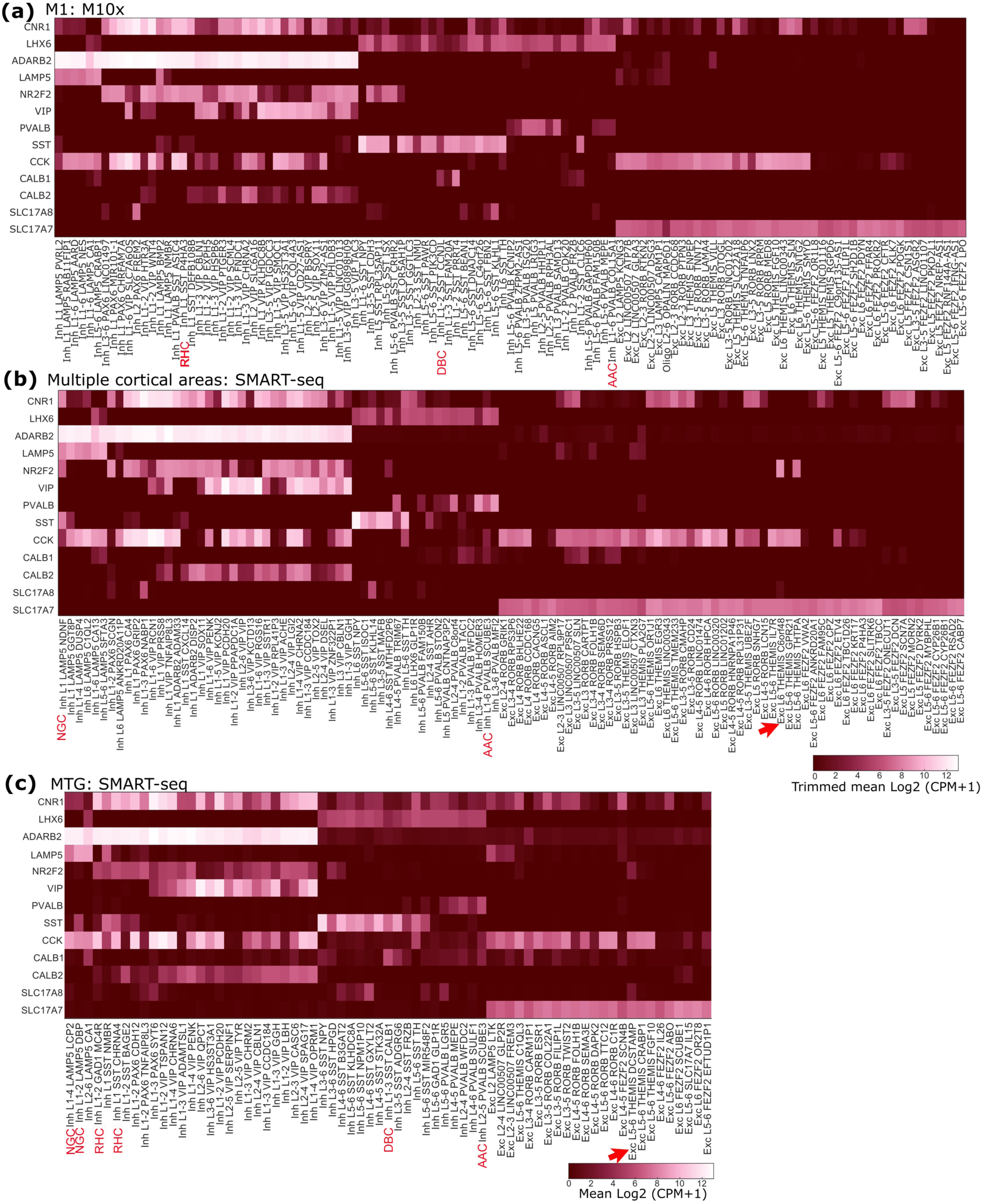
Analysis of the differential expression of CNR1 and SLC17A8, associated with some GABAergic or glutamatergic neuronal cell type markers in the Allen Institute human cortical transcriptomic data sets. Heatmaps of mRNA expression in three transcriptomic data sets **(a-c)** available at http://celltypes.brain-map.org/rnaseq/. (**a)** Data obtained from post-mortem tissue samples taken from human primary motor cortex and processed using the 10x Genomics RNA-seq methodology; trimmed means are shown. **(b)** Data set covering multiple cortical areas (middle temporal gyrus, anterior cingulate cortex, primary visual cortex, primary motor cortex, primary somatosensory cortex, and primary auditory cortex), profiled using SMART-seq v4; trimmed means are shown. **(c)** Middle temporal gyrus shows SMART-seq v4 processed data (2018 set), as untrimmed means. Note that the datasets differ in their cluster identifiers. Many GABAergic neuronal groups express CNR1, which strongly, but not completely, overlaps with the expression of ADARB2, NR2F2, VIP, CCK and CALB2, but infrequently with the expression of SST and PVALB. Very few GABAergic neuronal groups express SLC17A8 (for VGLUT3). Some easily identifiable GABAergic cell types such as rosehip cells (RH), double bouquet cells (DBC) and axo-axonic cells (AAC) are indicated. A number of SLC17A7 expressing (glutamatergic) neuronal types frequently express CNR1, but only one group Exc L6 THEMIS C6orf48 (arrow in b) has some cells also expressing SLC17A8. In the middle temporal gyrus, the group Exc L5-6 THEMIS DCSTAMP (arrow in c) has several cells expressing SLC17A8 as well as SLC17A7.

GAD1 expressing cells are GABAergic, but a distinct subpopulation also expresses SLC17A8 encoding VGLUT3, which in rodents also leads to the synaptic release of glutamate along with GABA (Pelkey et al., 2020). Therefore, we analysed SLC17A8 expression and found that in samples from multiple cortical areas 5 groups of interneurons showed some level of expression, two of which appeared to be stronger (Fig 1a). Similar, 5 groups of GAD1 expressing interneuron showed some SLC17A8 expression in M1. In samples from multiple areas, one SLC17A8 expressing group also expressed CCK, which is co-expressed with VGLUT3 in rodent cortical GABAergic neurons. In addition, one SLC17A8 weakly expressing group expressed SST and one PVALB. The fifth SLC17A8-expressing group, Inh L1-2PAX6 SCGN only expressed the common gene ADARB2 from those that we surveyed. Amongst the SLC17A7 (for VGLUT1) glutamatergic neuronal groups only one group, Exc L6 THEMIS CC6orf48 showed traces of SLC17A8 expression (Figure 1b,c) and none was found in the samples from M1. (Figure 1a).

The other CNR1 expressing GABAergic groups included SST expressing interneurons 4/11 (multiple areas) and 8/21 groups in M1. In contrast, amongst 10 PVALB expressing interneuron groups only 2 in multiple areas and none in M1 showed any CNR1 expression. The axo-axonic cell, identified as Inh L1-6 PVALB SCUBE3 was one notable PVALB expressing interneuron type showing low level of CNR1 (Fig 1a), in contrast to our observations by immunohistochemistry. Amongst the 5 CALB1 (for calbindin) expressing interneuron groups in data from multiple areas and 4 from M1 only one group in each expressed low levels of CNR1.

The above brief analysis suggests that the CB1 receptor is selectively expressed at high levels mainly in some of the ADARB2/VIP/CCK/CALB2 GABAergic neuronal groups derived from the caudal ganglionic eminence and rarely in others, such as PVALB expressing neurons (but see Figure 1c, MTG for PVALB/CNR1 co-expression). This raises the question of how the synaptic output of CB1-expressing and non-expressing neuronal families differs, one requiring CB1 receptor regulations whereas the other does not. We have analysed the synaptic placement of CB1-expressing terminals and recorded some of the CB1 expressing neurons listed above also testing their group identity by immunohistochemistry for cell type markers (see below). Similar questions arise for the glutamatergic neuronal groups, but their analysis is beyond the scope of this study.

### 3.2 Distribution of CB1-positive axons in frontal and temporal neocortex

Immunofluorescence visualisation of CB1-positive axons showed a rich network in all layers of both the superior frontal and inferior temporal gyri (Figure 2). In both areas and all layers, there was a wide range in the intensity of immunoreactivity in individual axons and their boutons. The reverse contrast fluorescent illustration (Figure 2) mainly shows the strongly immunopositive axons with highest density in layers 3 and 4. Layer 1 was particularly rich in CB1-positive axons, there was an increased density in layer 4 and much lower density in layers 5 and 6. In lower layer 3 to layer 5 some of the axons formed bundles aligned radially amongst the columns of neurons (Figures 2e,f). The intensity of reaction was uniform along individual axons indicating that the density of CB1 is characteristic of individual cells, and probably cell types. Although there were local clusters of boutons, these were not located around neuronal cell bodies. Basket-like formations of CB1 positive boutons around neurons, as they appear e.g. in rodent hippocampus (Katona et al., 1999) or in the monkey neocortex (Eggan et al., 2010), were rarely observed in the areas and layers studied. In addition to the continuously visualised axons, and their boutons, there were scattered immunopositive dots that could be detected at high magnification (not shown). Subsequent electron microscopic examination revealed were synaptic boutons, but without their connecting preterminal axons showing immunoreactivity (see below). To clarify the identity and network role of the various CB1-positive axon terminals we carried out electron microscopic analysis in layers 2-3 of several cortical areas.

**FIGURE 2.**
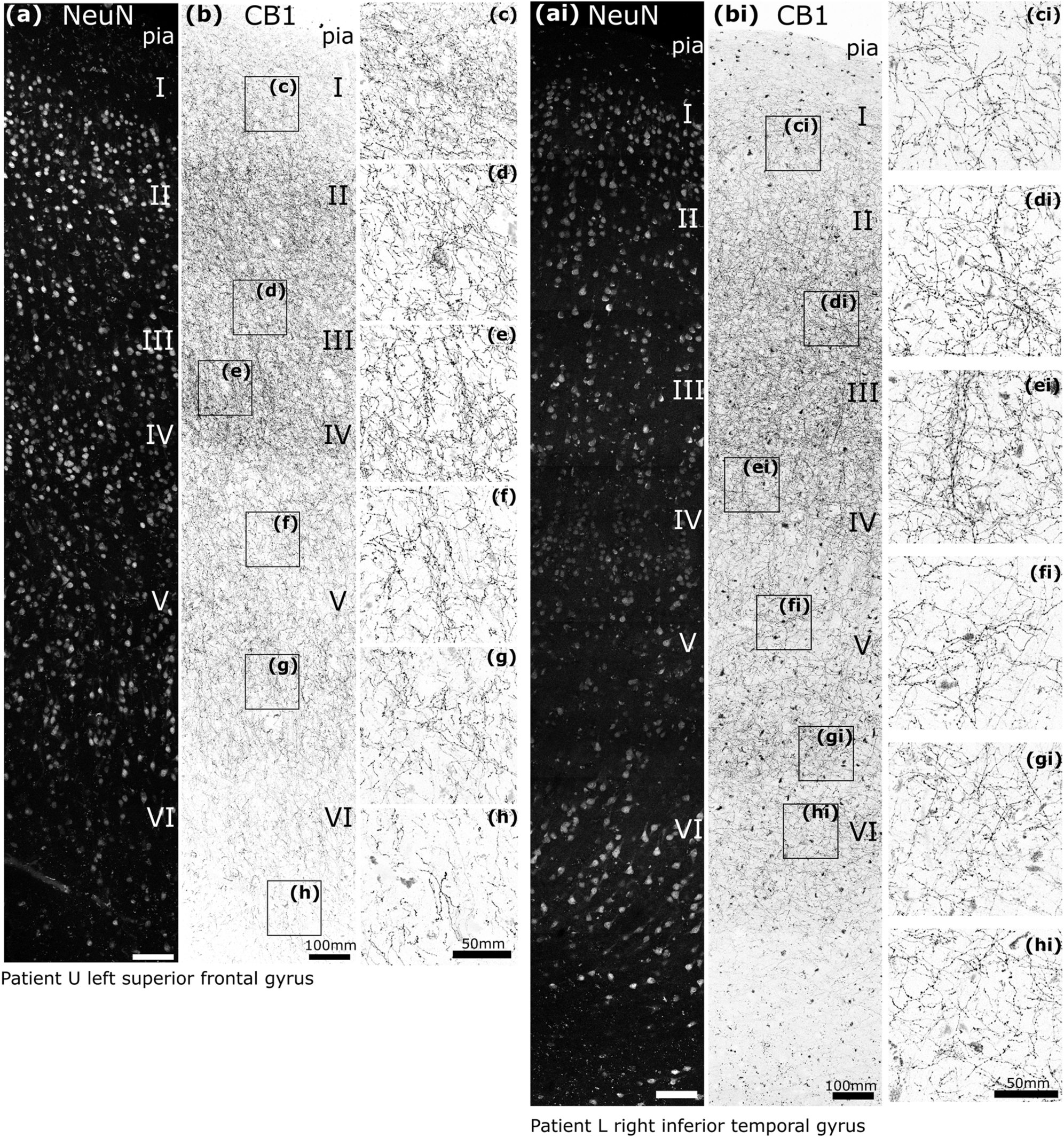
Distribution of strongly CB1 receptor immunoreactive axons and boutons in the human cerebral cortex. **(a, ai)** Confocal microscopic immunofluorescence images showing NeuN-positive neurons in layers I-VI in two cortical areas. **(b, bi)** Inverse contrast, immunofluorescence laser scanning images of the same cortical areas demonstrating different densities of strongly CB1-positive axons; framed areas are shown at higher magnification in **(c-h, ci-hi).** In some layers radial axon bundles are apparent. Z stacks of optical slices total thickness (a, b, 16 μm; ai, 8 μm; bi 6.7 μm), (c, ci – h, hi, 6-10 μm). Scale bars: (a, ai, b, bi) 100 μm; (c - h, ci – hi) 50 μm.

### 3.3 Synaptic targets and origin of CB1-immunoreactive boutons

The qualitative assessment of CB1-immunoreactivity revealed a very heterogeneous intensity of immunoreaction in individual axons and bouton-like dots suggesting a contribution from multiple sources and cell types in the human cortex. Indeed, from animal studies a multitude of intracortical (Bodor et al., 2005; Chou et al., 2022; Eggan and Lewis 2007; Eggan et al., 2010; Omiya et al., 2015) and subcortical (e.g. Nyiri et al., 2005) sources are expected to contribute to cortical CB1 receptor content.

First, we sought to establish the identity and targets of CB1-immunoreactive synaptic terminals. Electron microscopic analysis of immunolabelling of nerve terminals in layers 2-3 in three patients (P, S, T; Table 1) showed two distinct types of synaptic junctions made by immunopositive boutons (Figure 3). Strongly CB1-immunopositive boutons made type-2 junctions with small, or barely detectable postsynaptic membrane specialisation (Figure 3a-e). In contrast, some weakly immunopositive boutons established type-1 synapses with extensive postsynaptic membrane specialisations (Figure 3f,g). Glutamatergic terminals mostly make type-1 synapses and are erroneously called ‘excitatory’ although the released glutamate has many inhibitory actions, e.g. through presynaptic metabotropic receptors in the human cortex (e.g. Bocchio et al., 2019). Most GABAergic boutons make type-2 synapses, which are often called ‘inhibitory’ although GABA can be excitatory both directly and indirectly, e.g. reducing GABA release through presynaptic GABA_B_ receptors.

**FIGURE 3.**
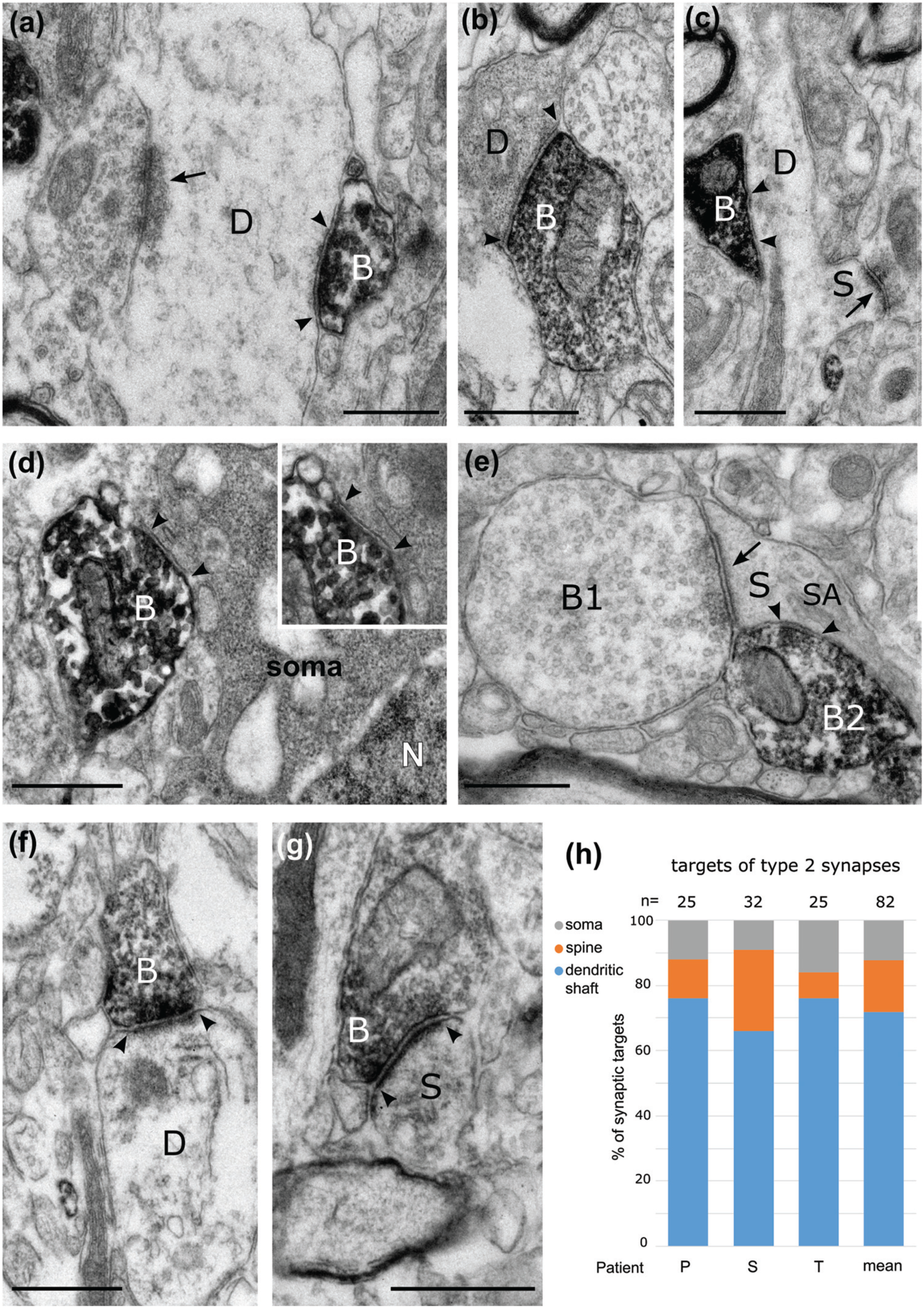
Electron micrographs of the synaptic targets and quantitative distribution of CB1 cannabinoid receptor expressing nerve terminals in layers 2-3 of the human cerebral cortex. Immunoperoxidase reactions show CB1-positive terminal boutons (B) filled with electron-opaque DAB-deposit (black); the synaptic junctions are marked by arrowheads. **(a)-(c).** Dendritic shafts (D) receive type-2 synapses from CB1-positive boutons. In *(a)*, a large dendrite also receives a type-1 synapse (arrow) from an unlabelled bouton. In *(c)*, the dendrite emits a sessile spine (S), which receives a type-1 synapse. **(d).** A CB1-positive bouton innervates a neuronal soma (N, nucleus); inset, the same synapse in a serial section. **(e)** A dendritic spine (S) with spine apparatus (SA) receives a type-1 synapse from a large bouton (B1) and a type-2 synapse from a CB1-positive bouton (B2). **(f, g)** CB1-positive boutons making type-1 synapses innervate a dendritic shaft (D) and a dendritic spine (S). The HRP reaction-product is concentrated at the presynaptic active zones. **(h)**. Distribution of the synaptic targets of CB1-positive boutons making type-2 synapses in three patients (P,S,T, and average); the majority of synaptic targets are dendritic shafts. Images (a, c, f, g) patient T, right inferior frontal gyrus; images (b, d) patient S, left middle temporal gyrus; image (e), patient P, right inferior temporal gyrus. Scale bars: 0.5 μm.

We have analysed 82 CB1-immunopositive boutons making type-2 synapses, a similar number from each of the three patients (Figure 3h). Most, but not all, of these boutons showed strong immunoreactivity that continued into the preterminal axons; both the boutons and axons had a uniform distribution of the cytoplasmic DAB reaction end-product. The antibody, raised in guinea pig, recognises epitope(s) on the intracellular side of the plasma membrane, hence the diffusible end-product fills the axons and boutons. A smaller number of boutons had weaker immunoreactivity that was also uniformly distributed along the plasma membrane. On average, the postsynaptic targets were mainly dendritic shafts (72 ± 3 %, patient P, *n*=25, 76 %; S, *n*=32, 66 %; T, *n*=25, 76%) and to a lesser extent dendritic spines (16 ± 5 %) and somata (12 ± 2 %). When they could be followed in serial section, postsynaptic dendritic spines always received an additional type-1 synapse from a CB1-immunonegative bouton (Fig. 3e). Some of the postsynaptic dendritic shafts emitted spines (Fig. 3c) suggesting that they originated from pyramidal cells, others received conspicuous type-1 synapses from CB1 immunonegative boutons (Fig. 3a), indicating an origin from interneurons. These observations are in agreement with the distribution of type-2 CB1-immunopositive synaptic targets in the monkey prefrontal cortex (Eggan et al., 2010).

Boutons immunopositive for CB1 receptor and making type-1 synapses were less frequently identified due to the weak reaction product, which was mainly localised to the presynaptic active zone and did not fill the entire bouton (Figure 3f,g). A total of 11 boutons pooled from two patients (patient S, n=3; T, n=8) made type-1 synapses with both dendritic shafts (*n*=5) and spines (*n*=6). Due to potential underrepresentation of CB1-positive type-1 synapses with our method, we did not attempt to quantify the relative proportion of CB1-positive boutons making type-1 or type-2 synapses.

In previous studies of rodent and monkey cortical structures (Bodor et al., 2005; Eggan and Lewis, 2007; Eggan et al., 2010; Katona et al., 2000), strongly CB1-immunopositive axons and boutons were shown to originate from local cortical GABAergic neurons (e.g. Marsicano and Lutz 1999). In the human cortex, we have tested the network of strongly CB1-immunoreactive axons for the presence of VGAT in boutons and most of them were immunopositive for both molecules (Fig 4a), but there were a large number of VGAT-positive boutons which were CB1-immunonegative (Fig 4a). This is in agreement with the transcriptomic data (Fig 1) showing that only certain types of GABAergic neuron express high levels of CB1 transcript.

**FIGURE 4.**
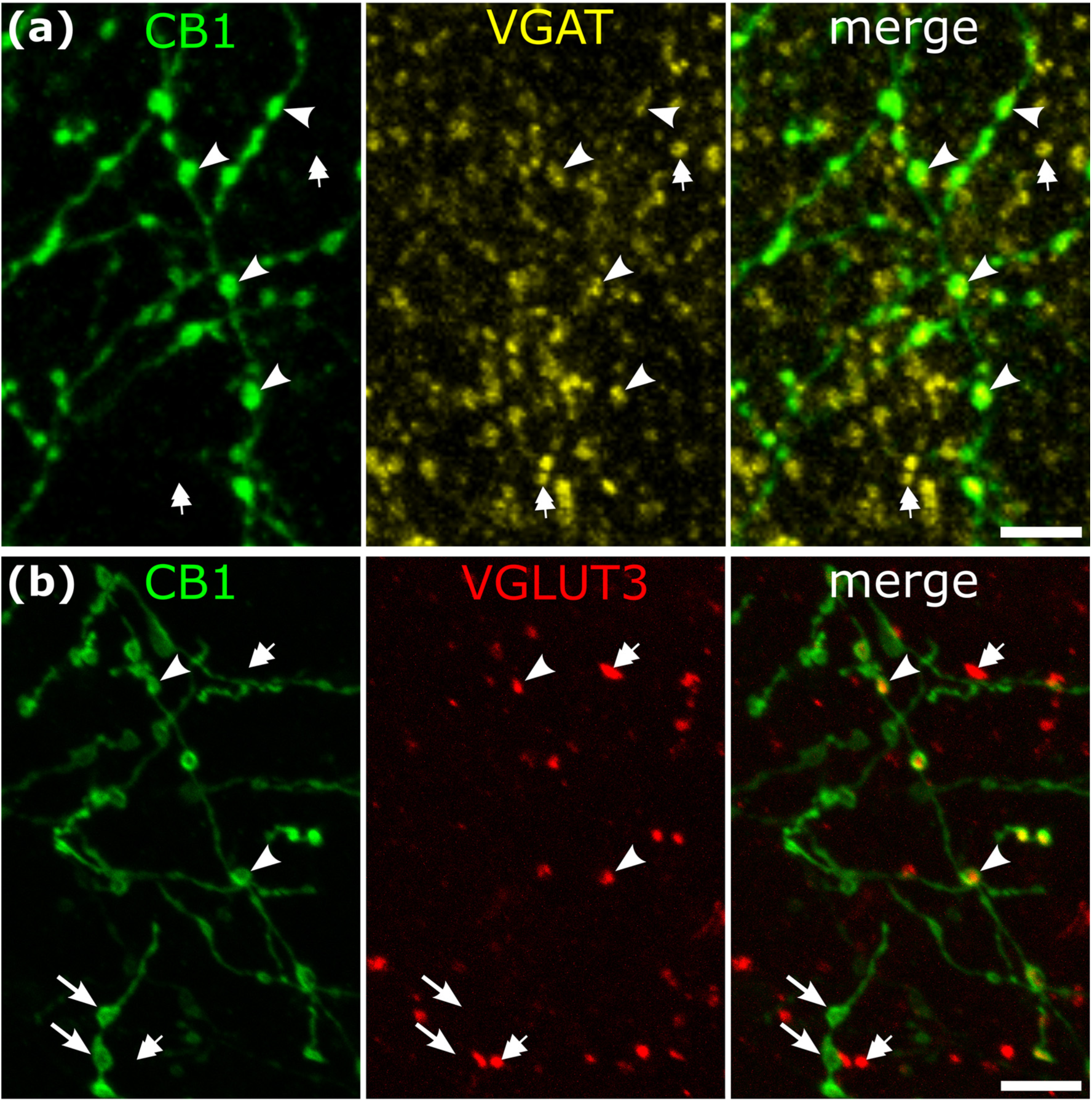
Neurotransmitter characteristics of strongly CB1-immunoreactive axons in layers 2-3 of human neocortex. **(a)** Most strongly CB1-immunoreactive boutons are VGAT-positive, hence GABAergic (e.g. arrowheads) Many VGAT-positive boutons are CB1-negative (e.g. double arrows). Left medial temporal gyrus. **(b)** Some strongly CB1-immunoreactive boutons are immunopositive for VGLUT3 (e.g. arrowheads), others are VGLUT3-negative (e.g. arrows); additional boutons are VGLUT3-positive but CB1-negative (e.g. double arrows). Middle frontal gyrus, patient J. Immunofluorescence laser scanning images, maximum intensity projection, Z stacks of 2.4 μm (a) and 3.1 μm (b) total thickness. Scale bars: 5 μm.

### 3.4 Some CB1-immunopositive boutons are immunoreactive for VGLUT3

In the rodent hippocampus and neocortex, CB1 is strongly expressed in axons of most CCK-expressing GABAergic interneurons (Katona et al 2000). Some of these also express VGLUT3 (Fasano et al., 2017; Favier et al., 2021; Omiya et al., 2015; Somogyi et al., 2004) and use both GABA and glutamate as transmitters (Fasano et al., 2017; Pelkey et al., 2020). Also, amongst the CNR1 expressing interneurons in the samples from multiple cortical areas, four groups expressed some level of SLC17A8 (Fig 1a). Therefore, we tested if in the human neocortex some strongly CB1-immunopositive boutons reacted for the VGLUT3 protein. Double immunofluorescence labelling (Fig 4b) and high-resolution individual bouton analysis (n=1542) in layers 2-3 of two patients (patient C, left middle temporal gyrus n=676; patient J left middle frontal gyrus n=866) revealed that, on average, about a quarter (patient C, 35.3%; patient J, 20.0%) of the strongly CB1-positive boutons (C, n=204; J, n=369) were also VGLUT3 immunopositive (CB1/VGLUT3+, C, *n*=72; J, n=73). About 13% (patient C, 13.2%; patient J, 13.0%) of VGLUT3 positive boutons (C, n=544; J, n=570) were strongly CB1-immunoreactive. Quantitative distributions of immunoreactive terminals in the categories were different in the two cortical areas from these two patients (Fisher’s exact test, *p* = 9.3×10^-10^, n=1542).

### 3.5 Synaptic targets, origins and diversity of VGLUT3-positive terminals

In the rodent hippocampus, many of the VGLUT3 positive GABAergic terminals are provided by CCK-expressing basket cells innervating pyramidal cell bodies and proximal dendrites (Klausberger et al., 2005; Somogyi et al., 2004) and making Gray’s type-2 synapses (Klausberger et al., 2005). In the rodent neocortex, some CCK-expressing interneurons have also been named as ‘basket cells’ (Freund et al 1986; Kubota and Kawaguchi 1997; Nunzi et al., 1985). Our immunofluorescence and light microscopic immunperoxidase analyses did not reveal any obvious basket formations around neuronal cell bodies by CB1 boutons or or VGLUT3 double positive boutons (Figures 2b, 4b) in the two cortical areas studied. Therefore, to test for the postsynaptic targets of VGLUT3-positive terminals we carried out electron microscopic immunoperoxidase analysis in three samples (patients P, inf. temp. gyrus; S, midl. temp. gyrus; T,inf. front. gyrus, Table 1). We expected to find mostly type-2 synapses as made by most identified human GABAergic cortical interneurons (Boldog et al., 2018; Kisvarday et al., 1990; Lukacs et al., 2023; Varga et al., 2015). Surprisingly, only 40±6% (SE) VGLUT3-positive boutons made type-2 synapses (total *n*=104; patient P, *n*=33; patient S, *n*=41; patient T, *n*=30). The rest made type-1 synapses (60±6%), similar to those of cortical glutamatergic neurons (Somogyi 1978) and primate thalamo-cortical afferents (Freund et al., 1989). This was unexpected as only one group of glutamatergic neurons, Exc L6 THEMIS CC6orf48, shows very weak SLC17A8 gene expression in the processed human cortical areas (http://celltypes.brain-map.org/rnaseq/).

#### Type-2 synapses

We have quantified the relative proportions of the synaptic targets of type-2 synapses, which we assume mainly originate from GABAergic interneurons (Figure 5f) and showed very thin postsynaptic membrane specialisations (Figure 5a-e). Of the 41 synapses from three cortical areas only one targeted a pyramidal cell body (Figure 5a), most postsynaptic elements were small dendritic shafts (Figure 5c,e), rarely apical dendrites of pyramidal cells (Figure 5b) or dendritic spines (Figure 5d, *n*=5). When the dendritic spines could be followed in serial sections, they also received a type-1 synapse each (Figure 5d) with thick postsynaptic membrane specialisation suggesting glutamatergic neurotransmission. The proportion of dendritic shaft to spine targets were not significantly different in the three samples (Figure 5f, Fisher’s exact test, p = 1, *n*=40), hence they were pooled. Overall, VGLUT3-positive boutons making type-2 synapses targeted mostly dendritic shafts (82%) and to lesser extent dendritic spines (16%) and a soma (2%). We assume that most of these type-2 synapses are made by VGLUT3-expressing GABAergic neuronal types, which partially overlap with the larger population of CB1-experssing GABAergic neuronal types. We have compared the postsynaptic target distributions of these two populations based on the electron microscopic data (Figure 3h; Figure 5f), and the distributions were not different (CB1 total *n*=82; VGLUT3, total *n*=41; Fischer’s exact test, *p*=0.156).

**FIGURE 5.**
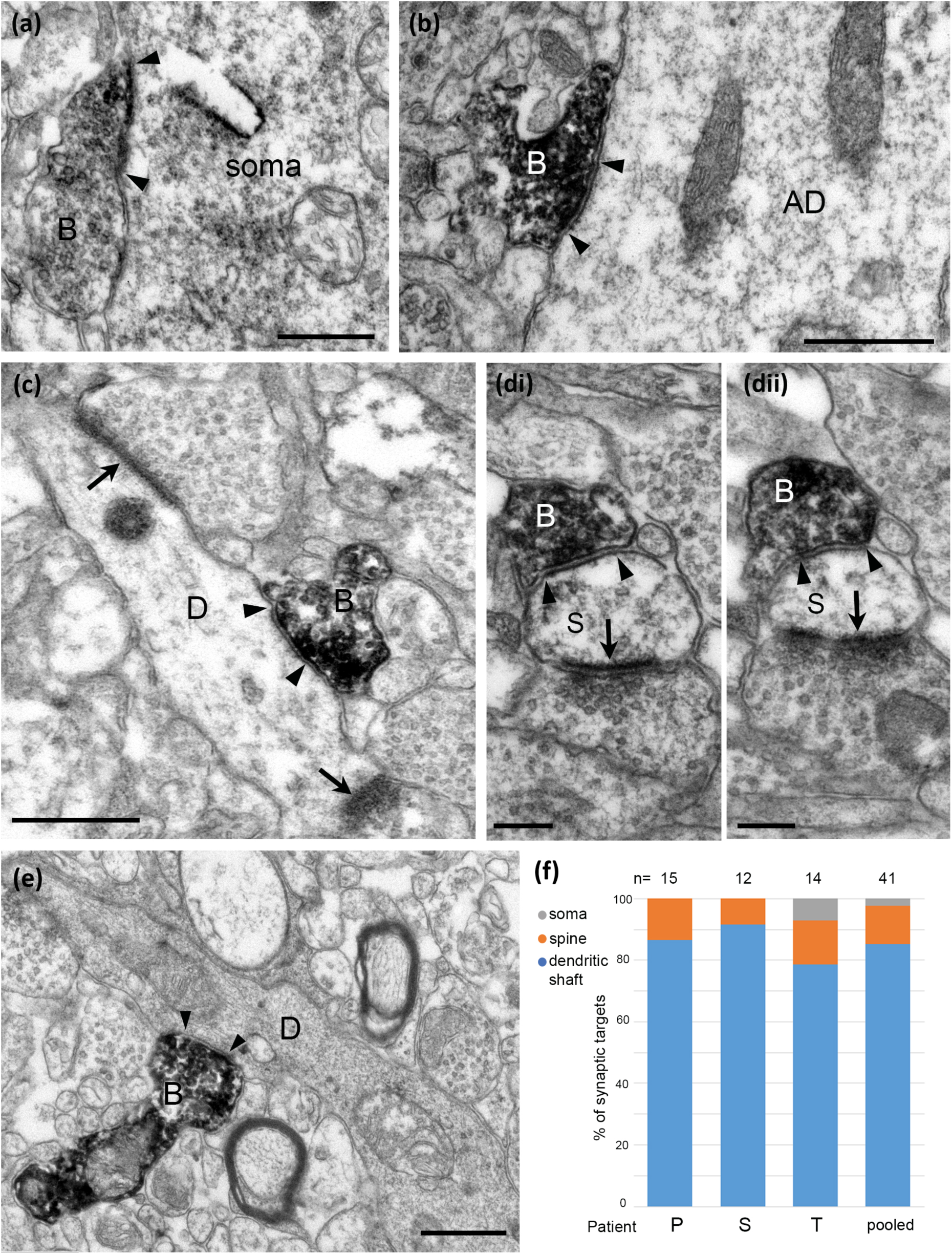
Electron micrographs of the synaptic targets and quantitative distribution of VGLUT3 expressing nerve terminals forming type-2 synapses. Immunoperoxidase reactions show VGLUT3-positive terminal boutons (B) filled with electron-opaque DAB-deposit (black); the synaptic junctions are marked by arrowheads. **(a)** A neuronal soma receives a type-2 synapse from VGLUT3-positive bouton. **(b)** A large apical dendrite (AD) as postsynaptic target. **(c)** A small diameter dendritic shaft (D) receives two type-1 synapses (arrows) in addition to the synapse from a VGLUT3-positive terminal and it likely originates from an interneuron. **(di, dii)** Serial sections of a dendritic spine (S) receiving a type-2 synapse from the VGLUT3-positive bouton and a type-1 synapse (arrow) from an unlabelled bouton. (e) A small diameter dendritic shaft (D) with electron opaque cytoplasm is innervated by a VGLUT3-positive terminal. **(f)** Quantification showing that most of the synaptic targets of type-2 synapses were dendritic shafts and far fewer were dendritic spines in samples from three patients. Scale bars: (a-c, e) 0.5 μm; (d), 0.2 μm.

#### Type-1 synapses

The proportions of identified postsynaptic dendritic shaft and spine targets of VGLUT3-positive boutons making type-1 synapses with extensive postsynaptic membrane specialisation (Figure 6), differed from those making type-2 synapses (Fisher’s exact test, p = 6. 1×10^-11^, (*n*=102) odds ratio, 0.0435, 95% confidence interval 0.011 - 0.138). Of the 62 synapses with identified targets (*n*=1 unidentified) pooled from three cortical areas, most targets (78%) were dendritic spines (Figure 6b-e), the rest dendritic shafts (22%, Figure 6a). Some dendritic shafts received other type-1 synapses and may have originated from interneurons (Figure 6a). Some dendritic spines had large spine apparatus (Figure 6b), or additional multivesicular bodies (Figure 6e); small spines without a spine apparatus (Figure 6c) and sessile spines without a spine neck (Figure 6d) were also targets. In patient P, only one of 18 synapses was on a dendritic shaft in the inferior temporal gyrus; the three samples did not differ in the proportion of dendritic shaft to spine synaptic targets (Fisher’s exact test, p = 0.11, *n*=62).

**FIGURE 6.**
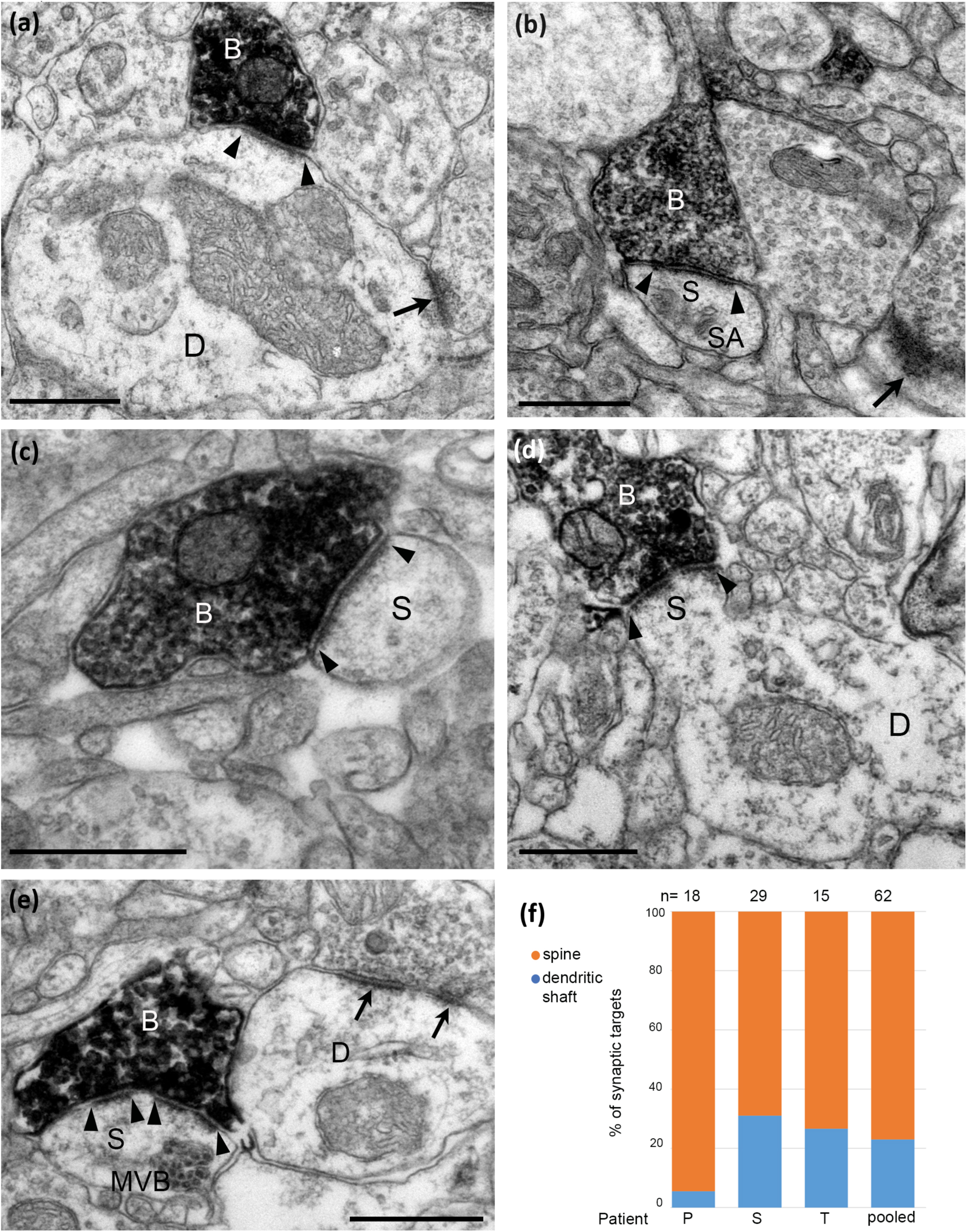
Electron micrographs of the synaptic targets and quantitative distribution of VGLUT3 expressing nerve terminals forming type-1 synapses. Immunoperoxidase reactions show VGLUT3-positive terminal boutons (B) filled with electron-opaque DAB-deposit (black); the synaptic junctions are marked by arrowheads. **(a)** A large dendritic shaft (D) receives a type-1 synapse from VGLUT3-positive bouton and an additional type-1 synapse from an unlabelled terminal (arrow). **(b-e)** Dendritic spines as synaptic targets. In **(b)** the spine (S) has a spine apparatus (SA) and another spine also receives a type-1 synapse (arrow); in **(d)** the innervated sessile spine (S) originates from a dendritic shaft (D); in **(e)** the spine (S) contains a multivesicular body (MVB) and the VGLUT3-positive bouton makes a perforated synapse as does a nearby bouton (arrows) with a dendritic shaft (D). **(f)** Quantification showing that most of the synaptic targets of type-1 synapses were dendritic spines and far fewer were dendritic shafts in samples from three patients. Scale bars: (a-f), 0.5 μm.

The VGLUT3-positive terminals making type-1 synapses resembled those made by cortical pyramidal cells and other glutamatergic neurons such as spiny stellate cells, all of which express VGLUT1 (Vigneault et al., 2015; Figure 1). Therefore, we tested if some VGLUT3-positive boutons also contained VGLUT1 by double immunofluorescence labelling and high-resolution confocal microscopy in layers 2-3 of two patients (patient S, mid. temp g.; U, sup. front. g.). A total of 403 VGLUT3 positive boutons (patient S, *n*=272; U, *n*=131) were tested, each bouton in several optical slices, for the presence of VGLUT1 immunoreactivity (Figure 7a). On average, 52% of VGLUT3-positive boutons (patient S, 46%; patient U, 58%) were also immunopositive for VGLUT1. Because of the high density of VGLUT1-positive boutons, we did not count those that expressed only this vesicular transporter.

**FIGURE 7.**
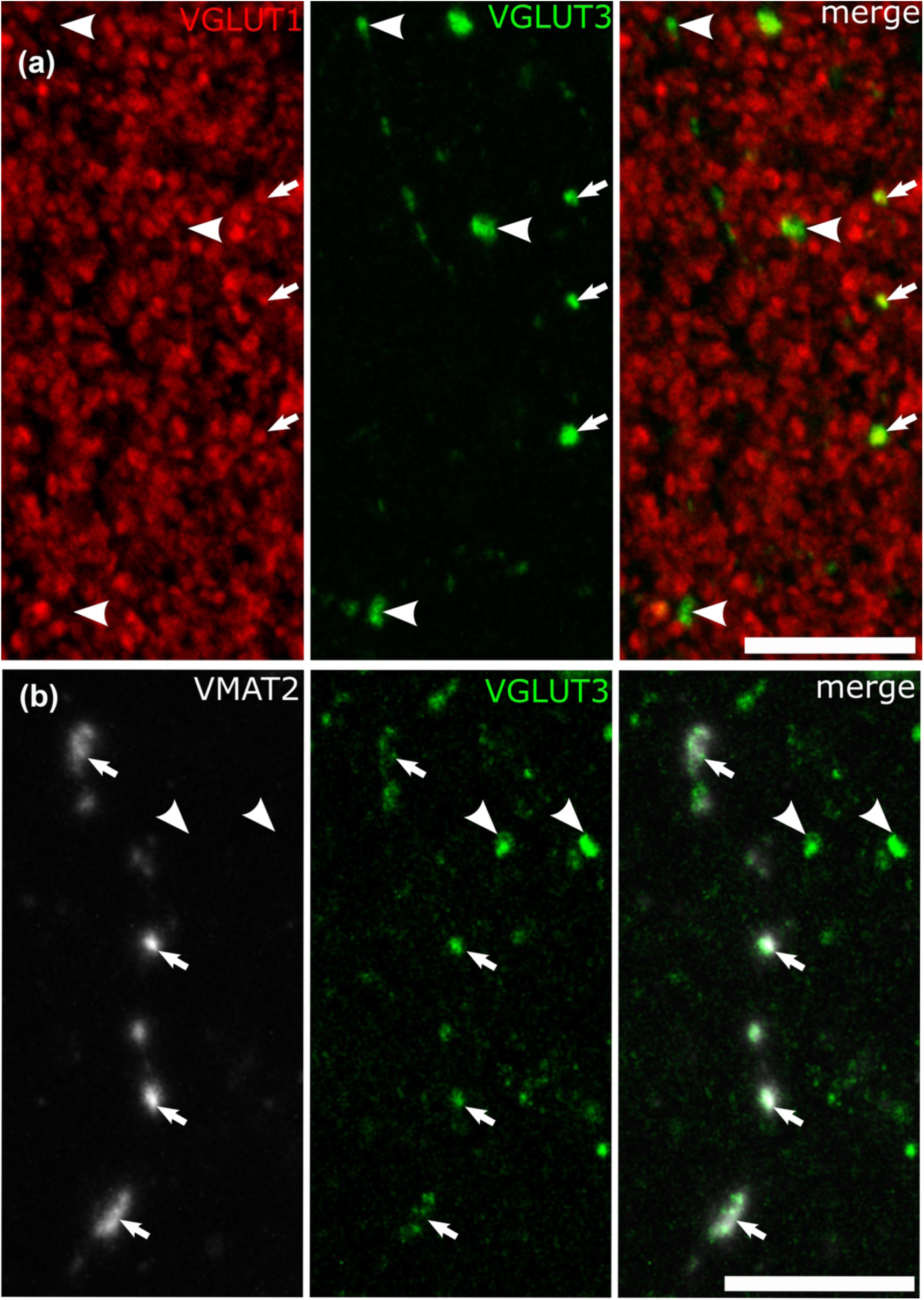
Co-expression of VGLUT3 with VGLUT1 or VMAT2 in some cortical axonal varicosities. **(a)** A high density of VGLUT1-positive terminals include infrequent VGLUT3-positive boutons (arrows), but other VGLUT3-positive boutons are immunonegative for VGLUT1 (e.g. arrowheads) in layer 3. (Patient S, left middle temporal gyrus). **(b)** Some of the large VGLUT3-positive boutons (green, arrows) are positive for VMAT2, but other boutons are immunonegative for VMAT2 (e.g. arrowhead) (Patient S, left middle temporal gyrus). Maximum intensity projection of optical slices with total Z-stack depth of 2.6 μm (a) and 8.5 μm (b). Scale bars: 10 μm.

An additional source of VGLUT3-expressing nerve terminals in the cortex of rodents is the dorsal and medial raphe nuclei, where some serotonergic neurons express VGLUT3 (Gras et al., 2002), as they do in humans (Vigneault et al., 2015), and their terminals can be identified by vesicular monoamine transporter-2 (VMAT2) (Somogyi et al., 2004). In two patients (patient L, inf. temp. g.; S, mid. temp. g.), we used double immunofluorescence labelling for VMAT2 and VGLUT3 and high-resolution confocal microscopy in layers 2-3. Most boutons immunolabelled for VMAT2 were very small and not labelled for VGLUT3, but there were rare, distinct fibres with large boutons, which were consistently labelled for VGLUT3 (Figure 7b). It is likely that these correspond to the boutons of 5-HT neurons of the raphe nuclei, as in rodents. The numerous small VMAT2-positive boutons could originate from dopaminergic and/or noradrenergic fibres, which are not known to express VGLUT3.

### 3.6 Interneuronal types immunopositive for CB1 and/or VGLUT3

There is a large expansion of ADARB2 expressing interneurons grouped by transcriptomic profiles in the human cortex (Figure 1), as compared to the mouse, most of which express CNR1 and 2 groups also VGLUT3 (Bakken et al., 2020; Hodge et al., 2019; Lee et al 2023; BRAIN Initiative Cell Census Network 2020). Each group probably evolved driven by as yet unknown synaptic input/output specialisations. To define cells within these groups, we visualised their dendritic and axonal arborizations by recording and labelling individual neurons *in vitro* in layers 1-3 then determining their expression of combinations of marker molecules by immunohistochemistry (Figures 8-15). We have selected non-pyramidal cells for testing of their immunoreactivity for CB1 from two previously published *in vitro* recorded and biocytin labelled sets of interneurons. Lukacs et al., (2023) reported 356 visualised interneurons in layers 2-3 from multiple cortical areas, whereas Field et al., (2021) recorded and labelled interneurons mostly in layer 1 (*n*= 84), and 5 cells in layer 2. Here, a total of 120 interneurons that had some of their axons visualised were tested with antibodies to CB1; 89 from the set of Lukacs et al., (2023) and 31 from the set of Field et al., (2021). The neurons were not randomly chosen for testing, but selected on the predicted probability of expressing CB1, or being axo-axonic cells. The latter population of 8 axo-axonic cells were chosen for comparison as they represent a relatively homogeneous and well recognisable cell type (Somogyi, 1977; 1982). Other interneurons with axons apparently targeting neuronal cell bodies were tested, because some so-called ‘basket cells’ express strong CB1 immunoreactivity in rodents (Dudok et al., 2015; Katona et al., 1999; Lee and Soltesz, 2011). Neurons with their soma in layers 2-3 and having descending axons were also chosen because some of these express CCK and/or VIP and/or CR, populations known to express CB1 receptors (Figure 1). Finally, some rosehip cells are known to express CCK (Boldog et al., 2018), which is frequently co-expressed with CB1 (Figure 1), were tested.

**FIGURE 8.**
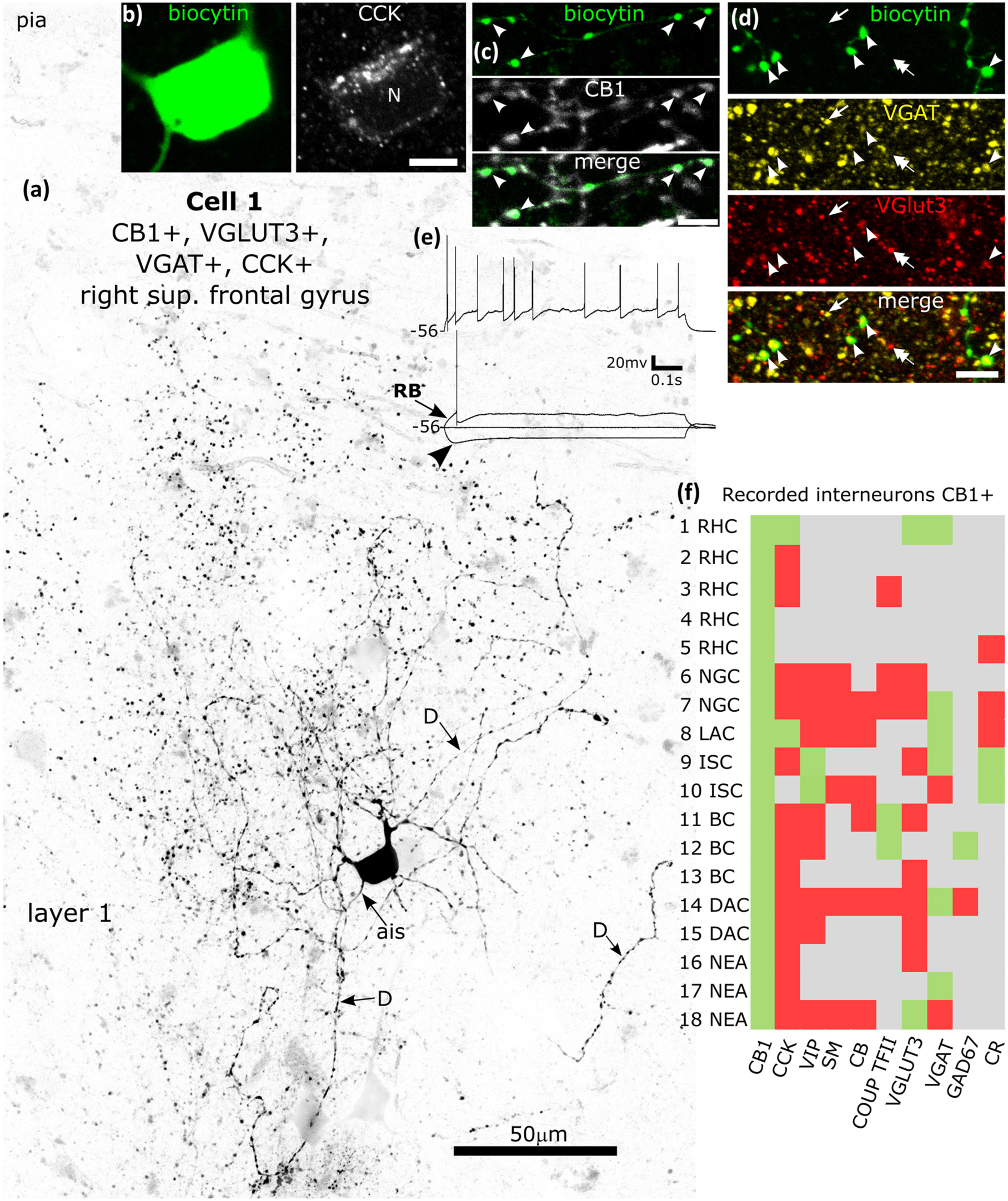
Immunohistochemical molecular testing of GABAergic interneurons recorded in vitro in whole-cell mode. **(a)** A rosehip cell is CB1-positive in layer 1 (cell 1 in Figure 8f). Reverse contrast LSM image of the biocytin labelled cell; note the high density of large boutons (dots). The axon initial segment (ais) and three dendrites (D) are indicated. **(b)** The soma is immunopositive for CCK (N, nucleus). **(c)** The axon and boutons are CB1-positive (e.g. arrowheads). **(d)** Boutons of the cell (green, arrowheads) are both VGAT-(yellow) and VGLUT3-positive (red) amongst additional VGAT (e.g. arrow) and/or VGLUT3-positive (e.g. double arrow) boutons. Maximum intensity projections of optical slices, Z-stack total depth: (a), tiled, 20 μm; (b), 4.4 μm; (c), 0.35 μm; (d), 3.5 μm. **(e)** Voltage responses of the cell to holding current and current injections of rheobase (RB), RB +100 pA (upper trace), and holding current −100 pA, showing a small voltage sag (arrowhead) in response to the hyperpolarising current injection. The cell fired irregularly with spike frequency adaptation and strong spike amplitude accommodation when continuously depolarised. **(f)** Immunoreactivities in all recorded and visualised CB1-positive interneurons (n=18) tested for 1-8 molecules: green, immunopositive; red, immunonegative; grey, not tested/inconclusive. Most neurons tested were VGAT-positive. BC, basket cell; DAC, descending axon cell; ISC, interneuron specific cell; LAC, loose axon cell; NEA, not enough axon; NGC, neurogliaform cell; RHC, rosehip cell; Scale bars, (b-d), 5 μm.

The soma, some dendrites and some axons were recovered from 17 CB1 immunopositive cells as tested on their axons (Figure 8f). For one cell only the axon was recovered (No 15, Figure 8f), which was a cell with descending translaminar axon, originating in layer 2, where the cell was recorded. The cell bodies of the other cells were in layer 1 (*n*=7), layer 2 (*n*=3) or layer 3 (*n*=7). The cells in the recorded slices were cut into 3-6 sections. To help interneuron type characterisation, we immunoreacted selected parts of the cells from individual sections, each section being reacted for several molecules, if necessary multiple times. In addition to CB1, some of the axons were tested for CCK, VIP, SM, VGAT, GAD67, VGLUT3 and calretinin; somata were tested for CCK, SM, calretinin, calbindin and COUP-TF2 (Figure 8f). An immunopositive or immunonegative result for a given cell was accepted if nearby non-recorded cells or axons showed high quality immunoreactivity. Nevertheless, we cannot exclude that the whole cell recording conditions adversely affected the amount of the molecule being tested in a given cell, resulting in a false negative score, or that the amount of molecule was below the threshold of detectability. Hence, immunonegative results need to be interpreted with caution.

We recognise cell types mainly based on the distribution, shape and density of the axonal branches and boutons. Three CB1-positive cells had not enough axon recovered (NEA, Cells 16-18, Figure 8f) for categorisation. The remaining 15 CB1-positive cells showed six distinct axonal patterns.

#### Rosehip cells

(RHC, *n*=5) (Boldog et al., 2018; Field et al., 2021) had a high volume-density of large boutons densely packed along the axonal branches mainly in layer 1 in our sample (Figures 8 and 9). One of the cells was immunopositive for CCK, VGLUT3 and VGAT (Cell 1, Figure 8), two others (Cell 2, Figure 9 and cell 3) were CCK-negative, one of these was also COUP-TF2 negative.

**FIGURE 9.**
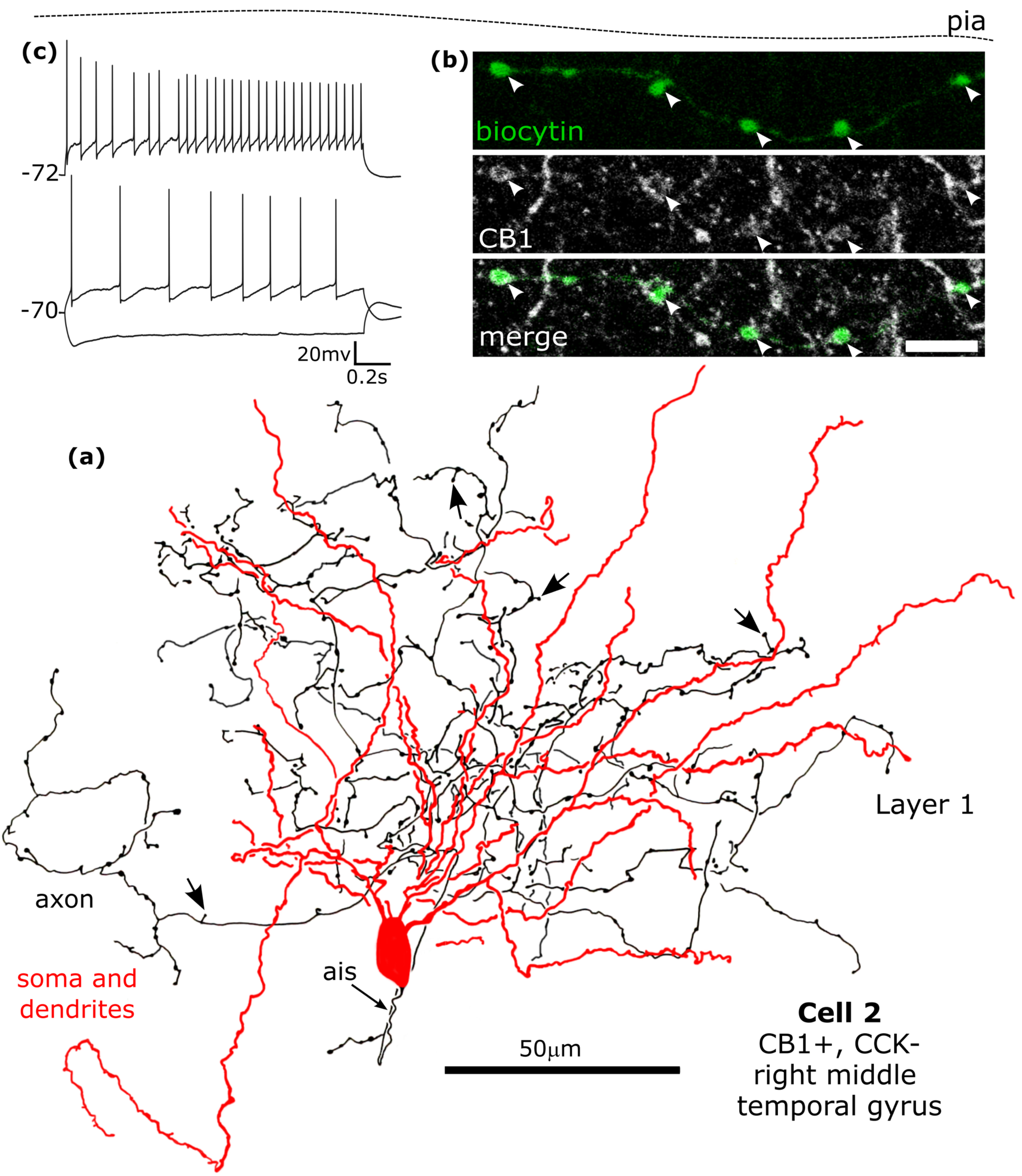
A CB1-immunopositve rosehip neuron recorded in vitro and labelled by biocytin that was CCK-immunonegative in layer 1. **(a)**. Reconstruction of the recovered parts of the cell (cell 2 in Figure 8f) with soma and dendrites (red) and a high density of axon with frequent stalked boutons (arrows). **(b)** The biocytin-labelled axon and boutons (green) were CB1-positive (e.g. arrowheads). Maximum intensity projection of optical slices, Z-stack total depth: (b), 2.1 μm. **(c)** Voltage responses of the cell to current injections of rheobase, RB +100 pA (upper trace), and holding current −100 pA; a small voltage sag was detected to the hyperpolarising current injection. The cell fired regularly with moderate spike frequency acceleration and strong spike amplitude accommodation when continuously depolarised. Scale bar: (b) 5 μm.

#### Neurogliaform cells

(NGC, *n*=2, cells 6 and 7 Figure 8f) located in layer 1 (Tamas et al., 2003) had high density of frequently branching ‘wavy’ axons in and around the dendritic field, with smaller and sparser boutons than rosehip cells (Boldog et al., 2018; Field et al., 2021; Kisvarday et al., 1990). One of them was immunopositive for VGAT; both of them were immunonegative for all tested other molecules.

One CB1-positive neuron had a sparse *loose axon* (LAC, Figure 10) with straight collaterals running for hundreds of micrometres and not forming clusters. This cell was positive for CCK and VGAT (Cell 8, Figure 8f).

**FIGURE 10.**
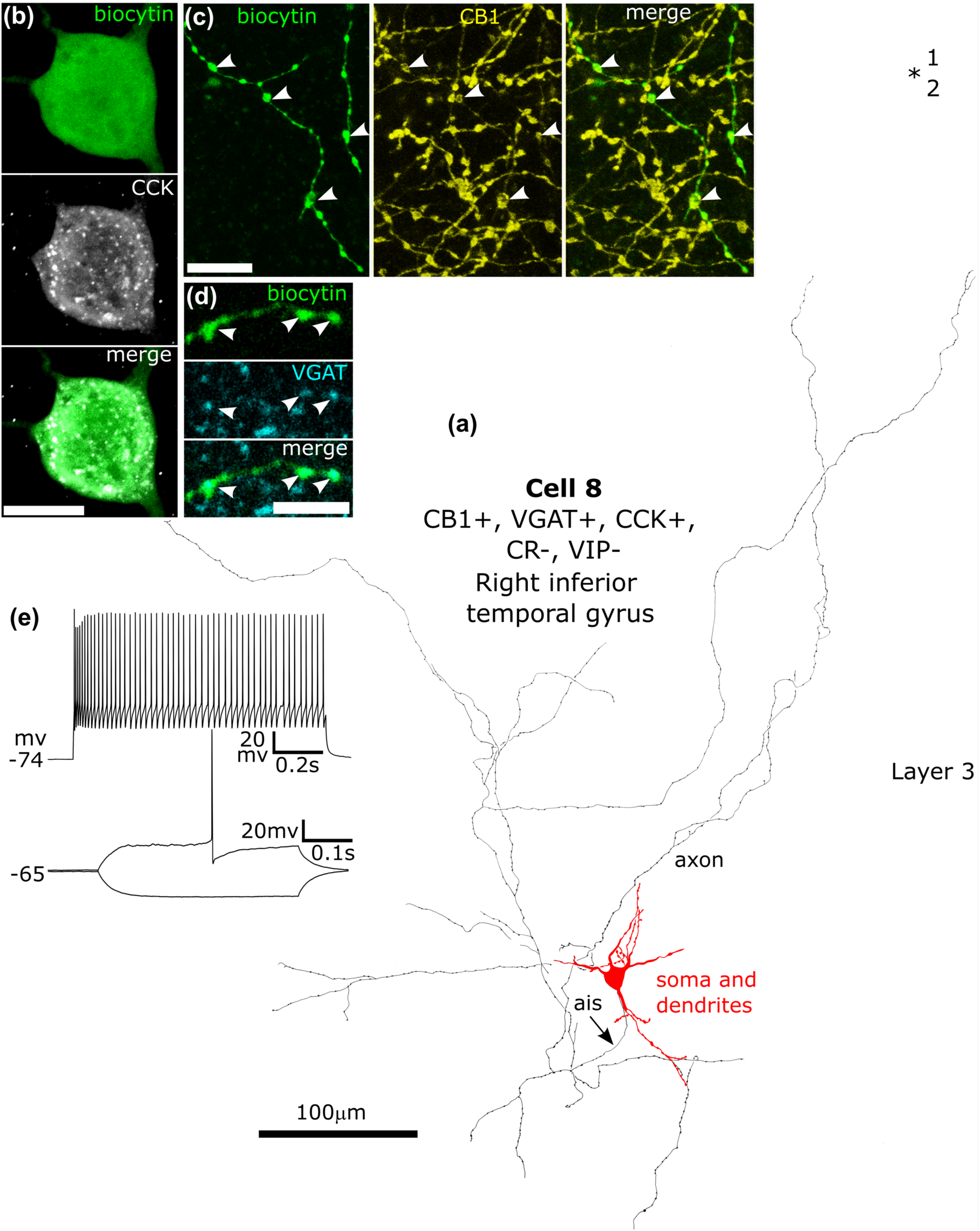
A CB1/VGAT/CCK-immunopositive neuron recorded in vitro and labelled by biocytin in layer 3. **(a)**. Reconstruction of the recovered parts of the cell (cell 8 in Figure 8f) with soma and dendrites (red) and a low-density loose axon. **(b)** The soma of the cell (green) showed granular immunoreactivity for CCK in the cytoplasm. **(c)** The biocytin-labelled axon (green) and boutons were CB1-positive (e.g. arrowheads). **(d)** The boutons (green, arrowheads) show immunoreactivity for VGAT (blue). Max intensity projections of optical slices in Z stack, total depth: (b), 10.5 μm; (c), 3 μm; (d), 0.35 μm. **(e)** Voltage responses of the cell to current injections of rheobase (400ms duration), RB +100 pA (upper trace, 1 s duration), and holding current −100 pA; no voltage sag was detected to the hyperpolarising current injection. The cell fired regularly with moderate spike frequency adaptation and an initial spike amplitude accommodation when continuously depolarised. Scale bars: (b,c) 10 μm; (d) 5 μm.

Two putative *interneuron specific cells* (ISC, Cells 9 and 10, Figure 8f) were identified in layer 3 as innervating other interneurons on their soma and proximal dendrites with multiple boutons (Figure 11). Most GABAergic neurons innervate both glutamatergic neurons and GABAergic interneurons to differing degrees, but some interneurons heavily innervate mainly interneurons (Gabbott et al., 1997; Meskenaite 1997). The degree of synaptic target selectivity of these two recorded cells could not be tested, but some of their features suggest that they are interneuron specific to a large extent. Both cells were VIP- and also calretinin-positive, molecules that characterise some interneuron specific cells (Gabbott et al., 1997; Meskenaite 1997). Cell 9 is shown to innervate another calretinin-positive interneuron via at least 4 boutons. Cell 10 (not shown) targeted the soma and proximal dendrites of some calbindin-positive interneurons via multiple boutons, in one case by more than 10 boutons forming a basket-like configuration around a calbindin-positive soma.

**FIGURE 11.**
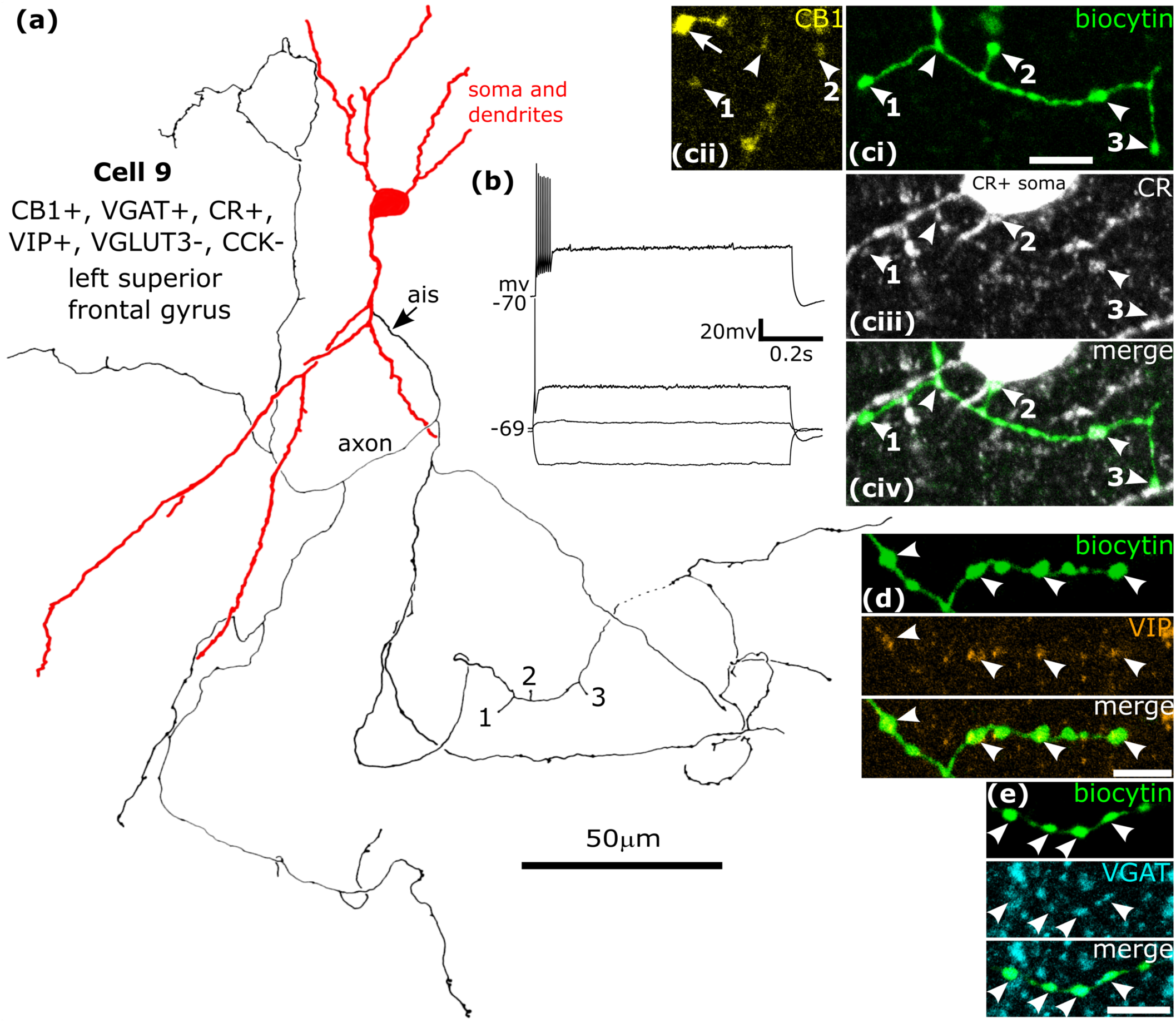
A CB1/VGAT/VIP/CR-immunopositive neuron recorded and labelled by biocytin in layer 3. **(a)**. Reconstruction of the recovered parts of the cell (cell 9 in Figure 8f) that was negative for CCK and VGLUT3, with soma and dendrites (red) and a low density axon originating from the descending dendrite (ais). Boutons 1, 2 and 3 are shown in (c). **(b)** Voltage responses of the cell to current injections +20 pA, rheobase, RB +100 pA (upper trace), and holding current −100 pA; no significant voltage sag was detected to the hyperpolarising current injection. The cell fired with an initial burst and spike amplitude accommodation when continuously depolarised. **(c)** The axonal boutons (ci, arrowheads, green, see 1, 2 and 3 in (a)) were weakly CB1-positive (civ), calretinin-positive (cii, arrowheads) and some of them contacted a CR-positive interneuron on the soma (2) and dendrites (1, 3). **(d)** The biocytin-labelled boutons (green, e.g arrowheads) were VIP-positive (arrowheads). **(e)** The boutons (green, arrowheads) were immunoreactive for VGAT (blue). Maximum intensity projections of optical slices, Z-stack total depth: (c), 2.3 μm; (d), 1.5 μm; (e), 1.7 μm. Scale bars: (c-e), 5 μm.

#### Basket cells

(BC, Figure 8f) are recognised from their axonal boutons occasionally around the cell bodies of other neurons, including pyramidal cells, and giving 2-10 boutons to a single innervated cell (Figures 12 and 13). Three such recoded cells were CB1-positive (Cells 11, 12, 13; Figure 8f) and two of them tested were also positive for the nuclear transcription factor COUP-TF2, which characterises a subset of human cortical interneurons (Varga et al., 2015). The cell body of one cell was in layer 2 (Figure 12) and those of the other two in layer 3. These basket cells had long descending axons emitting collaterals forming dense bouton clusters, some of which were around selected cell bodies. We could not ascertain if these soma targets were interneurons, pyramidal cells or both. Cell 11 had a very selective expansion of its axon in lower layer 3.

**FIGURE 12.**
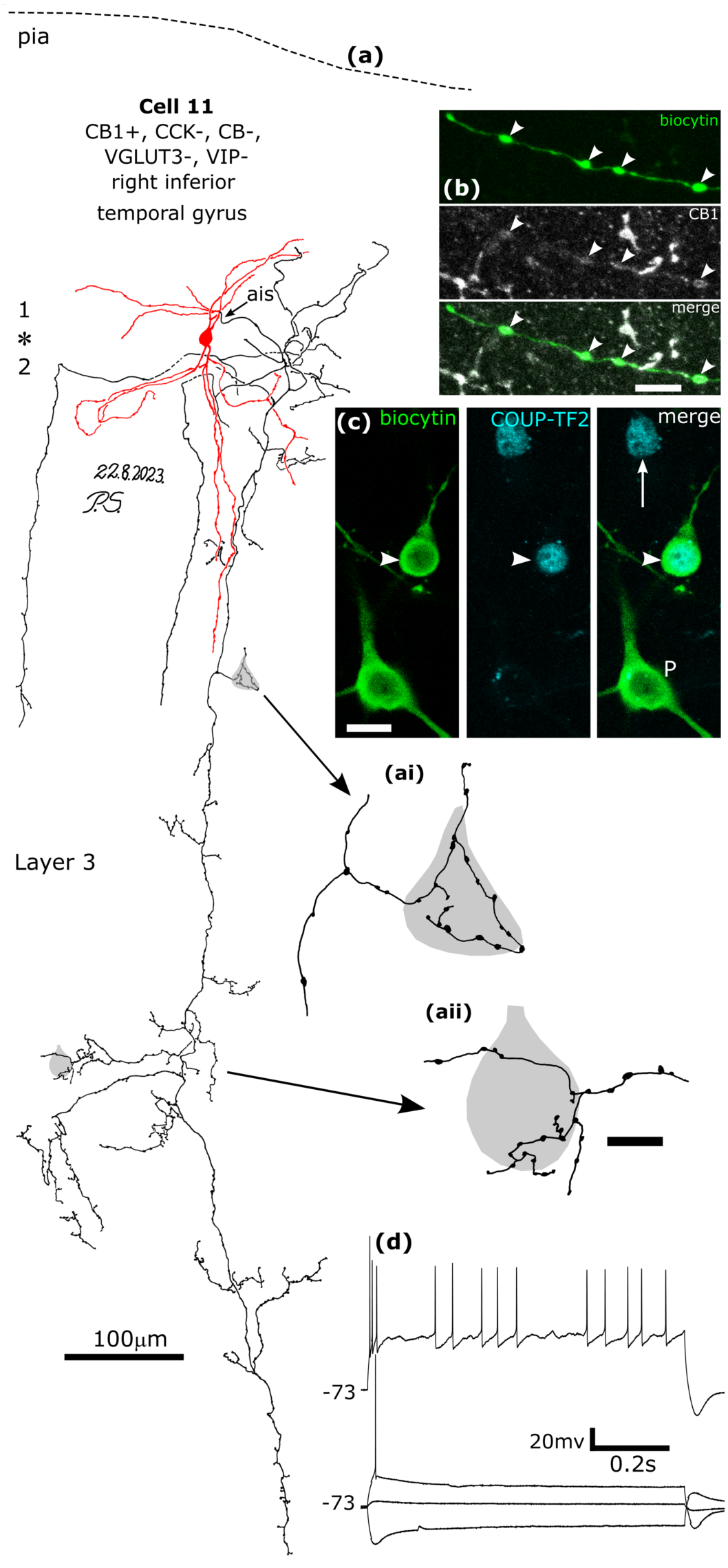
A CB1/COUP-TF2-immunopositive basket cell that was negative for CCK, CB, VGLUT3 and VIP, recorded and labelled by biocytin at the border of layers 1 and 2 (asterisk) **(a)**. Reconstruction of the recovered parts of the cell (cell 11 in Figure 8f) with soma and dendrites (red) and a descending axon originating from a dendrite (ais). The axon arborised extensively in lower layer 3 and formed clusters of boutons around select cell bodies (e.g. ai and aii). Asterisk, border of layers 1 and 2. **(b)** The axonal boutons (arrowheads, green) were weakly CB1-positive. **(c)** The nucleus of the biocytin-labelled neuron (arrowhead) was COUP-TF2-positive, as was another neuronal nucleus nearby (arrow), but the nucleus of a biocytin-labelled pyramidal cell was immunonegative (P, bottom). **(d)** Voltage responses of the cell to current injections of holding current +20 pA, rheobase, RB +100 pA (upper trace), and holding current −100 pA; a voltage sag is apparent to the hyperpolarising current injection. The cell fired with an initial burst followed by irregular firing with spike amplitude accommodation when continuously depolarised. Maximum intensity projections of optical slices, Z-stack total depth: (b), 2.9 μm; (c), 1.0 μm; Scale bars: (ai, aii) 5 μm; (b, c), 10 μm.

**FIGURE 13.**
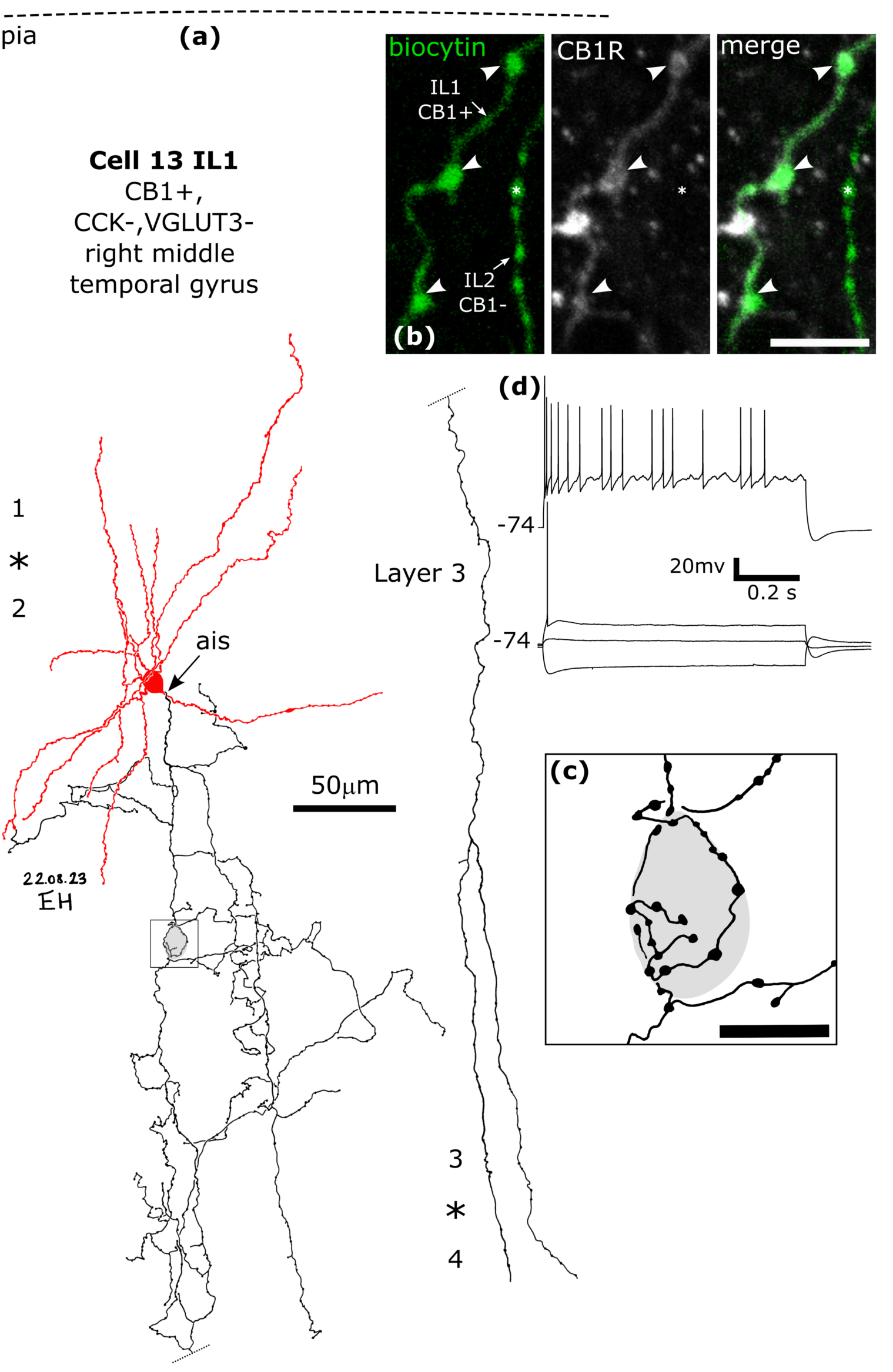
A CB1-immunopositive basket cell (cell 13 IL1) that was negative for CCK and VGLUT3, recorded and labelled by biocytin in layer 2. Asterisk, border of layers 1 and 2. **(a)**. Reconstruction of the recovered parts of the cell (cell 13 in Figure 8f) with soma and dendrites (red) and a descending axon originating from the soma (ais). The axon arborised in layer 3 and formed clusters of boutons around select cell bodies (e.g. framed area shown in (c)), then descended (dotted line continued on right) to layer 4 (border, asterisk) and could not be followed further. **(b)** The axonal boutons (arrowheads, green) were weakly CB1-immunopositive, but the boutons of a nearby labelled interneuron (IL2, e.g. asterisk) were immunonegative. **(c)** A cell body is innervated by a ‘basket’ of boutons. **(d)** Voltage responses of the cell to current injections of holding current +50pA, rheobase, RB +100 pA (upper trace), and holding current −100 pA; a voltage sag is apparent to the hyperpolarising current injection. The cell fired an irregular train of action potentials with spike amplitude accommodation when continuously depolarised. Maximum intensity projections of optical slices, Z-stack total depth: (b), 0.4 μm; Scale bars: (b), 5 μm; (c), 10 μm.

Two other cells with radial *descending axons* (DAC, Cells 14 and 15, Figures 8f and 14) did not show clustering of boutons around cell bodies. From one cell (Cell 15) only the CB1-positive axon was recovered originating in layer 2. Cell 14 was immunopositive for VGAT, but both cells 14 and 15 were immunonegative for all other tested molecules (Figure 8f). Cell 14 was recorded together with another interneuron to which it was presynaptic. Pairs of action potentials of DAC 14 evoked by small depolarising current injections, evoked a small compound IPSP of 1.4 ± 1.6 mV (mean ± sd, n = 50 sweeps) in the postsynaptic interneuron nominally clamped at −53 mV (Figure 14e). The axon of DAC 14 contacted the postsynaptic interneuron with 4 boutons at relatively proximal dendritic sites. The two cells had very similar firing patterns, similar bitufted dendritic trees and the postsynaptic interneuron also had a descending radial axon, which was however CB1-negative. The postsynaptic cell’s axon was not in contact with the recovered dendrites of Cell 14. In summary, CB1-immunopositive interneurons in layers 1-3 comprise of at least 6 cell types based on their axonal features. Their GABAergic nature is demonstrated by immunoreactivity for VGAT (6/8 tested cells) or glutamate decarboxylase (1/2 tested cells).

**FIGURE 14.**
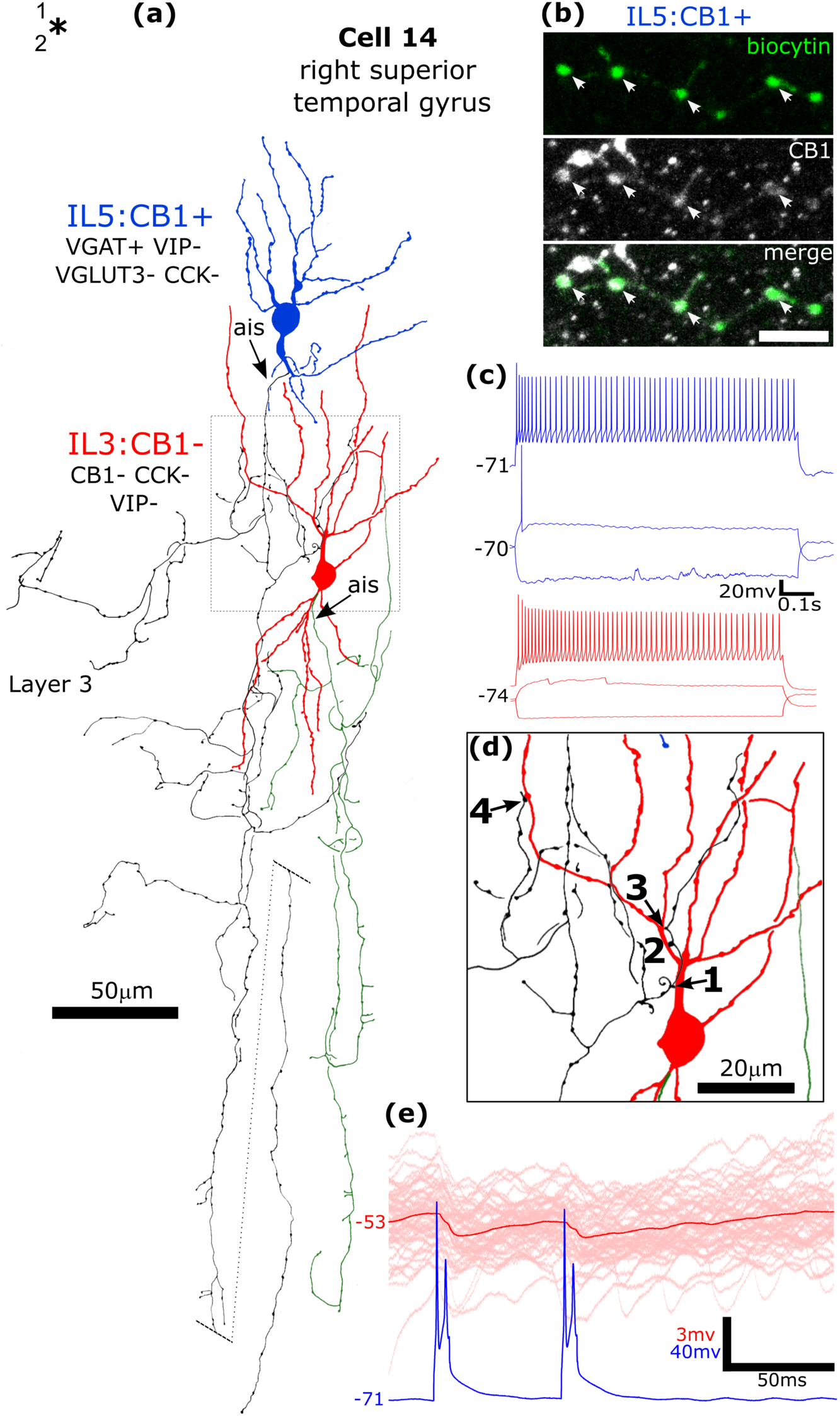
A CB1/VGAT-immunopositive interneuron (IL5, blue) innervates a CB1-negative interneuron (IL3) in layer 2/3 of the human right superior temporal gyrus. **(a)**. Reconstruction of the recovered parts of the cells with soma and dendrites (blue, IL5, cell 14 in Figure 8f; red IL3) and a low density, descending axons (black IL5; green IL3). The axon of the CB1-positive interneuron IL5 contacts the dendrites of IL3 at 4 dendritic sites, framed area (1-4) shown in **(d).** The descending axon of IL5 is shown shifted radially (dotted line). Asterisk, border of layers 1 and 2. **(b)** Confocal microscopic image of the biocytin labelled axon and boutons (green) of cell IL5 showing CB1-immunoreactivity (arrows). Maximum intensity projections of optical slices, Z-stack total depth: 2.4 μm. **(c)** Voltage responses of the cells to current injections close to rheobase, RB +100 pA (upper traces), and holding current −100 pA; no voltage sags were detected to the hyperpolarising current injection. **(e)** Single sweeps (pink, n=50) and average synaptic response (red line) of cell IL3 to action potentials of IL5 (single sweep) show a relatively small IPSP. The presynaptic CB1+ neuron fired doublet spikes to a 10 ms depolarising current pulse, and both spikes contributed to evoking the IPSP. Scale bar: (b), 5 μm.

Among the tested 101 CB1-immunonegative cells, there were 6 calbindin-positive *double bouquet cells* (Lukacs et al., 2023), 8 parvalbumin-positive *dendrite targeting cells* (DTC, Lukacs et al., 2023), 5 *basket cells* (Field et al., 2021), 2 *rosehip* and *2 neurogliaform cells* (Field et al., 2021), 3 *cells with descending axons*, 3 *cells with stalked axons* in layer 1 (Field et al., 2021) and 6 *axo-axonic cells* (Figure 15). The firing parameters of some of these cells have been reported (Field et al., 2021; Lukacs et al., 2023), here we did not compare them with the CB1-immunopositive cells. A remarkable characteristic of the CB1-immunonegative *axo-axonic cells*, which were double immunopositive for parvalbumin and calbindin with one exception, is their delayed firing to suprathreshold depolarization after the first action potential (Figure 15), which was consistent for all the 7 cells recorded and may serve as a signature for this cell type in future studies.

**FIGURE 15.**
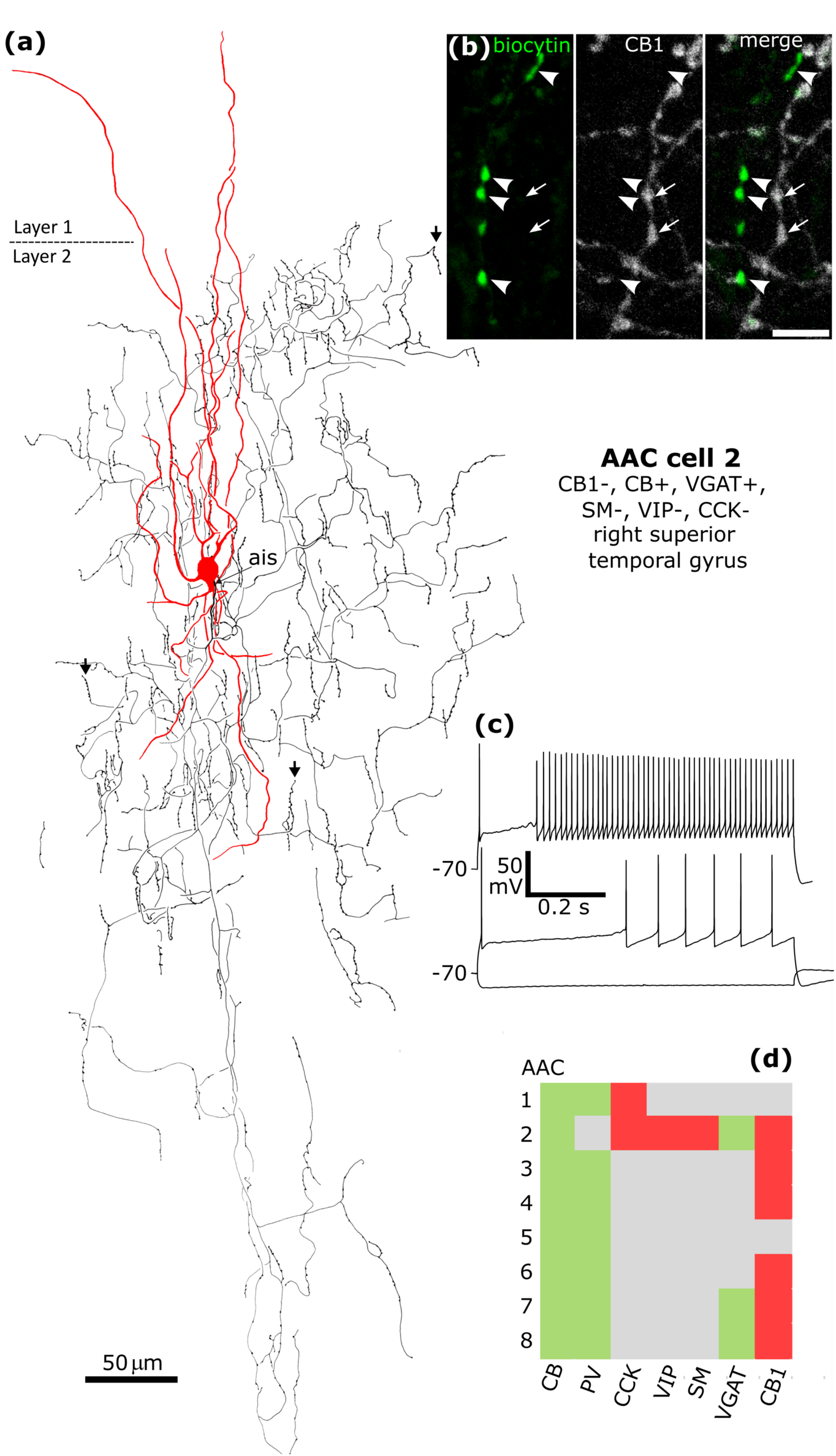
Axo-axonic cells are immunonegative for CB1 receptor. **(a**) Reconstruction of the recovered parts of an axo-axonic cell (No 2 in d) with soma and dendrites (red) and a descending axon (black) with characteristic radially oriented bouton cartridges (vertical arrows), each following the axon initial segment (ais) of a pyramidal cell. The ais originates from the base of the soma. **(b)** Biocytin labelled boutons (green) were immunonegative for CB1 (arrowheads) in an area of strongly CB1 positive axons (arrows). **(c)** Voltage responses of the cell to current injections of rheobase, RB +100 pA (upper trace), and holding current −100 pA; only a very slight voltage sag was detected to the hyperpolarising current injection. The cell fired an initial spike followed by a lag and fast spiking when continuously depolarised, which was characteristic of all eight axo-axonic cells recorded. **(d)** Immunoreactivities in all recorded and visualised axo-axonic cells (n=8) tested for 2-8 molecules: green, immunopositive; red, immunonegative; grey, not tested/inconclusive. All neurons tested were CB-, PV- and VGAT-positive, and none of the 6 cells tested showed CB1 immunoreactivity. Scale bars, (a), 50 μm; (b), 5 μm

Only 2 of 10 successfully tested CB1-positive cells were immunopositive for VGLUT3, one was a rosehip cell (Cell 1, Figure 8). Another cell had insufficient recovered axon for categorisation (Cell 18); this axon was mainly in layer 3 and was not similar to the axon of cell 1 or any basket cell axon. In addition to the 10 CB1-positive interneurons, three additional CB1-negative cells were also tested for VGLUT3 immunoreactivity of their boutons. One of these cells was VGLUT3-immunopositive (JR191213-5-IL2, not shown), located in layer 3 with bitufted dendrites and relatively dense axon around the cell body, with long axon collaterals in all directions also reaching into layer 1. The other two CB1- and VGLUT3-immunonegative cells were axo-axonic cells. Amongst the 10 VGLUT3-immunonegative cells, 8 were immunopositive for CB1 (Figure 8f), and included cells with descending axons (cells 14 and 15), basket cells (cells 11 and 13), a putative interneuron specific cell (cell 10), neurogliaform cells (cells 6 and 7) and one cell (cell 16) with insufficient recovered axon for identification.

## 4 DISCUSSION

The analysis of the origins, synaptic relationships and identities of CB1 receptor expressing nerve terminals in the human cerebral cortex has led to the following main conclusions:

1. Most GABAergic CB1 receptor expressing boutons make synapses with dendritic shafts and spines; neuronal somata are only around 10% of targets.
2. CB1 expressing GABAergic synaptic boutons originate from diverse presynaptic cell types expressing CCK and/or VIP and/or calretinin and/or COUP-TFII, such as basket cells, rosehip cells, interneuron specific cells, neurogliaform and other cell types.
3. A subpopulation of presumed GABAergic CB1 immunopositive nerve terminals also expresses VGLUT3.
4. VGLUT3 is also present in a subpopulation of VGLUT1 expressing glutamatergic nerve terminals which make type-1 synapses mainly with dendritic spines and are likely to be of intracortical origin.
5. VGLUT3 is also co-expressed in nerve terminals with VMAT2 and these boutons most likely originate from serotoninergic/glutamatergic neurons of the raphe nuclei.

### 4.1 Cell types providing CB1 expressing cortical terminals

The recreational use of cannabis derivatives produces intoxication, memory impairment, disruption of psychomotor behaviour, stimulation of appetite, as well as anti-emetic and antinociceptive actions in the central nervous system (see Iversen 2003). The potential harms and benefits of its medicinal use is under intense scrutiny (e.g. Schlag et al 2021). Cannabis use may also have profound effects on human embryonic development (Berghuis et al., 2007; Hurd et al., 2019). Some of these effects are produced via actions of cannabinoids on nerve terminals in cortical structures. Transcriptomic, immunocytochemical and physiological analyses show that CB1 receptors are expressed by glutamatergic pyramidal cells and GABAergic interneurons on their synaptic terminals as well as in some cortical afferents.

In the transcriptomic databases (http://celltypes.brain-map.org/rnaseq/, and Fig1) some somatostatin expressing GABAergic neuron groups also express a moderate level of CB1 transcript in the human cortex, but to our knowledge protein expression has not been confirmed. In the human MTG, which we have studied, amongst the 28 caudal ganglionic eminence derived ADARB2 expressing groups, 21 expresses CCK with high frequency and of these 14 show a high level of CB1 receptor expression; on the other hand, out of the 19 GABAergic groups that express high level of CB1 receptor, 13 have high levels of CCK expression. To what degree the variation in the level and expression of transcripts is due to sampling factors is not clear. Characterising the physiological and synaptic connectivity differences amongst these groups remains a major challenge in the human cortex (Lee et al., 2023; Varga et al., 2015).

The CB1-positive human cortical neurons we have identified correspond well to previously described GABAergic neurons in human and animal cortices (Boldog et al., 2018; Freund et al., 1986; Gonzalez-Albo et al., 2001; Kawaguchi and Kubota, 1998; Kubota and Kawaguchi, 1997; Kisvarday et al., 1990; Lee et al., 2023; Tsou et al 1999; Varga et al., 2015). In the human hippocampus most CCK expressing interneurons were found to be immunopositive for the CB1 receptor, whereas parvalbumin-positive neurons rarely showed immunoreactivity (Katona et al., 2000) similarly to the neocortex. Due to its laminar organisation, the hippocampus is an advantageous area for exploring cell type selectivity of structural and functional differentiation. In terms of CB1 receptor signalling, differences have been revealed between perisomatic innervating basket terminals, which are under tonic CB1 mediated inhibition and exclusively dendrite-innervating CB1/CCK expressing GABAergic neurons, whose GABA release may not be tonically regulated (Barti et al., 2024; Dudok et al., 2015; Lee et al., 2010; 2015). The two different CCK-expressing types of basket cells and the exclusively dendrite innervating GABAergic cell types were all shown to have CB1-positive axons (Ali 2007; Ali and Todorova 2010; Dudok et al., 2024; Klausberger et al., 2005, Koukouli et al., 2022; Lasztoczi 2011; Lee et al 2010). The latter interneuron types also innervate each other through functional CB1 receptor controlled GABAergic synapses (Ali, 2007; Ali and Todorova, 2010).

In our sample of human CB1-positive cortical cells, we distinguished 6 cell types based on their axonal patterns, enabling us to recognise some distinct rules. The soma innervating basket cells with descending axons are similar CCK-expressing interneurons in cat visual cortex (Freund et al., 1986) and rat frontal cortex (Kubota and Kawaguchi, 1997). Basket cells expressing CCK and CB1 often give translaminar axons also in mouse prefrontal cortex (Nagy-Pal et al., 2023), similar to the interneurons shown here in human association cortex. The suppression of GABA release from boutons of these neurons is likely to influence spiking of the postsynaptic neurons, as shown elegantly in mouse hippocampus during place cell firing (Dudok et al, 2024). In contrast, the CB1 positive rosehip and neurogliaform cells with spatially restricted axons would only influence local dendritic domains and the effectiveness of glutamatergic inputs to those domains. Some of these axons are restricted to layers I/II (Boldog et al., 2018; Chartrand et al., 2023; Field et al., 2023; Olah et al., 2007). In the neocortex glutamatergic innervation of the distal dendritic tufts of pyramidal cells in layer I is mainly from other cortical regions often in a feed-back manner. In contrast, the basal dendritic trees and apical dendritic oblique branches receive local, other cortical and thalamic glutamatergic inputs. The domain-specific glutamatergic input-dependent pairing of CB1/CCK GABAergic innervation of pyramidal cells is most clear in the hippocampus, where the two major glutamatergic inputs from the entorhinal cortex or CA3 pyramidal cells are associated with performant path-associated, or Schaffer collateral associated CB1/CCK/GABA expressing axonal interneurons, both in CA1 and CA3 (Ali, 2007; Klausberger et al., 2005; Lasztoczi et al., 2011;). This arrangement allows the selective CB1 receptor-dependent withdrawal of inhibition and local glutamatergic synaptic plasticity (Chevaleyre and Castillo 2004) upon either intracellular calcium increases (Boldog et al., 2015; Sugaya and Kano 2021) or the activation of mGluRs (Chevaleyre and Castillo, 2003) not only in the hippocampus but also in the neocortex involving the CB1 expressing dendrite innervating interneurons shown here.

We hypothesise that CB1-positive neurons immunoreactive for both VIP and calretinin are selectively interneuron innervating cells. In rat somatosensory cortex, calretinin-positive cell bodies are not immunoreactive for CB1 (Bodor et al., 2005) and VIP-positive neuronal cell bodies only rarely express CB1, indicating different organisation. Suppression of inhibition mediated by CB1 receptors was demonstrated in the hippocampus between interneurons (Ali, 2007; Ali end Todorova, 2010). We have found innervation of interneuron dendrites by CB1-positive synaptic terminals and visualised interneuron axons heavily innervating other interneuron somata. Synaptic suppression of GABA release between interneurons could lead to increased firing and inhibition of the postsynaptic principal cells (Ali and Todorova, 2010).

To our knowledge neurogliaform cells, which are thought to act by volume transmission of non-synaptic GABA release (Tamas et al., 2003), have not been shown to be controlled by axonal CB1 receptors. However, neurogliaform cells also form proper synaptic junctions with some of their boutons (Tamas et al., 2003; Olah et al., 2007; Fuentealba et al., 2010), which might be controlled by retrograde release of postsynaptic endocannabinoids. In addition, the dense axonal cloud of neurogliaform cells may also be under the influence of endocannabinoids during high intensity neuronal population activity. Neurogliaform axons can be spatially very restricted in the human cortex and even localised only to sublayers of layer I (Field et al., 2022; Chartrand et al., 2023). The suppression of their GABA release would be particularly suitable for providing opportunity for dendritic domain selective glutamatergic synaptic plasticity (see below).

Considering the diversity of CB1 expressing GABAergic cell groups, many more cell types remain to be discovered and defined in terms of their synaptic circuit organisation. There may also be difference between cortical areas in CB1 receptor mediated synaptic plasticity. For example, in the mouse primary visual cortex CB1expressing basket cells innervate local pyramidal cells and are not under tonic CB1 receptor control. In contrast, in the secondary visual cortex V2M they provide translaminar axons in which endocannabinoids tonically suppress GABA release (Koukouli et al., 2022) leading to different firing dynamics in the two visual areas.

Perisomatic innervating GABAergic cortical neurons include the axo-axonic cells (Somogyi et al., 1982; Dudok et al., 2021). Because of their strategically located GABAergic synapses exclusively on the axon initial segment, which is involved in synaptic plasticity, we tested such boutons for CB1 immunoreactivity and could not detect any. In the MTG and multiple area transcriptomic samples the CALB1 expressing cluster *Inh L1-6 PVALB SCUBE3* homologous to the mouse clusters of axo-axonic cells (Hodge et al., 2019) as well as the corresponding cluster in M1 (*Inh L1-6 PVALB COL15A1*) both show a low frequency of transcript for CB1, even if the protein may not be expressed, or was below our detection threshold. Similarly, several SST expressing GABAergic neuronal groups show moderate levels of transcript expression, but we are unaware of any demonstration of protein expression or functional tests for CB1 receptor.

### 4.2 Location of CB1 receptors - technical considerations

The membrane location of CB1 receptors relative to endocannabinoid sources and effectors is an important functional determinant. We have used antibodies to the cytoplasmic C-terminal domain of the CB1 receptor (Fukudome et al., 2004) with the sensitive but low resolution immunoperoxidase technique resulting in diffusion of the HRP reaction product within the neuronal processes. Super-resolution microscopic quantitative analysis of immunoreactive CB1 receptor distribution and abundance in single nerve terminals showed that on GABAergic hippocampal terminals CB1 receptors are not uniformly distributed and there appears to be no correlation between receptor abundance and the suppression of transmitter release from GABAergic terminals (Dudok et al., 2015; Lenkey et al., 2015). However, the key parameter that appears to determine the effectiveness of tonic inhibition of GABA release is the number of CB1 receptors in and around the presynaptic release site relative to the number of release sites in single boutons (Barti et al., 2024). Indeed, high resolution postembedding immunogold (Nyiri et al., 2005) and freeze fracture membrane replica immunogold analysis (Lenkey et al., 2015) of CB1 receptors on the presynaptic bouton surface relative to synaptic active zones shows that they are enriched in a peri-junctional annulus and also on preterminal axons (Nyiri et al., 2005). The receptors are largely absent from the presynaptic active zone. Pre-embedding silver-intensified immunogold reactions for electron microscopy also reveal a lack of immunoreactivity in the presynaptic active zones of both type-2 presumed GABAergic) and type-1 (presumed glutamatergic) terminals in the hippocampus and cerebellum (Bodor et al., 2005; Katona et al., 1999; 2006; Kawamura at el., 2006). However, with these techniques it cannot be completely excluded that in a chemically fixed tissue the cytoplasmic domain epitopes in the crowded active zone are masked by interacting protein(s) resulting in false negative immunoreactivity. Against this possibility is the consistent localisation of group III mGluRs specifically in the presynaptic active zone of both type-1 and type-2 cortical synaptic junctions (Dalezios et al., 2002; Ferraguti et al., 2005; Shigemoto et al., 1996; Somogyi et al., 2003). However, epitopes may be differently exposed on these proteins.

In presumed glutamatergic boutons making type-1 synapses we found denser CB1-immunoreactivity at the presynaptic active zone than in the rest of the boutons, implying that the effector(s) are distributed differently from those in GABAergic terminals. However, with the immunoperoxidase method it cannot be excluded that the actual antibody binding for CB1 was at the edge of the active zone in a peri-junctional ring and that the reaction product diffused onto the protein matrix of the active zone. We did not attempt to quantify this reaction, as the reaction strength depends on the depth of penetration of antibodies. Nevertheless, boutons making type-1 synapses consistently showed a lower level of immunoreactivity as compared to those making type-2 synapses, possibly reflecting a difference in CB1 receptor density between GABAergic and glutamatergic terminals.

Our immunohistochemical and recorded neuron samples derive from few cortical associational areas. There is evidence that the circuit organisation of CB1 receptor mediated synaptic plasticity of GABAergic synapses can differ in various cortical areas (Koukouli et al., 2022). Future studies comparing a wider range of distinct cortical areas in the human cortex could reveal specialisations. Amongst the GABAergic interneurons recorded in vitro, we consistently observed weaker CB1 immunoreactivity on their axons than on some of the nearby immunopositive axons of unknown cells. It is possible that the induction of repeated firing of the recorded neurons and/or the dialysis of the axons with the recording electrode solution reduced the receptors in the membrane by internalisation. Depending on the duration of the recording, this may have resulted in false immunonegativity in some of the tested neurons. Some of the single recorded neurons reported here were included in two previous studies (Field et al., 2021; Lukacs et al., 2023). Their action potential and firing characteristics were quantitatively reported, similar to other human cortical interneuron types (Chartrand et al., 2023; Lee et al., 2023). We did not attempt to compare the firing characteristics of CB1-positive and CB-negative neurons due to the large diversity of both groups and the relatively few CB1-positive cells in our sample, several of which had regular accommodating firing and a sag potential.

### 4.3 Endocannabinoid mediated synaptic plasticity

The predominant location of CB1-expressing GABAergic terminals on dendritic shafts and spines in human cortex is consistent with their role in synaptic plasticity. However, considering the importance of how recreational cannabinoids and those used as medication affect the cortex, relatively little is known about underlying molecular mechanisms in human cortical neurons. The activation of CB1 receptors on nerve terminals by endocannabinoids reduces presynaptic neurotransmitter release both tonically and phasically through several distinct mechanisms (see Barti et al., 2024; Sugaya and Kano, 2021). In addition, in some cortical neurons the activation of somato-dendritic potassium channels by endocannabinoids leading to self-inhibition was also reported (Bacci et al., 2004; Marinelli et al., 2008). Endocannabinoid retrograde signalling plays a major role in short and long-term synaptic plasticity (Kreitzer and Regehr, 2001; Ohno-Shoshaku et al., 2001), and has been studied extensively at cortical GABAergic synapses (Hajos et al., 2000; Wilson and Nicoll, 2001; Wilson et al., 2001; for review see Sugaya and Kano, 2021; Chevaleyre and Piskorowski 2014). Dendritic location of most CB1-positive GABAergic synapses in our samples, as well as in the monkey cortex (Eggan et al., 2010), is in line with the demonstration that CB1 receptor activation enhances backpropagating dendritic action potentials (Hsieh and Levine 2013) involved in synaptic plasticity. Furthermore, group I mGluR activation-dependent long-term depression of inhibition (I-LTD, Chevaleyre and Castillo, 2003) is also expected to be dendritically located. The group I mGluRs, mGluR1 and mGluR5 required for I-LTD are enriched in dendritic spine membrane, particularly at the edge of postsynaptic membrane specialisation (Baude et al.1993, Lujan et al., 1996), in a microdomain with phospholipase C-beta-1 and DAG-lipase (Fukaya et al., 2008; Katona et al., 2006), the two enzymes required for 2-AG generation. Chevaleyre and Castillo (2003) demonstrated in the hippocampus that heterosynaptic I-LTD, depends on postsynaptic group I mGluR activation resulting in PLC-mediated diacylglycerol (DAG) production. The DAG is converted by DAG-lipase to 2-AG released by postsynaptic pyramidal cells. Endocannabinoid-evoked I-LTD is mediated by CB1 receptor activation, inhibition of presynaptic adenylyl cyclase, leading to reduction of PKA signalling via RIM1alpha of the release machinery (Chevaleyre et al., 2007). This I-LTD is not blocked by buffering postsynaptic calcium, thus endocannabinoids can be released without a rise in postsynaptic calcium. It takes several minutes of CB1 activation to produce I-LTD, unlike depolarisation induced suppression of inhibition (DSI, Pitler and Alger, 2004; Yoshida et al., 2002), which is transient.

Human cortical neurons also show DSI (Kovacs et al., 2012). In contrast to I-LTD, endocannabinoid-mediated DSI or suppression of excitation (DSE) requires global activation of voltage gated calcium channels in neurons (Kreitzer and Regehr 2001; Wilson et al., 2001; but see Diana and Marty 2004) leading to a rise of intracellular calcium and release of endocannabinoids on a time-scale of seconds. It does not require mGluR activation, or inhibition of presynaptic cAMP/PKA signalling (Chevaleyre et al., 2006). In the hippocampus, DSI is mediated by the CB1 receptor activation of G_β/γ_ acting directly on N-type voltage-gated calcium channels (Hefft and Jonas 2005; Wilson et al., 2001), which are present in the terminals of CB1/CCK expressing GABAergic interneurons.

The functional consequences of short and long-term suppression of inhibition of pyramidal cells are not only increased excitability of the affected neurons, but also long-term increase of the efficacy of glutamatergic synapses (Chevaleyre and Castillo, 2004), i.e. LTP and consequent excitation-spike (E-S) coupling (Bliss and Lomo 1973), which depends on disinhibition. For increased E-S coupling both mGluR and CB1 receptor activation are required (Chevaleyre and Castillo, 2003) resulting in cannabinoid-mediated I-LTD, which is likely to contribute to learning. Chevaleyre and Castillo (2004) suggested that in the hippocampus I-LTD results in metaplasticity of glutamatergic inputs localised to distinct parts of dendritic trees. This idea is supported by our demonstration of CB1 receptor expressing GABAergic neurons that have spatially restricted axonal overlap with dendritic trees, such as neurogliaform and rosehip cells.

The recreational use of cannabis products disrupts cognitive processes including various forms of memory (Lichtman et al., 2002; Sullivan, 2000), most likely linked to alteration of synaptic plasticity mechanisms. In mice, a single moderate dose of delta9-tetrahydrocannabinol, the main psychoactive component of cannabis, abolished endocannabinoid mediated LTD of glutamatergic synaptic transmission in the nucleus accumbens and I-LTD in the hippocampus (Mato et al., 2004). Similar treatment also reduced CB1 receptor abundance on GABAergic terminals (Dudok et al., 2015) in the hippocampus for several days.

### 4.4 Neuronal network state-related physiological activity of CB1/CCK expressing cortical GABAergic neurons

The physiological role of CB1 receptor expressing GABAergic interneurons, which are mostly, but not exclusively, the CCK expressing groups, is most likely related to retrograde endocannabinoid signalling mediated synaptic plasticity, which requires knowledge of the spike timing of pre-and postsynaptic neurons. The expected evolutionary differentiation of spatio-temporal patterns amongst CCK/CB1 expressing vs other GABAergic neurons is clearly revealed in the rodent hippocampus, where the strict laminar differentiation of at least 5 CCK-expressing neuronal types in the CA1 area alone (Somogyi and Klausberger, 2015) have been extensively studied in relation to specific behavioural state dependent physiological network activity (Dudok et al., 2024; Farrell et al., 2021; Forro et al., 2023; Klausberger et al., 2005; Lasztoczi et al., 2011). The endocannabinoid mediated suppression of inhibition during movement-related 5-12 Hz theta network oscillations was suggested to contribute to place cell firing (Freund et al., 2003). The demonstration of the in vivo firing patterns of CCK/CB1 expressing GABAergic neurons revealed that during theta activity they fire preferentially at the ascending phase of individual 80-200 ms duration theta cycles in contrast to other GABAergic neurons (Klausberger et al. 2005; Lasztoczi et al, 2011). In another brain state, during the 30-120 ms duration sharp wave associated ripple oscillation events, CCK/CB1 expressing GABAergic cells reduce their activity, but often show post-ripple rebound, in contrast to some parvalbumin-expressing GABAergic neuronal types innervating the same pyramidal neurons. Thus, retrograde endocannabinoid-evoked reduction of inhibition selectively to active pyramidal cells is likely to operate at the above specific time windows. Hippocampal pyramidal place cells fire most intensely in the place field, starting around the theta phase when CB1/CCK cells fire, as the animal enters the place field. The firing evoked endocannabinoid release reducing inhibition selectively by CCK/CB1 expressing presynaptic neurons to the active pyramidal cell, leads to acceleration of firing lifting the active cells out of the population that did not reach firing threshold and remain suppressed by CB1/CCK interneurons (Klausberger et al., 2005). Indeed, this has now been elegantly demonstrated by simultaneously recording optically both postsynaptic pyramidal cell firing and endocannabinoid levels in mice running on a treadmill (Dudok et al., 2024). In place cells that fire complex spikes in specific location, firing increased intracellular calcium and led to the synthesis of 2-AG (Farrell et al., 2021) resulting in the suppression of presynaptic inhibition specifically from presynaptic CCK basket cell terminals (Dudok et al., 2024). Further experiments showed that this signalling contributes to the specificity of place field firing activity. Such a mechanism of increasing contrast in firing frequency between pyramidal cells that reached firing threshold and those that did not, through disinhibition, may be a general role of CB1 expressing cortical GABAergic neurons in all cortical areas.

The somatic innervation of the same pyramidal cells by distinct parvalbumin or CCK/CB1 expressing basket cells has provided fertile ground for experimentation leading to many suggestions of the physiological significance of this duality (Allene et al., 2015; Armstrong and Soltesz 2012; Bodor et al., 2005; Forro et al., 2023; Glickfeld and Scanziani, 2006; Klausberger et al., 2003, 2005; Nagy-Pal et al., 2023). It is possible that PV and CCK may be co-expressed in some mouse but not rat cortical interneurons (Grieco et al., 2023), which remain to be characterised. Much less attention has been paid to the similar dual innervation by CCK/CB1 or parvalbumin expressing interneurons of the same dendritic domains (Forro et al., 2023; Klausberger et al., 2004; 2005). With regard to somatic innervation, genetically engineered mice with highly specific genetically labelled CCK-expressing basket cells in the hippocampus of mice, Dudok et al. (2021, 2024) elegantly demonstrated that the CCK basket cells whose axons are endowed by CB1 cannabinoid receptors strongly reduce their firing during running, in contrast to parvalbumin expressing basket cells which strongly increase their firing (see also Forro et al., 2023). The CCK basket cells increased firing when the mice stopped running (Dudok et al. 2021; Forro et al., 2023). Remarkably, the contrasting firing activity of the two types of basket cells was also evident in the completely different behavioural and brain state when the mice sat quietly and sharp wave associated ripple oscillations dominated the network. In this state, CCK expressing basket cells fire at a higher level on average than parvalbumin expressing ones. As to the mechanism of this division of labour, Dudok et al. (2021) showed that inputs from PV interneurons inhibit CCK/CB1 basket cells, which also inversely scales activity relative to all place cells and PV cells on long time scales. The firing of CCK cells negatively correlated with pyramidal cell ensemble activity, which is an additional mechanism for temporal specificity of GABA release in addition to the endocannabinoid/CB1 receptor-mediated suppression of GABA release to physiologically activated single pyramidal neurons.

The location and concentration of CB1 receptors in cortical neurons is also relevant to pathological events such as seizures and stroke. The development of an ingenious genetically encoded endocannabinoid imaging sensor that tracks the level of 2-AG during physiological and pathological neuronal activity in the brain revealed dynamic changes in 2-AG, but not in anandamide concentration, both at the network level and with single cell and single axonal bouton resolution (Dong et al., 2020; Farrell et al., 2021) in the hippocampus and amygdala. Individual pyramidal cells showed place-cell firing related increases in intracellular calcium concentration that positively correlates with 2-AG concentration. Furthermore, kindling-induced seizures increase the endocannabinoid 2-AG concentration to several fold of physiological levels which returns to normal within seconds (Farrell et al., 2021). The pharmacological inhibition of the synthesis of 2-AG reduces its level and prolongs seizures, demonstrating that 2-AG is involved in seizure control. Increase in 2-AG production leads to elevated vasoactive prostaglandin synthesis via the hydrolysis of 2-AG by monoacylglycerol lipase (MAGL) and consequent hypoxia that is prevented by a 2-AG synthesis inhibitor (Farrell et al., 2021). What effects these changes in endocannabinoid levels during seizures have on CB1-expressing GABAergic terminals remains unknown. However, one possibility is that extremely high levels of endocannabinoids may induce internalisation of CB1 receptors resulting in reduced presynaptic inhibition of GABA release in effect increasing inhibition, thus contributing to seizure control.

Furthermore, CB1 receptors on CCK-basket cell terminals in the prefrontal cortex have been shown to regulate social behaviour. Specifically, chronic social isolation stress (CSIS) has been shown to disrupt CB1 receptor mediated regulation of GABAergic synaptic transmission from CCK expressing cells leading to increased inhibition of principal cells in the anterior cingulate cortex (ACC), which resulted in the impairment of social behaviour (Guo et al., 2024). Importantly, CSIS-induced social impairments could be rescued by GABAergic CCK cell-specific optogenetic activation of CB1 receptors in the ACC (Guo et al., 2024).

### 4.5 VGLUT3 in human cortical nerve terminals

Neurons expressing VGLUT3 are involved in psychiatric and neurological disorders such as epilepsy, Alzheimer disease, Huntington chorea, Tourette syndrome and Parkinson disease, difficulties in stress coping, deafness and other disorders (Favier et al., 2021). For example, a human VGLUT3-pT8I mutation predisposes to substance abuse and eating disorders (Sakae et al., 2015) and is related to mis-regulation of ACh and glutamate synergy in synaptic vesicles resulting in altered striatal acetylcholine and dopamine release (Favier et al., 2024). Here, we have demonstrated the presence of VGLUT3 in the human neocortex in a subset of GABAergic cortical terminals also described in rodents (Fasano et al., 2017; Omiya et al., 2015; Pelkey et al., 2020; Somogyi et al., 2004), which most likely derive from local interneurons. Indeed, evidence was obtained by recording and labelling two VGLUT3 positive human interneurons *in vitro*. In the hippocampus and neocortex, as well as in the basolateral amygdala and entorhinal cortex VGLUT3 is expressed in a subset of CCK-expressing interneurons (Fasano et al., 2017; Omiya et al., 2015; Somogyi et al., 2004). One such cell type is a basket cell that is different from VIP-co-expressing CCK-positive basket cells, at least in the hippocampus (Somogyi et al, 2004) and in the basolateral amygdala (Omiya et al., 2015). As in the mouse basolateral amygdala, entorhinal cortex (Omiya et al., 2025) and hippocampus (Fasano et al., 2017), the VGLUT3-containing GABAergic terminals are also positive for CB1R in the human neocortex. Furthermore, Omiya et al. (2015) described that terminals are selectively apposed to DGL-alpha clumps in the postsynaptic somatic membrane of principal cells synthesizing the endocannabinoid 2-AG and invaginate into the postsynaptic soma, increasing the CB1 loaded presynaptic membrane surface and decreasing diffusion away from the presynaptic terminal. Such DGL-alpha clumps are specific to certain principal cells receiving GABA/VGLUT3/CCK somatic innervation. Furthermore, they called attention to potential cooperation of postsynaptic depolarisation evoked calcium entry and G_i/q_-coupled mGluR5 and CCK-2 receptor activation at these synapses by the release of a cocktail of glutamate and CCK. Although, we have not detected invaginated VGLUT3 containing GABAergic nerve terminals in the examined areas of the human cortex, such terminals were also reported in the rat hippocampus (Klausberger et al., 2005). It is possible that in some cortical structures such as the basolateral amygdala and the hippocampus, the perisomatic innervation of principal cells by CB1-expressing GABA/CCK/VGLUT3 terminals is more highly developed than in associational isocortical areas that we studied in the human cortex, where the weight of such innervation has shifted to the more distal dendrites. However, more homologous cortical structures and different cortical layers need to be compared to test this assumption.

The functional significance of the co-release of GABA and glutamate was tested in the rodent hippocampus (Fasano et al., 2017; Pelkey et al., 2020). By removing VGLUT3 selectively from GABAergic neurons Fasano et al., (2017) showed that the frequency of IPSCs increased in postsynaptic pyramidal cells, most likely due to a lack of glutamate release and the absence of group III presynaptic mGluR activation on GABAergic terminals. As reported in cholinergic interneurons and on 5-HT neurons, VGLUT3 may also enhance GABA loading into synaptic vesicles (Fasano et al., 2017; but see Pelkey et al., 2020). The absence of VGLUT3 in GABAergic terminals also altered theta frequency network oscillations and synaptic plasticity. When the synthesis of GABA was compromised the glutamatergic phenotype of the VGLUT3/CCK interneurons promoted hyperexcitability (Pelkey et al., 2020). The testing of endocannabinoid signalling at VGLUT3/GABA/CCK synapses which express CB1 receptors on pyramidal cells in the mouse hippocampus showed that they can undergo DSI (Pelkey et al., 2020). CB1 receptors on these terminals are likely to suppress the release of both GABA and glutamate, which at the human dendritic synapses could relieve mGluR5-mediated DGL-alpha activation in a feedback manner. The released glutamate may also act on postsynaptic ionotropic glutamate receptors (Pelkey et al., 2020), although Omiya et al. (2015) did not detect AMPA-type receptors at CCK/GABA/CB1 synapses. However, Szabadits et al. (2011) showed the presence of NMDA-type glutamate receptors in hippocampal somatic synaptic junctions which are likely to face presynaptic GABAergic terminals. Overall, the combined co-release of GABA and glutamate from local interneurons regulated by CB1 receptors likely to participate in fine-tuning excitability in the human cortex.

Surprisingly, 60% of VGLUT3 immunopositive boutons made type-1, presumably purely glutamatergic synaptic junctions. This proportion is similar to the 52% of VGLUT3-positive boutons which also expressed VGLUT1. Type-1 synaptic junctions have an extensive postsynaptic density enriched in PSD-95 amongst other proteins specific to glutamatergic synapses. Such synapses made by VGLUT3-positive boutons of unknown origin were also reported in the rodent hippocampus and dorsal raphe nucleus (Gras et al., 2002). The electron microscopically detected boutons most likely correspond to the terminals which we co-labelled for both VGLUT3 and VGLUT1, the latter being mainly present in glutamatergic terminals of cortical origin. The transcriptomic screen revealed a very restricted and low frequency occurrence only in one group of VGLUT1 expressing cortical neurons at least in the reported cortical areas, namely the MTG and in the samples from multiple cortical areas that also included the MTG. In situ hybridization for VGLUT3 in rodent neocortex also showed some signal in possible deep layer pyramidal cells in frontal cortex (unpublished results, El Mestikawy), but they were not tested for VGLUT1 expression. It would be surprising if one deep layer infrequent glutamatergic cell group in our samples provided the frequent VGLUT3-positive type-1 synapses in the supragranular layers, but this cannot be excluded. Another possibility is that some isocortical areas that have not been identified yet are endowed by a pyramidal cell population that provides widespread VGLUT3/VGLUT1 innervation to the rest of the neocortex in humans, although a study of several cortical areas has not identified message in pyramidal cells (Vigneault et al., 2015). The dual expression of vesicular transporters may endow the vesicles with special release mechanisms, which may also be related to unusual synaptic plasticity rules on the spines that we identified as the main postsynaptic targets. Due to the lack of access to suitable antibodies we were not able to test the potential co-expression of VGLUT2 and VGLUT3 in cortical terminals. In the mouse thalamus there is weak expression of slc17A8 transcripts for VGLUT3 in neurons of the midline nuclei (Allen Institute, https://mouse.brain-map.org), which also express VGLUT2, but it is not known if these cells project to the cortex, or if the homologous human thalamic nuclei express slc17A8.

We have also confirmed the co-localisation of VMAT2 and VGLUT3 in the human cortex. Based on rodent studies, these boutons are likely to originate from serotoninergic neurons of the raphe nuclei (Gras et al., 2002; Fremeau et al., 2002, Herzog et al., 2004) which expresses VGLUT3 also in humans (Vigneault et al., 2015). The role of VGLUT3 in serotoninergic terminals has been suggested to provide a glutamatergic neurotransmitter phenotype (Varga et al., 2009). Indeed, the hippocampus, the medial septum and the prefrontal cortex are innervated by median raphe serotonergic/glutamatergic terminals which activate postsynaptic ionotropic glutamate receptors Varga et al 2009; Szonyi et al., 2016).

## ABBREVIATIONS

2-AG: 2-arachidonoylglycerol
5-HT: serotonin
AAC: axo-axonic cell
ACC: anterior cingulate cortex
BC: basket cell
cAMP/PKA: cyclic adenosine monophosphate/protein kinase A
CALB1: gene for calbindin
CB1: cannabinoid receptor type 1
CCK: cholecystokinin
COUP-TF2: Chicken ovalbumin upstream promoter transcription factor II
CSIS: chronic social isolation stress
DAC: cell with radial descending axons
DAG: diacylglycerol
DAGLα: diacylglycerol-lipase-alpha
DSE: depolarisation induced suppression of excitation
DSI: depolarisation induced suppression of inhibition
DTC: dendrite targeting cell
E-S: excitation-spike coupling
eIPSPs: evoked monosynaptic inhibitory postsynaptic potentials
GAD67: glutamate decarboxylase, 67 kDalton
GABA: gamma aminobutyric acid
HRP: horseradish peroxidase
II-LTD: long-term depression of inhibition
ISC: interneuron specific cell
LAC: loose axon cell
LTD: long-term depression
LTP: long-term potentiation
M1: primary motor cortex
MAGL: monoacylglycerol lipase
M1: 3 muscarinic acetylcholine receptors
MTG: middle temporal gyrus
PB: phosphate buffer
PSD-95: postsynaptic density protein 95kDalton
PVALB: gene for parvalbumin
RB: rheobase RH, rosehip cell
SE: standard error of the mean
SM: somatostatin
SST: gene for somatostatin
STD: short-term depression
VGLUT1: vesicular-glutamate-transporter-1
VGLUT2: vesicular-glutamate-transporter-2
VGLUT3: vesicular-glutamate-transporter-3
VIP: vasoactive intestinal polypeptide
VMAT2: vesicular monoamine transporter-2

## ACKNWOLEDGEMENTS

The authors thank Zalan Ilyes for the reconstruction of the axo-axonic cell in figure 15, Martin Field for contributing labelled neurons in layer 1, Ruggiero Francavilla for contributing axo-axonic cells that he recorded for immunohistochemical testing here and Jozsef Somogyi for advice on confocal microscopy. We thank Istvan Katona for helpful comments on a previous version of the manuscript. The study was supported by the European Research Council (ERC-2015-AdG 694988 to P.S.); the Oxford National Institute for Health Research Biomedical Research Centre (MC_UU_12024/4 to P.S.); the Medical Research Council (MR/R011567/1 to P.S.), Nuffield Benefaction for Medicine and the Wellcome Institutional Strategic Support Fund (grant 0009985, to T.J.V.). Istvan Lukacs’ doctoral scholarship was supported by the Medical Sciences Division of the University of Oxford and the Dulverton Trust.

## AUTHOR CONTRIBUTIONS

**Peter Somogyi**: designed research, obtained and analysed data, writing and editing draft, funding acquisition; **Sawa Horie**: designed research, obtained and analysed data, editing draft; **Istvan Lukacs**, designed research, obtained and analysed data, editing draft; **Emily Hunter**: designed research, obtained and analysed data, editing draft; **Barbara Sarkany**: analysed data, editing draft; **Tim Viney**, designed research, obtained and analysed data, editing draft, funding acquisition; **James Livermore**: contributing materials, editing draft; **Puneet Plaha**: contributing materials, editing draft; **Richard Stacey**, contributing materials, editing draft; **Olaf Ansorge**, contributing materials, editing draft; **Salah El Mestikawy**: contributing tools, editing draft; **Quianru Zhao**: designed research, obtained and analysed data, editing draft.

## CONFLICT OF INTEREST STATEMENT

The authors declare no competing interests.

## DATA AVAILABILITY

Peter Somogyi

## ETHICS APPROVAL AND PATIENT CONSENT

Human tissue samples from neurosurgery at the John Radcliffe Hospital (Oxford) for the treatment of brain tumours or temporal lobe epilepsy (table 1) were collected in accordance with the Human tissue Act 2004 (UK), under the licence (15/SC/0639) of the Oxford Brain Bank, John Radcliffe Hospital, Oxford, UK. Fully informed patients formally consented to providing samples, which were access tissue that were removed in order to access the diseased part of the brain

## Notes

### Competing Interest Statement

The authors have declared no competing interest.

## RERERENCES

Airaksinen, M.S., Eilers, J., Garaschuk, O., Thoenen, H., Konnerth, A. & Meyer, M. (1997) Ataxia and altered dendritic calcium signaling in mice carrying a targeted null mutation of the calbindin D28k gene. Proc Nat Acad Sci USA, 94, 1488–1493.

Ali, A.B. (2007) Presynaptic inhibition of GABA_A_ receptor-mediated unitary IPSPs by cannabinoid receptors at synapses between CCK-positive interneurons in rat hippocampus. J Neurophysiol, 98, 861–869.

Ali, A.B. & Todorova, M. (2010) Asynchronous release of GABA via tonic cannabinoid receptor activation at identified interneuron synapses in rat CA1. Eur J Neurosci, 31, 1196–1207.

Allene, C., Lourenco, J. & Bacci, A. (2015) The neuronal identity bias behind neocortical GABAergic plasticity. Trends Neurosci, 38, 524–534.

Armstrong, C. & Soltesz, I. (2012) Basket cell dichotomy in microcircuit function. J Physiol, 590, 683–694.

Bacci, A., Huguenard, J.R. & Prince, D.A. (2004) Long-lasting self-inhibition of neocortical interneurons mediated by endocannabinoids. Nature, 431, 312–316.

Bakken, T.E. & Jorstad, N.L. & Hu, Q. & Lake, B.B. & Tian, W. & Kalmbach, B.E. & Crow, M. & Hodge, R.D. & Krienen, F.M. & Sorensen, S.A. & Eggermont, J. & Yao, Z. & Aevermann, B.D. & Aldridge, A.I. & Bartlett, A. & Bertagnolli, D. & Casper, T. & Castanon, R.G. & Crichton, K. & Daigle, T.L. & Dalley, R. & Dee, N. & Dembrow, N. & Diep, D. & Ding, S.L. & Dong, W. & Fang, R. & Fischer, S. & Goldman, M. & Goldy, J. & Graybuck, L.T. & Herb, B.R. & Hou, X. & Kancherla, J. & Kroll, M. & Lathia, K. & van Lew, B. & Li, Y.E. & Liu, C.S. & Liu, H. & Lucero, J.D. & Mahurkar, A. & McMillen, D. & Miller, J.A. & Moussa, M. & Nery, J.R. & Nicovich, P.R. & Niu, S.Y. & Orvis, J. & Osteen, J.K. & Owen, S. & Palmer, C.R. & Pham, T. & Plongthongkum, N. & Poirion, O. & Reed, N.M. & Rimorin, C. & Rivkin, A. & Romanow, W.J. & Sedeno-Cortes, A.E. & Siletti, K. & Somasundaram, S. & Sulc, J. & Tieu, M. & Torkelson, A. & Tung, H. & Wang, X. & Xie, F. & Yanny, A.M. & Zhang, R. & Ament, S.A. & Behrens, M.M. & Bravo, H.C. & Chun, J. & Dobin, A. & Gillis, J. & Hertzano, R. & Hof, P.R. & Hollt, T. & Horwitz, G.D. & Keene, C.D. & Kharchenko, P.V. & Ko, A.L. & Lelieveldt, B.P. & Luo, C. & Mukamel, E.A. & Pinto-Duarte, A. & Preissl, S. & Regev, A. & Ren, B. & Scheuermann, R.H. & Smith, K. & Spain, W.J. & White, O.R. & Koch, C. & Hawrylycz, M. & Tasic, B. & Macosko, E.Z. & McCarroll, S.A. & Ting, J.T. & Zeng, H. & Zhang, K. & Feng, G. & Ecker, J.R. & Linnarsson, S. & Lein, E.S. (2021) Comparative cellular analysis of motor cortex in human, marmoset and mouse. Nature, 598, 111–119.

Barti, B., Dudok, B., Kenesei, K., Zöldi, M., Miczán, V., Balla, G.Y., Zala, D., Tasso, M., Sagheddu, C., Kisfali, M., Tóth, B., Ledri, M., Vizi, E.S., Melis, M., Barna, L., Lenkei, Z., Soltész, I. & Katona, I. (2024) Presynaptic nanoscale components of retrograde synaptic signaling. Sci Adv, 10, eado0077.

Baude, A., Nusser, Z., Roberts, J.D.B., Mulvihill, E., McIlhinney, R.A.J. & Somogyi, P. (1993) The metabotropic glutamate receptor (mGluR1a) is concentrated at perisynaptic membrane of neuronal subpopulations as detected by immunogold reaction. Neuron, 11, 771–787.

Berghuis, P., Rajnicek, A.M., Morozov, Y.M., Ross, R.A., Mulder, J., Urbán, G.M., Monory, K., Marsicano, G., Matteoli, M., Canty, A., Irving, A.J., Katona, I., Yanagawa, Y., Rakic, P., Lutz, B., Mackie, K. & Harkany, T. (2007) Hardwiring the brain: endocannabinoids shape neuronal connectivity. Science, 316, 1212–1216.

Bernal-Chico, A., Tepavcevic, V., Manterola, A., Utrilla, C., Matute, C. & Mato, S. (2023) Endocannabinoid signaling in brain diseases: Emerging relevance of glial cells. Glia, 71, 103–126.

Bloss, E.B., Cembrowski, M.S., Karsh, B., Colonell, J., Fetter, R.D. & Spruston, N. (2016) Structured Dendritic Inhibition Supports Branch-Selective Integration in CA1 Pyramidal Cells. Neuron, 89, 1016–1030.

Bocchio, M., Lukacs, I., Stacey, R., Plaha, P., Apostolopoulos, V., Livermore, L., Argue, S., Ansorge, O., Gillies, M.J., Somogyi, P. & Capogna, M. (2019) Group II metabotropic glutamate receptors mediate presynamptic inhibition of excitatory transmission in pyramidal neurons of the human cerebral cortex. Frontiers in Cellular Neuroscience, 12, 508.

Bodor, A.L., Katona, I., Nyíri, G., Mackie, K., Ledent, C., Hájos, N. & Freund, T.F. (2005) Endocannabinoid signaling in rat somatosensory cortex: laminar differences and involvement of specific interneuron types. J Neurosci, 25, 6845–6856.

Boldog, E., Bakken, T.E., Hodge, R.D., Novotny, M., Aevermann, B.D., Baka, J., Borde, S., Close, J.L., Diez-Fuertes, F., Ding, S.L., Farago, N., Kocsis, A.K., Kovacs, B., Maltzer, Z., McCorrison, J.M., Miller, J.A., Molnar, G., Olah, G., Ozsvar, A., Rozsa, M., Shehata, S.I., Smith, K.A., Sunkin, S.M., Tran, D.N., Venepally, P., Wall, A., Puskas, L.G., Barzo, P., Steemers, F.J., Schork, N.J., Scheuermann, R.H., Lasken, R.S., Lein, E.S. & Tamas, G. (2018) Transcriptomic and morphophysiological evidence for a specialized human cortical GABAergic cell type. Nat Neurosci, 21, 1185–1195.

Booker, S.A., Althof, D., Gross, A., Loreth, D., Müller, J., Unger, A., Fakler, B., Varro, A., Watanabe, M., Gassmann, M., Bettler, B., Shigemoto, R., Vida, I. & Kulik, Á. (2017) KCTD12 Auxiliary Proteins Modulate Kinetics of GABAB Receptor-Mediated Inhibition in Cholecystokinin-Containing Interneurons. Cereb Cortex, 27, 2318–2334.

Buhl, E.H., Halasy, K. & Somogyi, P. (1994) Diverse sources of hippocampal unitary inhibitory postsynaptic potentials and the number of synaptic release sites. Nature, 368, 823–828.

Buhl, E.H., Tamas, G. & Somogyi, P. (1995) Recurrent unitary EPSPs evoked in anatomically identified inhibitory interneurones of the cat visual cortex in vitro. J Physiol (Lond), 487, 51P.

Chartrand, T., Dalley, R., Close, J., Goriounova, N.A., Lee, B.R., Mann, R., Miller, J.A., Molnar, G., Mukora, A., Alfiler, L., Baker, K., Bakken, T.E., Berg, J., Bertagnolli, D., Braun, T., Brouner, K., Casper, T., Csajbok, E.A., Dee, N., Egdorf, T., Enstrom, R., Galakhova, A.A., Gary, A., Gelfand, E., Goldy, J., Hadley, K., Heistek, T.S., Hill, D., Jorstad, N., Kim, L., Kocsis, A.K., Kruse, L., Kunst, M., Leon, G., Long, B., Mallory, M., McGraw, M., McMillen, D., Melief, E.J., Mihut, N., Ng, L., Nyhus, J., Oláh, G., Ozsvár, A., Omstead, V., Peterfi, Z., Pom, A., Potekhina, L., Rajanbabu, R., Rozsa, M., Ruiz, A., Sandle, J., Sunkin, S.M., Szots, I., Tieu, M., Toth, M., Trinh, J., Vargas, S., Vumbaco, D., Williams, G., Wilson, J., Yao, Z., Barzo, P., Cobbs, C., Ellenbogen, R.G., Esposito, L., Ferreira, M., Gouwens, N.W., Grannan, B., Gwinn, R.P., Hauptman, J.S., Jarsky, T., Keene, C.D., Ko, A.L., Koch, C., Ojemann, J.G., Patel, A., Ruzevick, J., Silbergeld, D.L., Smith, K., Sorensen, S.A., Tasic, B., Ting, J.T., Waters, J., de Kock, C.P.J., Mansvelder, H.D., Tamas, G., Zeng, H., Kalmbach, B. & Lein, E.S. (2023) Morphoelectric and transcriptomic divergence of the layer 1 interneuron repertoire in human versus mouse neocortex. Science, 382, eadf0805.

Chevaleyre, V. & Castillo, P.E. (2003) Heterosynaptic LTD of hippocampal GABAergic synapses: A novel role of endocannabinoids in regulating excitability. Neuron, 38, 461–472.

Chevaleyre, V. & Castillo, P.E. (2004) Endocannabinoid-mediated metaplasticity in the hippocampus. Neuron, 43, 871–881.

Chevaleyre, V., Heifets, B.D., Kaeser, P.S., Südhof, T.C. & Castillo, P.E. (2007) Endocannabinoid-mediated long-term plasticity requires cAMP/PKA signaling and RIM1alpha. Neuron, 54, 801–812.

Chevaleyre, V. & Piskorowski, R. (2014) Modulating excitation through plasticity at inhibitory synapses. Front Cell Neurosci, 8, 93.

Chevaleyre, V., Takahashi, K.A. & Castillo, P.E. (2006) Endocannabinoid-mediated synaptic plasticity in the CNS. Annu Rev Neurosci, 29, 37–76.

Chou, S., Ranganath, T., Fish, K.N., Lewis, D.A. & Sweet, R.A. (2022) Cell type specific cannabinoid CB1 receptor distribution across the human and non-human primate cortex. Sci Rep, 12, 9605.

Dalezios, Y., Lujan, R., Shigemoto, R., Roberts, J.D.B. & Somogyi, P. (2002) Enrichment of mGluR7a in the presynaptic active zones of GABAergic and non-GABAergic terminals on interneurons in the rat somatosensory cortex. Cereb Cortex, 12, 961–974.

De Giacomo, V., Ruehle, S., Lutz, B., Häring, M. & Remmers, F. (2022) Cell type-specific genetic reconstitution of CB1 receptor subsets to assess their role in exploratory behaviour, sociability, and memory. Eur J Neurosci, 55, 939–951.

De Petrocellis, L., Nabissi, M., Santoni, G. & Ligresti, A. (2017) Actions and Regulation of Ionotropic Cannabinoid Receptors. Adv Pharmacol, 80, 249–289.

Del Pino, I., Brotons-Mas, J.R., Marques-Smith, A., Marighetto, A., Frick, A., Marín, O. & Rico, B. (2017) Abnormal wiring of CCK(+) basket cells disrupts spatial information coding. Nat Neurosci, 20, 784–792.

del Rio, M.R. & Defelipe, J. (1995) A light and electron microscopic study of calbindin D-28k immunoreactive double bouquet cells in the human temporal cortex. Brain Res, 690, 133–140.

Diana, M.A. & Marty, A. (2004) Endocannabinoid-mediated short-term synaptic plasticity: depolarization-induced suppression of inhibition (DSI) and depolarization-induced suppression of excitation (DSE). Br J Pharmacol, 142, 9–19.

Dong, A., He, K., Dudok, B., Farrell, J.S., Guan, W., Liput, D.J., Puhl, H.L., Cai, R., Wang, H., Duan, J., Albarran, E., Ding, J., Lovinger, D.M., Li, B., Soltesz, I. & Li, Y. (2022) A fluorescent sensor for spatiotemporally resolved imaging of endocannabinoid dynamics in vivo. Nat Biotechnol, 40, 787–798.

Dudok, B., Barna, L., Ledri, M., Szabo, S.I., Szabadits, E., Pinter, B., Woodhams, S.G., Henstridge, C.M., Balla, G.Y., Nyilas, R., Varga, C., Lee, S.H., Matolcsi, M., Cervenak, J., Kacskovics, I., Watanabe, M., Sagheddu, C., Melis, M., Pistis, M., Soltesz, I. & Katona, I. (2015) Cell-specific STORM super-resolution imaging reveals nanoscale organization of cannabinoid signaling. Nat Neurosci, 18, 75–86.

Dudok, B., Fan, L.Z., Farrell, J.S., Malhotra, S., Homidan, J., Kim, D.K., Wenardy, C., Ramakrishnan, C., Li, Y., Deisseroth, K. & Soltesz, I. (2024) Retrograde endocannabinoid signaling at inhibitory synapses in vivo. Science, 383, 967–970.

Dudok, B., Klein, P.M., Hwaun, E., Lee, B.R., Yao, Z., Fong, O., Bowler, J.C., Terada, S., Sparks, F.T., Szabo, G.G., Farrell, J.S., Berg, J., Daigle, T.L., Tasic, B., Dimidschstein, J., Fishell, G., Losonczy, A., Zeng, H. & Soltesz, I. (2021) Alternating sources of perisomatic inhibition during behavior. Neuron, 109, 997–1012

Dudok, B., Klein, P.M. & Soltesz, I. (2022) Toward Understanding the Diverse Roles of Perisomatic Interneurons in Epilepsy. Epilepsy Curr, 22, 54–60.

Eggan, S.M. & Lewis, D.A. (2007) Immunocytochemical distribution of the cannabinoid CB1 receptor in the primate neocortex: a regional and laminar analysis. Cereb Cortex, 17, 175–191.

Eggan, S.M., Melchitzky, D.S., Sesack, S.R., Fish, K.N. & Lewis, D.A. (2010) Relationship of cannabinoid CB1 receptor and cholecystokinin immunoreactivity in monkey dorsolateral prefrontal cortex. Neuroscience, 169, 1651–1661.

El Mestikawy, S., Wallen-Mackenzie, A., Fortin, G.M., Descarries, L. & Trudeau, L.E. (2011) From glutamate co-release to vesicular synergy: vesicular glutamate transporters. Nat Rev Neurosci, 12, 204–216.

Fan, P. (1995) Cannabinoid agonists inhibit the activation of 5-HT3 receptors in rat nodose ganglion neurons. J Neurophysiol, 73, 907–910.

Farrell, J.S., Colangeli, R., Dong, A., George, A.G., Addo-Osafo, K., Kingsley, P.J., Morena, M., Wolff, M.D., Dudok, B., He, K., Patrick, T.A., Sharkey, K.A., Patel, S., Marnett, L.J., Hill, M.N., Li, Y., Teskey, G.C. & Soltesz, I. (2021) In vivo endocannabinoid dynamics at the timescale of physiological and pathological neural activity. Neuron, 109, 2398–2403.e2394.

Fasano, C., Rocchetti, J., Pietrajtis, K., Zander, J.F., Manseau, F., Sakae, D.Y., Marcus-Sells, M., Ramet, L., Morel, L.J., Carrel, D., Dumas, S., Bolte, S., Bernard, V., Vigneault, E., Goutagny, R., Ahnert-Hilger, G., Giros, B., Daumas, S., Williams, S. & El Mestikawy, S. (2017) Regulation of the Hippocampal Network by VGLUT3-Positive CCK-GABAergic Basket Cells. Front Cell Neurosci, 11, 140.

Favier, M., Martin Garcia, E., Icick, R., de Almeida, C., Jehl, J., Desplanque, M., Zimmermann, J., Henrion, A., Mansouri-Guilani, N., Mounier, C., Ribeiro, S., Henderson, F., Geoffroy, A., Mella, S., Poirel, O., Bernard, V., Fabre, V., Li, Y., Rosenmund, C., Jamain, S., Vorspan, F., Mourot, A., Duriez, P., Pinhas, L., Maldonado, R., Pietrancosta, N., Daumas, S. & El Mestikawy, S. (2024) The human VGLUT3-pT8I mutation elicits uneven striatal DA signaling, food or drug maladaptive consumption in male mice. Nat Commun, 15, 5691.

Favier, M., Pietrancosta, N., El Mestikawy, S. & Gangarossa, G. (2021) Leveraging VGLUT3 Functions to Untangle Brain Dysfunctions. Trends Pharmacol Sci, 42, 475–490.

Ferraguti, F., Klausberger, T., Cobden, P., Baude, A., Roberts, J.D.B., Szucs, P., Kinoshita, A., Shigemoto, R., Somogyi, P. & Dalezios, Y. (2005) Metabotropic glutamate receptor 8-expressing nerve terminals target subsets of GABAergic neurons in the hippocampus. J Neurosci, 25, 10520–10536.

Field, M., Lukacs, I.P., Hunter, E., Stacey, R., Plaha, P., Livermore, L., Ansorge, O. & Somogyi, P. (2021) Tonic GABA_A_ receptor mediated currents of human cortical GABAergic interneurons vary amongst cell types. J Neurosci, 41, 9702–9719.

Földy, C., Malenka, R.C. & Südhof, T.C. (2013) Autism-associated neuroligin-3 mutations commonly disrupt tonic endocannabinoid signaling. Neuron, 78, 498–509.

Foldy, C., Neu, A., Jones, M.V. & Soltesz, I. (2006) Presynaptic, activity-dependent modulation of cannabinoid type 1 receptor-mediated inhibition of GABA release. J Neurosci, 26, 1465–1469.

Forro, T. & Klausberger, T. (2023) Differential behavior-related activity of distinct hippocampal interneuron types during odor-associated spatial navigation. Neuron, 111, 2399–2413.e2395.

Forte, N., Nicois, A., Marfella, B., Mavaro, I., D’Angelo, L., Piscitelli, F., Scandurra, A., De Girolamo, P., Baldelli, P., Benfenati, F., Di Marzo, V. & Cristino, L. (2024) Early endocannabinoid-mediated depolarization-induced suppression of excitation delays the appearance of the epileptic phenotype in synapsin II knockout mice. Cell Mol Life Sci, 81, 37.

Fremeau, J., R.T., Burman, J., Qureshi, T., Tran, C.H., Proctor, J., Johnson, J., Zhang, H., Sulzer, D., Copenhagen, D.R., Storm-Mathisen, J., Reimer, R.J., Chaudhry, F.A. & Edwards, R.H. (2002) The identification of vesicular glutamate transporter 3 suggests novel modes of signaling by glutamate Proc Nat Acad Sci USA, 99, 14488–14493.

Freund, T.F., Magloczky, Z., Soltesz, I. & Somogyi, P. (1986) Synaptic connections, axonal and dendritic patterns of neurons immunoreactive for cholecystokinin in the visual cortex of the cat. Neuroscience, 19, 1133–1159.

Freund, T.F., Martin, K.A.C., Soltesz, I., Somogyi, P. & Whitteridge, D. (1989) Arborisation pattern and postsynaptic targets of physiologically identified thalamocortical afferents in the monkey striate cortex. J Comp Neurol, 289, 315–336.

Fuentealba, P., Klausberger, T., Karayannis, T., Suen, W.Y., Huck, J., Tomioka, R., Rockland, K., Capogna, M., Studer, M., Morales, M. & Somogyi, P. (2010) Expression of COUP-TFII nuclear receptor in restricted GABAergic neuronal populations in the adult rat hippocampus. J Neurosci, 30, 1595–1609.

Fukaya, M., Uchigashima, M., Nomura, S., Hasegawa, Y., Kikuchi, H. & Watanabe, M. (2008) Predominant expression of phospholipase Cbeta1 in telencephalic principal neurons and cerebellar interneurons, and its close association with related signaling molecules in somatodendritic neuronal elements. Eur J Neurosci, 28, 1744–1759.

Fukudome, Y., Ohno-Shosaku, T., Matsui, M., Omori, Y., Fukaya, M., Tsubokawa, H., Taketo, M.M., Watanabe, M., Manabe, T. & Kano, M. (2004) Two distinct classes of muscarinic action on hippocampal inhibitory synapses: M2-mediated direct suppression and M1/M3-mediated indirect suppression through endocannabinoid signalling. Eur J Neurosci, 19, 2682–2692.

Gabbott, P.L. & Bacon, S.J. (1997) Vasoactive intestinal polypeptide containing neurones in monkey medial prefrontal cortex (mPFC): colocalisation with calretinin. Brain Res, 744, 179–184.

Glickfeld, L.L. & Scanziani, M. (2006) Distinct timing in the activity of cannabinoid-sensitive and cannabinoid-insensitive basket cells. Nat Neurosci, 9, 807–815.

Gonzalez-Albo, M.C., Elston, G.N. & DeFelipe, J. (2001) The human temporal cortex: characterization of neurons expressing nitric oxide synthase, neuropeptides and calcium-binding proteins, and their glutamate receptor subunit profiles. Cereb Cortex, 11, 1170–1181.

Gras, C., Herzog, E., Bellenchi, G.C., Bernard, V., Ravassard, P., Pohl, M., Gasnier, B., Giros, B. & ElMestikawy, S. (2002) A third vesicular glutamate transporter expressed by cholinergic and serotoninergic neurons. J Neurosci, 22, 5442–5451.

Gray, E.G. (1959) Axo-somatic and axo-dendritic synapses of the cerebral cortex: an electron microscope study. J Anat, 93, 420–433.

Grieco, S.F., Johnston, K.G., Gao, P., Garduño, B.M., Tang, B., Yi, E., Sun, Y., Horwitz, G.D., Yu, Z., Holmes, T.C. & Xu, X. (2023) Anatomical and molecular characterization of parvalbumin-cholecystokinin co-expressing inhibitory interneurons: implications for neuropsychiatric conditions. Mol Psychiatry, 28, 5293–5308.

Guo, B., Xi, K., Mao, H., Ren, K., Xiao, H., Hartley, N.D., Zhang, Y., Kang, J., Liu, Y., Xie, Y., Zhou, Y., Zhu, Y., Zhang, X., Fu, Z., Chen, J.F., Hu, H., Wang, W. & Wu, S. (2024) CB1R dysfunction of inhibitory synapses in the ACC drives chronic social isolation stress-induced social impairments in male mice. Neuron, 112, 441–457.e446.

Hajos, N., Katona, I., Naiem, S.S., Mackie, K., Ledent, C., Mody, I. & Freund, T.F. (2000) Cannabinoids inhibit hippocampal GABAergic transmission and network oscillations. Eur J Neurosci, 12, 3239–3249.

Hashimotodani, Y., Ohno-Shosaku, T., Tsubokawa, H., Ogata, H., Emoto, K., Maejima, T., Araishi, K., Shin, H.S. & Kano, M. (2005) Phospholipase Cbeta serves as a coincidence detector through its Ca2+ dependency for triggering retrograde endocannabinoid signal. Neuron, 45, 257–268.

Hefft, S. & Jonas, P. (2005) Asynchronous GABA release generates long-lasting inhibition at a hippocampal interneuron-principal neuron synapse. Nat Neurosci, 8, 1319–1328.

Henry, D.J. & Chavkin, C. (1995) Activation of inwardly rectifying potassium channels (GIRK1) by co-expressed rat brain cannabinoid receptors in Xenopus oocytes. Neurosci Lett, 186, 91–94.

Herzog, E., Gilchrist, J., Gras, C., Muzerelle, A., Ravassard, P., Giros, B., Gaspar, P. & El Mestikawy, S. (2004) Localization of VGLUT3, the vesicular glutamate transporter type 3, in the rat brain. Neuroscience, 123, 983–1002.

Hill, E.L., Gallopin, T., Ferezou, I., Cauli, B., Rossier, J., Schweitzer, P. & Lambolez, B. (2007) Functional CB1 receptors are broadly expressed in neocortical GABAergic and glutamatergic neurons. J Neurophysiol, 97, 2580–2589.

Hodge, R.D., Bakken, T.E., Miller, J.A., Smith, K.A., Barkan, E.R., Graybuck, L.T., Close, J.L., Long, B., Johansen, N., Penn, O., Yao, Z., Eggermont, J., Höllt, T., Levi, B.P., Shehata, S.I., Aevermann, B., Beller, A., Bertagnolli, D., Brouner, K., Casper, T., Cobbs, C., Dalley, R., Dee, N., Ding, S.L., Ellenbogen, R.G., Fong, O., Garren, E., Goldy, J., Gwinn, R.P., Hirschstein, D., Keene, C.D., Keshk, M., Ko, A.L., Lathia, K., Mahfouz, A., Maltzer, Z., McGraw, M., Nguyen, T.N., Nyhus, J., Ojemann, J.G., Oldre, A., Parry, S., Reynolds, S., Rimorin, C., Shapovalova, N.V., Somasundaram, S., Szafer, A., Thomsen, E.R., Tieu, M., Quon, G., Scheuermann, R.H., Yuste, R., Sunkin, S.M., Lelieveldt, B., Feng, D., Ng, L., Bernard, A., Hawrylycz, M., Phillips, J.W., Tasic, B., Zeng, H., Jones, A.R., Koch, C. & Lein, E.S. (2019) Conserved cell types with divergent features in human versus mouse cortex. Nature, 573, 61–68.

Hsieh, L.S. & Levine, E.S. (2013) Cannabinoid modulation of backpropagating action potential-induced calcium transients in layer 2/3 pyramidal neurons. Cereb Cortex, 23, 1731–1741.

Huang, C.C., Lo, S.W. & Hsu, K.S. (2001) Presynaptic mechanisms underlying cannabinoid inhibition of excitatory synaptic transmission in rat striatal neurons. J Physiol, 532, 731–748.

Hurd, Y.L., Manzoni, O.J., Pletnikov, M.V., Lee, F.S., Bhattacharyya, S. & Melis, M. (2019) Cannabis and the Developing Brain: Insights into Its Long-Lasting Effects. J Neurosci, 39, 8250–8258.

Ilyasov, A.A., Milligan, C.E., Pharr, E.P. & Howlett, A.C. (2018) The Endocannabinoid System and Oligodendrocytes in Health and Disease. Front Neurosci, 12, 733.

Iversen, L. (2003) Cannabis and the brain. Brain, 126, 1252–1270.

Katona, I., Sperlagh, B., Magloczky, Z., Santha, E., Kofalvi, A., Czirjak, S., Mackie, K., Vizi, E.S. & Freund, T.F. (2000) GABAergic interneurons are the targets of cannabinoid actions in the human hippocampus. Neuroscience, 100, 797–804.

Katona, I., Sperlagh, B., Sik, A., Kafalvi, A., Vizi, E.S., Mackie, K. & Freund, T.F. (1999) Presynaptically located CB1 cannabinoid receptors regulate GABA release from axon terminals of specific hippocampal interneurons. J Neurosci, 19, 4544–4558.

Katona, I., Urbán, G.M., Wallace, M., Ledent, C., Jung, K.M., Piomelli, D., Mackie, K. & Freund, T.F. (2006) Molecular composition of the endocannabinoid system at glutamatergic synapses. J Neurosci, 26, 5628–5637.

Kawaguchi, Y. & Kubota, Y. (1998) Neurochemical features and synaptic connections of large physiologically-identified GABAergic cells in the rat frontal cortex. Neuroscience, 85, 677–701.

Kawamura, Y., Fukaya, M., Maejima, T., Yoshida, T., Miura, E., Watanabe, M., Ohno-Shosaku, T. & Kano, M. (2006) The CB1 cannabinoid receptor is the major cannabinoid receptor at excitatory presynaptic sites in the hippocampus and cerebellum. J Neurosci, 26, 2991–3001.

Kim, H.H., Park, J.M., Lee, S.H. & Ho, W.K. (2019) Association of mGluR-Dependent LTD of Excitatory Synapses with Endocannabinoid-Dependent LTD of Inhibitory Synapses Leads to EPSP to Spike Potentiation in CA1 Pyramidal Neurons. J Neurosci, 39, 224–237.

Kim, J., Isokawa, M., Ledent, C. & Alger, B.E. (2002) Activation of muscarinic acetylcholine receptors enhances the release of endogenous cannabinoids in the hippocampus. J Neurosci, 22, 10182–10191.

Kisvarday, Z.F., Adams, C.B.T. & Smith, A.D. (1985) Axo-axonic (chandelier) cells in human epileptic temporal cortex: afferent and efferent synapses. Neurosci Letts, 22, S87.

Kisvarday, Z.F., Gulyas, A., Beroukas, D., North, J.B., Chubb, I.W. & Somogyi, P. (1990) Synapses, axonal and dendritic patterns of GABA-immunoreactive neurons in human cerebral cortex. Brain, 113, 793–812.

Klausberger, T., Marton, L.F., O’Neill, J., Huck, J.H.J., Dalezios, Y., Fuentealba, P., Suen, W.Y., Papp, E., Kaneko, T., Watanabe, M., Csicsvari, J. & Somogyi, P. (2005) Complementary roles of cholecystokinin- and parvalbumin-expressing GABAergic neurons in hippocampal network oscillations. J Neurosci, 25, 9782–9793.

Koukouli, F., Montmerle, M., Aguirre, A., De Brito Van Velze, M., Peixoto, J., Choudhary, V., Varilh, M., Julio-Kalajzic, F., Allene, C., Mendéz, P., Zerlaut, Y., Marsicano, G., Schlüter, O.M., Rebola, N., Bacci, A. & Lourenço, J. (2022) Visual-area-specific tonic modulation of GABA release by endocannabinoids sets the activity and coordination of neocortical principal neurons. Cell Rep, 40, 111202.

Kovacs, F.E., Knop, T., Urbanski, M.J., Freiman, I., Freiman, T.M., Feuerstein, T.J., Zentner, J. & Szabo, B. (2012) Exogenous and endogenous cannabinoids suppress inhibitory neurotransmission in the human neocortex. Neuropsychopharmacology, 37, 1104–1114.

Kreitzer, A.C. & Regehr, W.G. (2001) Retrograde inhibition of presynaptic calcium influx by endogenous cannabinoids at excitatory synapses onto Purkinje cells. Neuron, 29, 717–727.

Krienen, F.M., Goldman, M., Zhang, Q., R, C.H.D.R., Florio, M., Machold, R., Saunders, A., Levandowski, K., Zaniewski, H., Schuman, B., Wu, C., Lutservitz, A., Mullally, C.D., Reed, N., Bien, E., Bortolin, L., Fernandez-Otero, M., Lin, J.D., Wysoker, A., Nemesh, J., Kulp, D., Burns, M., Tkachev, V., Smith, R., Walsh, C.A., Dimidschstein, J., Rudy, B., L, S.K., Berretta, S., Fishell, G., Feng, G. & McCarroll, S.A. (2020) Innovations present in the primate interneuron repertoire. Nature, 586, 262–269.

Kubota, Y. & Kawaguchi, Y. (1997) Two distinct subgroups of cholycystokinin-immunoreactive cortical interneurons Brain Res, 752, 175–183.

Larkum, M.E., Zhu, J.J. & Sakmann, B. (1999) A new cellular mechanism for coupling inputs arriving at different cortical layers. Nature, 398, 338–341.

Lee, B.R. & Dalley, R. & Miller, J.A. & Chartrand, T. & Close, J. & Mann, R. & Mukora, A. & Ng, L. & Alfiler, L. & Baker, K. & Bertagnolli, D. & Brouner, K. & Casper, T. & Csajbok, E. & Donadio, N. & Driessens, S.L.W. & Egdorf, T. & Enstrom, R. & Galakhova, A.A. & Gary, A. & Gelfand, E. & Goldy, J. & Hadley, K. & Heistek, T.S. & Hill, D. & Hou, W.H. & Johansen, N. & Jorstad, N. & Kim, L. & Kocsis, A.K. & Kruse, L. & Kunst, M. & León, G. & Long, B. & Mallory, M. & Maxwell, M. & McGraw, M. & McMillen, D. & Melief, E.J. & Molnar, G. & Mortrud, M.T. & Newman, D. & Nyhus, J. & Opitz-Araya, X. & Ozsvár, A. & Pham, T. & Pom, A. & Potekhina, L. & Rajanbabu, R. & Ruiz, A. & Sunkin, S.M. & Szöts, I. & Taskin, N. & Thyagarajan, B. & Tieu, M. & Trinh, J. & Vargas, S. & Vumbaco, D. & Waleboer, F. & Walling-Bell, S. & Weed, N. & Williams, G. & Wilson, J. & Yao, S. & Zhou, T. & Barzó, P. & Bakken, T. & Cobbs, C. & Dee, N. & Ellenbogen, R.G. & Esposito, L. & Ferreira, M. & Gouwens, N.W. & Grannan, B. & Gwinn, R.P. & Hauptman, J.S. & Hodge, R. & Jarsky, T. & Keene, C.D. & Ko, A.L. & Korshoej, A.R. & Levi, B.P. & Meier, K. & Ojemann, J.G. & Patel, A. & Ruzevick, J. & Silbergeld, D.L. & Smith, K. & Sørensen, J.C. & Waters, J. & Zeng, H. & Berg, J. & Capogna, M. & Goriounova, N.A. & Kalmbach, B. & de Kock, C.P.J. & Mansvelder, H.D. & Sorensen, S.A. & Tamas, G. & Lein, E.S. & Ting, J.T. (2023) Signature morphoelectric properties of diverse GABAergic interneurons in the human neocortex. Science, 382, eadf6484.

Lee, C.T., Li, L., Takamoto, N., Martin, J.F., Demayo, F.J., Tsai, M.J. & Tsai, S.Y. (2004) The nuclear orphan receptor COUP-TFII is required for limb and skeletal muscle development. Mol Cell Biol, 24, 10835–10843.

Lee, S.H., Foldy, C. & Soltesz, I. (2010) Distinct endocannabinoid control of GABA release at perisomatic and dendritic synapses in the hippocampus. J Neurosci, 30, 7993–8000.

Lee, S.H., Ledri, M., Tóth, B., Marchionni, I., Henstridge, C.M., Dudok, B., Kenesei, K., Barna, L., Szabó, S.I., Renkecz, T., Oberoi, M., Watanabe, M., Limoli, C.L., Horvai, G., Soltesz, I. & Katona, I. (2015) Multiple Forms of Endocannabinoid and Endovanilloid Signaling Regulate the Tonic Control of GABA Release. J Neurosci, 35, 10039–10057.

Lee, S.H. & Soltesz, I. (2011) Requirement for CB1 but not GABA_B_ receptors in the cholecystokinin mediated inhibition of GABA release from cholecystokinin expressing basket cells. J Physiol, 589, 891–902.

Lenkey, N., Kirizs, T., Holderith, N., Mate, Z., Szabo, G., Vizi, E.S., Hajos, N. & Nusser, Z. (2015) Tonic endocannabinoid-mediated modulation of GABA release is independent of the CB1 content of axon terminals. Nat Commun, 6, 6557.

Losonczy, A., Biro, A.A. & Nusser, Z. (2004) Persistently active cannabinoid receptors mute a subpopulation of hippocampal interneurons. Proc Nat Acad Sci USA, 101, 1362–1367.

Losonczy, A. & Magee, J.C. (2006) Integrative properties of radial oblique dendrites in hippocampal CA1 pyramidal neurons. Neuron, 50, 291–307.

Lourenço, J. & Bacci, A. (2017) Human-Specific Cortical Synaptic Connections and Their Plasticity: Is That What Makes Us Human? PLoS Biol, 15, e2001378.

Lovett-Barron, M., Turi, G.F., Kaifosh, P., Lee, P.H., Bolze, F., Sun, X.H., Nicoud, J.F., Zemelman, B.V., Sternson, S.M. & Losonczy, A. (2012) Regulation of neuronal input transformations by tunable dendritic inhibition. Nat Neurosci, 15, 423–430, s421-423.

Lowe, H., Toyang, N., Steele, B., Bryant, J. & Ngwa, W. (2021) The Endocannabinoid System: A Potential Target for the Treatment of Various Diseases. Int J Mol Sci, 22.

Ludanyi, A., Hu, S.S., Yamazaki, M., Tanimura, A., Piomelli, D., Watanabe, M., Kano, M., Sakimura, K., Magloczky, Z., Mackie, K., Freund, T.F. & Katona, I. (2011) Complementary synaptic distribution of enzymes responsible for synthesis and inactivation of the endocannabinoid 2-arachidonoylglycerol in the human hippocampus. Neuroscience, 174, 50–63.

Lujan, R., Nusser, Z., Roberts, J.D.B., Shigemoto, R. & Somogyi, P. (1996) Perisynaptic location of metabotropic glutamate receptors mGluR1 and mGluR5 on dendrites and dendritic spines in the rat hippocampus. Eur. J. Neurosci., 8, 1488–1500.

Lukacs, I.P., Francavilla, R., Field, M., Hunter, E., Howarth, M., Horie, S., Plaha, P., Stacey, R., Livermore, L., Ansorge, O., Tamas, G. & Somogyi, P. (2023) Differential effects of group III metabotropic glutamate receptors on spontaneous inhibitory synaptic currents in spine-innervating double bouquet and parvalbumin-expressing dendrite-targeting GABAergic interneurons in human neocortex. Cerebral Cortex, 33, 2101–2142.

Lutz, B., Marsicano, G., Maldonado, R. & Hillard, C.J. (2015) The endocannabinoid system in guarding against fear, anxiety and stress. Nat Rev Neurosci, 16, 705–718.

Marinelli, S., Pacioni, S., Bisogno, T., Di Marzo, V., Prince, D.A., Huguenard, J.R. & Bacci, A. (2008) The endocannabinoid 2-arachidonoylglycerol is responsible for the slow self-inhibition in neocortical interneurons. J Neurosci, 28, 13532–13541.

Marsicano, G. & Lutz, B. (1999) Expression of the cannabinoid receptor CB1 in distinct neuronal subpopulations in the adult mouse forebrain. Eur J Neurosci, 11, 4213–4225.

Mato, S., Chevaleyre, V., Robbe, D., Pazos, A., Castillo, P.E. & Manzoni, O.J. (2004) A single in-vivo exposure to delta 9THC blocks endocannabinoid-mediated synaptic plasticity. Nat Neurosci, 7, 585–586.

Meskenaite, V. (1997) Calretinin-immunoreactive local circuit neurons in area 17 of the cynomolgus monkey, Macaca fascicularis. J Comp Neurol, 379, 113–132.

Morales, P. & Reggio, P.H. (2017) An Update on Non-CB(1), Non-CB(2) Cannabinoid Related G-Protein-Coupled Receptors. Cannabis Cannabinoid Res, 2, 265–273.

Musella, A. & Centonze, D. (2023) Electrophysiology of Endocannabinoid Signaling. Methods Mol Biol, 2576, 461–475.

Nagy-Pál, P., Veres, J.M., Fekete, Z., Karlócai, M.R., Weisz, F., Barabás, B., Reéb, Z. & Hájos, N. (2023) Structural Organization of Perisomatic Inhibition in the Mouse Medial Prefrontal Cortex. J Neurosci, 43, 6972–6987.

Nakagawa, S., Watanabe, M., Isobe, T., Kondo, H. & Inoue, Y. (1998) Cytological compartmentalization in the staggerer cerebellum, as revealed by calbindin immunohistochemistry for Purkinje cells. J Comp Neurol, 395, 112–120.

Navarrete, M., Díez, A. & Araque, A. (2014) Astrocytes in endocannabinoid signalling. Philos Trans R Soc Lond B Biol Sci, 369, 20130599.

Navarro, D., Gasparyan, A., Navarrete, F., Torregrosa, A.B., Rubio, G., Marín-Mayor, M., Acosta, G.B., Garcia-Gutiérrez, M.S. & Manzanares, J. (2022) Molecular Alterations of the Endocannabinoid System in Psychiatric Disorders. Int J Mol Sci, 23.

Neu, A., Földy, C. & Soltesz, I. (2007) Postsynaptic origin of CB1-dependent tonic inhibition of GABA release at cholecystokinin-positive basket cell to pyramidal cell synapses in the CA1 region of the rat hippocampus. J Physiol, 578, 233–247.

Nunzi, M.G., Gorio, A., Milan, F., Freund, T.F., Somogyi, P. & Smith, A.D. (1985) Cholecystokinin-immunoreactive cells form symmetrical synaptic contacts with pyramidal and non-pyramidal neurons in the hippocampus. J Comp Neurol, 237, 485–505.

Nyíri, G., Cserép, C., Szabadits, E., MacKie, K. & Freund, T.F. (2005) CB1 cannabinoid receptors are enriched in the perisynaptic annulus and on preterminal segments of hippocampal GABAergic axons. Neuroscience, 136, 811–822.

Nyiri, G., Szabadits, E., Cserep, C., Mackie, K., Shigemoto, R. & Freund, T.F. (2005) GABA_B_ and CB_1_ cannabinoid receptor expression identifies two types of septal cholinergic neurons. Eur J Neurosci, 21, 3034–3042.

Ohno-Shosaku, T., Maejima, T. & Kano, M. (2001) Endogenous cannabinoids mediate retrograde signals from depolarized postsynaptic neurons to presynaptic terminals. Neuron, 29, 729–738.

Ohno-Shosaku, T., Shosaku, J., Tsubokawa, H. & Kano, M. (2002) Cooperative endocannabinoid production by neuronal depolarization and group I metabotropic glutamate receptor activation. Eur J Neurosci, 15, 953–961.

Olah, S., Komlosi, G., Szabadics, J., Varga, C., Toth, E., Barzo, P. & Tamas, G. (2007) Output of neurogliaform cells to various neuron types in the human and rat cerebral cortex. Front Neural Circuits, 1, 4.

Omiya, Y., Uchigashima, M., Konno, K., Yamasaki, M., Miyazaki, T., Yoshida, T., Kusumi, I. & Watanabe, M. (2015) VGluT3-expressing CCK-positive basket cells construct invaginating synapses enriched with endocannabinoid signaling proteins in particular cortical and cortex-like amygdaloid regions of mouse brains. J Neurosci, 35, 4215–4228.

Pelkey, K.A., Calvigioni, D., Fang, C., Vargish, G., Ekins, T., Auville, K., Wester, J.C., Lai, M., Mackenzie-Gray Scott, C., Yuan, X., Hunt, S., Abebe, D., Xu, Q., Dimidschstein, J., Fishell, G., Chittajallu, R. & McBain, C.J. (2020) Paradoxical network excitation by glutamate release from VGluT3(+) GABAergic interneurons. Elife, 9.

Piomelli, D. & Mabou Tagne, A. (2022) Endocannabinoid-Based Therapies. Annu Rev Pharmacol Toxicol, 62, 483–507.

Pitler, T.A. & Alger, B.E. (1994) Depolarization-induced suppression of GABAergic inhibition in rat hippocampal pyramidal cells: G protein involvement in a presynaptic mechanism. Neuron, 13, 1447–1455.

Schiffmann, S.N., Cheron, G., Lohof, A., d’Alcantara, P., Meyer, M., Parmentier, M. & Schurmans, S. (1999) Impaired motor coordination and Purkinje cell excitability in mice lacking calretinin. Proc Nat Acad Sci, 96, 5257–5262.

Schlag, A.K., O’Sullivan, S.E., Zafar, R.R. & Nutt, D.J. (2021) Current controversies in medical cannabis: Recent developments in human clinical applications and potential therapeutics. Neuropharmacology, 191, 108586.

Schmitz, S.K., King, C., Kortleven, C., Huson, V., Kroon, T., Kevenaar, J.T., Schut, D., Saarloos, I., Hoetjes, J.P., de Wit, H., Stiedl, O., Spijker, S., Li, K.W., Mansvelder, H.D., Smit, A.B., Cornelisse, L.N., Verhage, M. & Toonen, R.F. (2016) Presynaptic inhibition upon CB1 or mGlu2/3 receptor activation requires ERK/MAPK phosphorylation of Munc18-1. Embo j, 35, 1236–1250.

Schwaller, B., Dick, J., Dhoot, G., Carroll, S., Vrbova, G., Nicotera, P., Pette, D., Wyss, A., Bluethmann, H., Hunziker, W. & Celio, M.R. (1999) Prolonged contraction-relaxation cycle of fast-twitch muscles in parvalbumin knockout mice. American Journal of Physiology - Cell Physiology, 276, C395–C403.

Shi, Y., An, J., Liang, J., Hayes, S.E., Sandusky, G.E., Stramm, L.E. & Yang, N.N. (1999) Characterization of a Mutant Pancreatic eIF-2&#x3b1; Kinase, PEK, and Co-localization with Somatostatin in Islet Delta Cells. Journal of Biological Chemistry, 274, 5723–5730.

Shigemoto, R., Kulik, A., Roberts, J.D.B., Ohishi, H., Nusser, Z., Kaneko, T. & Somogyi, P. (1996) Target-cell-specific concentration of a metabotropic glutamate receptor in the presynaptic active zone. Nature, 381, 523–525.

Sigel, E., Baur, R., Rácz, I., Marazzi, J., Smart, T.G., Zimmer, A. & Gertsch, J. (2011) The major central endocannabinoid directly acts at GABA(A) receptors. Proc Natl Acad Sci U S A, 108, 18150–18155.

Soltesz, I. & Staley, K. (2006) High times for memory: cannabis disrupts temporal coordination among hippocampal neurons. Nat Neurosci, 9, 1461–1463.

Somogyi, J., Baude, A., Omori, Y., Shimizu, H., El Mestikawy, S., Fukaya, M., Shigemoto, R., Watanabe, M. & Somogyi, P. (2004) GABAergic basket cells expressing cholecystokinin contain vesicular glutamate transporter type 3 (VGLUT3) in their synaptic terminals in hippocampus and isocortex of the rat. Eur J Neurosci, 19, 552–569.

Somogyi, P. (1978) The study of Golgi stained cells and of experimental degeneration under the electron microscope : a direct method for the identification in the visual cortex of three successive links in a neuron chain. Neuroscience, 3, 167–180.

Somogyi, P. & Cowey, A. (1981) Combined Golgi and electron microscopic study on the synapses formed by double bouquet cells in the visual cortex of the cat and monkey. J Comp Neurol, 195, 547–566.

Somogyi, P., Dalezios, Y., Luján, R., Roberts, J.D.B., Watanabe, M. & Shigemoto, R. (2003) High level of mGluR7 in the presynaptic active zones of select populations of GABAergic terminals innervating interneurones in the rat hippocampus. Eur J Neurosci, 17, 2503–2520.

Somogyi, P., Freund, T.F. & Cowey, A. (1982) The axo-axonic interneuron in the cerebral cortex of the rat, cat and monkey. Neuroscience, 7, 2577–2607.

Somogyi, P., Kisvarday, Z.F., Martin, K.A.C. & Whitteridge, D. (1983) Synaptic connections of morphologically identified and physiologically characterized large basket cells in the striate cortex of cat. Neuroscience, 10, 261–294.

Somogyi, P. & Klausberger, T. (2018) Hippocampus: Intrinsic organisation. In Shepherd, G.M., Grillner, S. (eds) Handbook of Brain Microcircuits, 2nd Edition. Oxford University Press, Oxford, pp. 624.

Somogyi, P., Tamas, G., Lujan, R. & Buhl, E.H. (1998) Salient features of synaptic organisation in the cerebral cortex. Brain Res Rev, 26, 113–135.

Sugaya, Y. & Kano, M. (2021) Endocannabinoid-Mediated Control of Neural Circuit Excitability and Epileptic Seizures. Front Neural Circuits, 15, 781113.

Szabadits, E., Cserep, C., Szonyi, A., Fukazawa, Y., Shigemoto, R., Watanabe, M., Itohara, S., Freund, T.F. & Nyiri, G. (2011) NMDA receptors in hippocampal GABAergic synapses and their role in nitric oxide signaling. J Neurosci, 31, 5893–9504.

Szegedi, V., Paizs, M., Baka, J., Barzó, P., Molnár, G., Tamas, G. & Lamsa, K. (2020) Robust perisomatic GABAergic self-innervation inhibits basket cells in the human and mouse supragranular neocortex. Elife, 9.

Szegedi, V., Paizs, M., Csakvari, E., Molnar, G., Barzo, P., Tamas, G. & Lamsa, K. (2016) Plasticity in Single Axon Glutamatergic Connection to GABAergic Interneurons Regulates Complex Events in the Human Neocortex. PLoS Biol, 14, e2000237.

Szentagothai, J. (1978) The neuron network of the cerebral cortex: a functional interpretation. The Ferrier Lecture, 1977. Proc R Soc Lond B, 201, 219–248.

Szonyi, A., Mayer, M.I., Cserep, C., Takacs, V.T., Watanabe, M., Freund, T.F. & Nyiri, G. (2016) The ascending median raphe projections are mainly glutamatergic in the mouse forebrain. Brain Struct Funct, 221, 735–751.

Tamas, G., Buhl, E.H. & Somogyi, P. (1997) Fast IPSPs elicited via multiple synaptic release sites by distinct types of GABAergic neurone in the cat visual cortex. J Physiol (Lond), 500, 715–738.

Tamas, G., Lörincz, A., Simon, A. & Szabadics, S. (2003) Identified sources and targets of slow inhibition in the neocortex. Science, 299, 1902–1905.

Tanimura, A., Yamazaki, M., Hashimotodani, Y., Uchigashima, M., Kawata, S., Abe, M., Kita, Y., Hashimoto, K., Shimizu, T., Watanabe, M., Sakimura, K. & Kano, M. (2010) The endocannabinoid 2-arachidonoylglycerol produced by diacylglycerol lipase alpha mediates retrograde suppression of synaptic transmission. Neuron, 65, 320–327.

Trettel, J. & Levine, E.S. (2002) Cannabinoids depress inhibitory synaptic inputs received by layer 2/3 pyramidal neurons of the neocortex. J Neurophysiol, 88, 534–539.

Tsou, K., Mackie, K., SanudoPena, M.C. & Walker, J.M. (1999) Cannabinoid CB1 receptors are localized primarily on cholecystokinin-containing gabaergic interneurons in the rat hippocampal formation. Neuroscience, 93, 969–975.

Varga, C., Lee, S.Y. & Soltesz, I. (2010) Target-selective GABAergic control of entorhinal cortex output. Nat Neurosci, 13, 822–824.

Varga, C., Tamas, G., Barzo, P., Olah, S. & Somogyi, P. (2015) Molecular and electrophysiological characterization of GABAergic interneurons expressing the transcription factor COUP-TFII in the adult human temporal cortex. Cereb. Cortex, 25, 4430–4449.

Varga, V., Losonczy, A., Zemelman, B.V., Borhegyi, Z., Nyiri, G., Domonkos, A., Hangya, B., Holderith, N., Magee, J.C. & Freund, T.F. (2009) Fast synaptic subcortical control of hippocampal circuits. Science, 326, 449–453.

Vicente-Acosta, A., Ceprian, M., Sobrino, P., Pazos, M.R. & Loría, F. (2022) Cannabinoids as Glial Cell Modulators in Ischemic Stroke: Implications for Neuroprotection. Front Pharmacol, 13, 888222.

Vigneault, E., Poirel, O., Riad, M., Prud’homme, J., Dumas, S., Turecki, G., Fasano, C., Mechawar, N. & El Mestikawy, S. (2015) Distribution of vesicular glutamate transporters in the human brain. Front Neuroanat, 9, 23.

Walter, L., Dinh, T. & Stella, N. (2004) ATP induces a rapid and pronounced increase in 2-arachidonoylglycerol production by astrocytes, a response limited by monoacylglycerol lipase. J Neurosci, 24, 8068–8074.

White, E.L. (1979) Thalamocortical synaptic relations: a review with emphasis on the projections of specific thalamic nuclei to the primary sensory areas of the neocortex. Brain Res Rev, 1, 275–311.

Wilson, R.I., Kunos, G. & Nicoll, R.A. (2001) Presynaptic specificity of endocannabinoid signaling in the hippocampus. Neuron, 31, 453–462.

Wilson, R.I. & Nicoll, R.A. (2001) Endogenous cannabinoids mediate retrograde signalling at hippocampal synapses. Nature, 410, 588–592.

Wong, H.C., Sternini, C., Lloyd, K., De Giorgio, R. & Walsh, J.H. (1996) Monoclonal antibody to VIP: production, characterization, immunoneutralizing activity, and usefulness in cytochemical staining. Hybridoma, 15, 133–139.

Yoshida, T., Hashimoto, K., Zimmer, A., Maejima, T., Araishi, K. & Kano, M. (2002) The cannabinoid CB1 receptor mediates retrograde signals for depolarization-induced suppression of inhibition in cerebellar Purkinje cells. J Neurosci, 22, 1690–1697.

